# Striosomes Target Nigral Dopamine-Containing Neurons via Direct-D1 and Indirect-D2 Pathways Paralleling Classic Direct-Indirect Basal Ganglia Systems

**DOI:** 10.1101/2024.06.01.596922

**Authors:** Iakovos Lazaridis, Jill R. Crittenden, Gun Ahn, Kojiro Hirokane, Tomoko Yoshida, Ian R. Wickersham, Ara Mahar, Vasiliki Skara, Johnny H. Loftus, Krishna Parvataneni, Konstantinos Meletis, Jonathan T. Ting, Emily Hueske, Ayano Matsushima, Ann M. Graybiel

**Affiliations:** McGovern Institute for Brain Research and Department of Brain and Cognitive Sciences; Karolinska Institutet, Stockholm, SE; Human Cell Types Dept, Allen Institute for Brain Science, Seattle WA 98109, USA; Department of Physiology and Biophysics, University of Washington, Seattle WA 98195, USA

**Keywords:** Striosomes, direct-D1 pathway, indirect-D2 pathway, dopamine control, striatal dopamine release, movement modulation, striatal projection neurons (SPNs), central zone of globus pallidus external (cGPe), substantia nigra pars compacta (SNpc), substantia nigra pars reticulata (SNpr)

## Abstract

Balanced activity of canonical direct D1 and indirect D2 basal ganglia pathways is considered a core requirement for normal movement, and their imbalance is an etiologic factor in movement and neuropsychiatric disorders. We present evidence for a conceptually equivalent pair of direct-D1 and indirect-D2 pathways that arise from striatal projection neurons (SPNs) of the striosome compartment rather than from SPNs of the matrix, as do the canonical pathways. These S-D1 and S-D2 striosomal pathways target substantia nigra dopamine-containing neurons instead of basal ganglia motor output nuclei. They modulate movement oppositely to the modulation by the canonical pathways: S-D1 is inhibitory and S-D2 is excitatory. The S-D1 and S-D2 circuits likely influence motivation for learning and action, complementing and reorienting canonical pathway modulation. A major conceptual reformulation of the classic direct-indirect pathway model of basal ganglia function is needed, as well as reconsideration of the effects of D2-targeting therapeutic drugs.

**HIGHLIGHTS:**

- Direct S-D1 and Indirect S-D2 striosomal pathways target SNpc dopamine cells
- The S-D2 indirect pathway targets a distinct central external pallidal zone (cGPe)
- Stimulation of S-D2 increases, of S-D1 decreases, striatal dopamine and movement
- S-D1 SPNs activity brackets task, inverse to a mid-task peak of dopamine release

## INTRODUCTION

The striatum, the largest nucleus of the basal ganglia, powerfully modulates movement and mood under normal conditions and enables reinforcement-driven plasticity to shape behavior. Massive loss of striatal dopamine occurs in Parkinson’s disease, and massive degeneration of striatal projection neurons (SPNs) occurs in Huntington’s disease. Thus, the striatum and its output pathways are a central focus in attempts to combat these and related basal ganglia disorders, and increasingly to combat neuropsychiatric conditions. Under normal conditions, the neural modulation of mood and movement by the basal ganglia is thought to be the result of a push-pull control system deploying opposing yet coordinated interactions between two canonical basal ganglia pathways originating in the striatal SPNs^1–8^. These classic circuits, the so-called direct and indirect pathways, target basal ganglia output nuclei either monosynaptically (the direct pathway) or via intermediary connectivity (the indirect pathway). In the simplest formulation of the direct-indirect model, dopamine increases direct pathway “Go” activity via D1 dopamine receptors expressed on direct-pathway SPNs in the striatal matrix and decreases “No-Go” activity of the indirect pathway via D2 dopamine receptors on indirect-pathway SPNs in the matrix. These actions of the receptors are referable to their opposite regulation of intracellular pathways^9–11^. As a result, the circuits can favor or impede intended movements, including promoting one intended movement sequence while inhibiting others^6,12–15^. In the classic model, the direct-D1 and indirect-D2 pathways arise from different ensembles of SPNs, target different output nuclei (pallidal segments and substantia nigra), and in turn differentially influence effector circuits that ultimately modulate corticothalamic, brainstem and spinal processing for movement control. Important amendments to this model allow for cooperative direct-indirect pathway operation, and for extensive collateral pathway crosstalk and other neuromodulatory and oscillatory effects^8,16–22^; but the direct-indirect model has been the cornerstone of clinical and basic science fields alike for nearly half a century.

An outlier in this model has been the strong output of some SPNs to the dopamine-containing neurons of the substantia nigra pars compacta (SNpc)—the very origin of the nigrostriatal tract that severely degenerates in Parkinson’s disease. These SPNs mainly, though not exclusively, lie in the so-called striosomes of the striatum. Striosomes are neurochemically distinct macroscopic compartments of the striatum distributed through much of the large extra-striosomal matrix from which the classical direct-indirect pathways arise. Despite the potential power of this striatonigral feedback circuit as a regulator of dopamine neuron activity, and despite the prominence of striosomes in the human striatum, as originally demonstrated^23^, the striosomal pathway has never been integrated into the canonical view of the direct and indirect pathways other than that it arises from D1-expressing SPNs. Nor has it yet been fully characterized.

Here we demonstrate that striosomes not only give rise to a direct D1 pathway to the SNpc, but also to an indirect pathway originating in D2-expressing striosomal SPNs that target a central, commonly overlooked subdivision of the external pallidum (GPe) that itself innervates nigral dopamine-containing neurons. This discovery suggests that the classic motor output pathways of the basal ganglia have twins emerging from striosomes that can affect the activity of the dopamine-containing nigral neurons that, in turn, modulate mood and movement through their actions on intraneuronal circuits, on inputs from the neocortex and striatum, and on circuits exiting from the SPNs themselves. We engineered and used multiple transgenic mouse lines, employed rabies viral tracing, optogenetics and novel striosome- and matrix-preferring ATAC-seq-based enhancers in experiments to examine these pathways.

We present here first anatomical evidence, and then functional evidence, converging on the view that dual direct-indirect pathways control voluntary behavioral action. We demonstrate an anatomical double dissociation between the striosomal direct (S-D1) and indirect (S-D2) pathways and the classic direct (M-D1) and indirect (M-D2) pathways of the matrix (M). By simultaneous monitoring of ensemble striatal activity and dopamine release, and by selective optogenetic manipulation of striosomal D1 and D2 SPNs in engineered mice, we then demonstrate that these identified S-D1 and S-D2 pathways are also functionally dissociable, and at least opponent both in their effects on dopamine release in the striatum and on their encoding of behaviors.

These effects of the S-D1 and S-D2 circuits on striatal dopamine release and locomotor behavior appear to be the reverse of those known for the classic direct D1 and indirectly D2 pathways insofar as we have tested for them, with the S-D2 circuit promoting locomotion and the S-D1 pathway reducing such movements. These findings call for a deep revision of the classic model of basal ganglia function, wherein these and other effects of the striosomal circuits can be incorporated.

Our findings suggest a parallel between basal ganglia pathways that share the motif of direct and indirect output components, yet radically differ in their targets and potential effects on behavior, with the classic pathways targeting motor pallidal and nigral output nuclei of the basal ganglia, and the striosomal pathways targeting the nigral dopamine system. We suggest that the fundamental parallelism of the direct and indirect striosomal and matrix circuits is likely to underlie equally fundamental differences in their parallel functions in motivation and movement control across states of health and illness.

## RESULTS

### Identification of striosomal output pathways to the pallidum and substantia nigra by the use of engineered mouse lines

Both D1 and D2 receptor-expressing SPNs lie in striosomes, with different gradients of topographic expression to be present in striosomes^42^, and our baseline counts indicated roughly equal proportions of D1 and D2 SPNs in the striosomes (S-D1 and S-D2 SPNs) (**Figures S1A-S1C**). We searched for mouse lines that could selectively label either the S-D1 or S-D2 striosomal populations.

We identified four lines of mice with striosome-enriched fluorophore expression, and one line with matrix-enriched fluorophore expression (**Figures S1 and S2, Table S1)**. We verified compartmentally biased labeling in three of the lines: striosomal MOR1-mCherry and P172-mCitrine, and matrix CalDAG-GEFI-GFP^42–45^. Two additional lines had striosome enrichment: AT1-tdTomato lines (14 and F), a newly generated transgenic line carrying a partial BAC for AT1 (angiotensin II receptor type 1) and as well a prepronociceptin-Cre line (Pnoc-Cre) developed by Bruchas and colleagues^46^ ^158^ **(Figures 1A, 1C and 1E**). Highly enriched striosomal expression occurred in a neurotensin-Cre line (Nts-Cre) crossed with a Cre-dependent tdTomato reporter line (Ai14)^47,48^ (**Figures 1B, 1D and 1E**).

**Figure 1.**
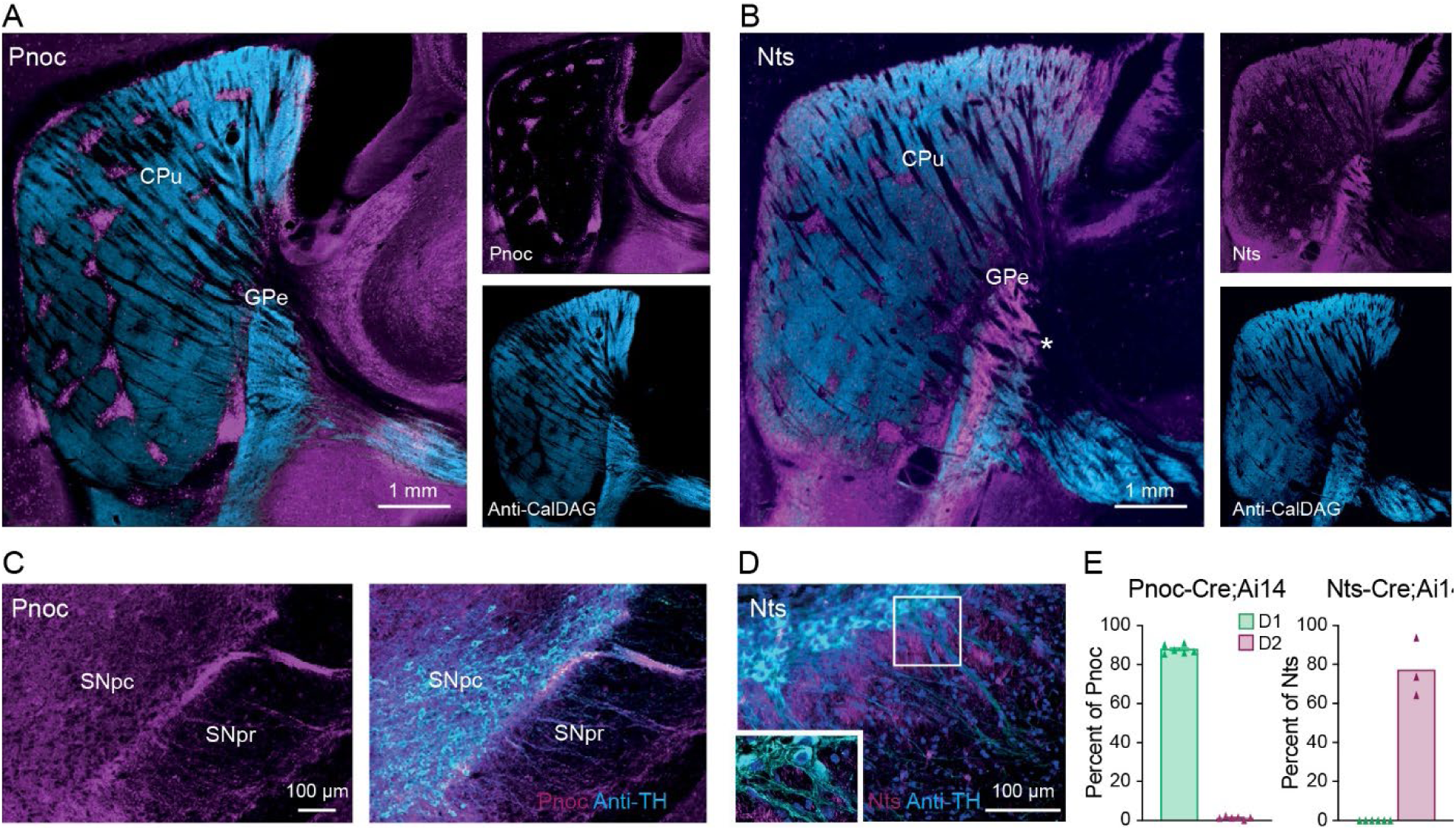
Striosomal S-D1 and S-D2 populations form parallel pathways to dopamine-containing nigral neurons. A. Sagittal section from Pnoc-Cre;Ai14-tdTomato (magenta) transgenic mouse brain immunolabeled for CalDAG-GEFI (cyan). CPu: caudoputamen. B. Section from Nts-Cre;Ai14-tdTomato (magenta) transgenic mouse brain immunolabeled for CalDAG-GEFI (cyan). C. Pnoc-Cre;Ai14 mouse section double-labeled for Pnoc (magenta) and tyrosine hydroxylase (TH, cyan) to identify dopaminergic cell bodies and dendrites. Labeling of striosome-dendron bouquets was found in all 4 mice used for cell counts.D. Nts-Cre;Ai14 line F mouse double-labeled for Nts (magenta) and TH (cyan) to identify dopamine cell bodies and descending dendrites. Lack of Nts-Cre;Ai14-tdTomato labeling of the striosome-dendron bouquet was found in all 3 mice used for cell counts. Cell nuclei stained with DAPI (blue). E. Percentage of striosome-marker-positive SPNs double-labeled for either D1 or D2/A2a (Pnoc-Cre;Ai14 n = 4 mice, Nts-Cre;Ai14;D1-GFP n = 2 mice and Nts-Cre;Ai14;D2-GFP n = 1 mice). For each mouse, three coronal sections (anterior, mid-level and caudal) were evaluated. Counts made only for cells within striosomes identified by MOR1.

To test whether the SPN transgene reporters in these lines were expressed in D1 and D2 SPNs in the striosomes, we generated double-transgenic mice with either D1 or D2 BAC (Bacterial Artificial Chromosome) reporter mouse lines and then counted single- and double-labeled SPNs at anterior, mid-level and posterior sections through the striatum (**Figure S1**). The P172-mCitrine and AT1-tdTomato lines drove reporter expression in both S-D1 and S-D2 SPNs, Pnoc-Cre and MOR1-mCherry labeled D1 SPNs, and a single line, the Nts-Cre;Ai14 reporter cross, drove expression selectively in D2 SPNs with striosome-enrichment. We for completeness re-examined the matrix immunomarker CalDAG-GEFI, already known to be expressed in both D1 and D2 SPNs^49,50^ as was GFP in the CalDAG-GEFI-GFP BAC line^51^.

Of the striosome-enriched lines, the Pnoc-Cre line (**Figures 1A, 1C and 1E, and S1J-S1P**) was the most selective and complete in marking D1-SPNs in striosomes^158^. We used this line for functional studies. We found D2-selective expression of the Nts-Cre line and used this line for S-D2 identification in functional studies (**Figures 1 B, 1D and 1E, and S1F and S1I**). For both Pnoc-Cre and Nts-Cre lines, we generated crosses with a LSL-FlpO mouse line^153^ (Pnoc;Flp and Nts;Flp respectively) to confer stable recombinase (Flp) expression in adulthood, as both lines are suspected or confirmed to have greater expression during development than subsequently. We also used these Cre lines in combination with the Ai14 tdTomato reporter mouse line to confer stable tdTomato expression in Pnoc and Nts populations.

The Nts-Cre;Ai14-positive D2 SPNs were vividly clustered in striosomes, with scattered positive cells also throughout a crescent-shaped dorso-anterior region of what appeared to be matrix but which overlapped the calbindin-poor extra-striosomal zone^52^ and Aldh1a1/Anxa1- positive fiber rich zone^53,54^, features of striosomes (**Figures 1B, 1D** and **1E**, and **S2J-S2L)**. These dispersed Nts-targeted SPNs appeared to comprise a small minority of SPNs in the crescent. We deemed this to be is a neurochemically specialized region of the mouse striatum, given its overlap with the Aldh1A1-rich, calbindin-poor dorsolateral zones previously noted^53–55^ and possibly corresponding in part to the so called ‘compartment-free space’^56^.

We found a remarkably clear double dissociation between outputs of the S-D1-expressing Pnoc line and the Nts-Cre;Ai14 S-D2-expressing line (**Figure 1**). The S-D2 SPNs (Nts SPNs) had a sharply defined projection to a central zone of the GPe (here named cGPe). This centralized projection was complementary to the classical indirect D2 pathway receiving zone, from the matrix, which was in the peripheral zone of the GPe (here named pGPe) (see **Figure 1** for anti-CalDAG-GEFI immunostaining, compare also CalDAG-GEFI-GFP fluorescence in **Figures S1Q and S2A-S2C**). The D1-selective Pnoc line SPNs projected to the striosome-dendron bouquets of the ventral SNpc (**Figure 1C**), a hallmark of D1 SPNs^42^ that was true of all of the D1 enriched lines (**See Figure S1Q for MOR1**). The Nts;Ai14 S-D2 line lacked a projection to the SNpc (**Figure 1D**). We conclude from this survey of seven mouse lines that D1 and D2 SPNs in striosomes project to different targets, the S-D1 SPNs targeting SNpc dopamine^42,106,158^, and S-D2 SPNs targeting the specialized central zone of the GPe.

### Retrograde rabies identification of the outputs of the cGPe and pGPe and comparison with the direct S-D1 outputs

As a baseline to use in identifying the outputs of the cGPe and pGPe zones, we began by using a selective trans-neuronal tracing method with rabies^30^ to compare the projections of the striosomes and matrix, based on prior evidence that striosomes, and specifically the D-1 expressing striosomes, project to the dopamine-containing SNpc, whereas the matrix projects primarily to the substantia nigra pars reticulata (SNpr)^42,106,158^. We focused on examining the distribution of GPe neurons that target the dopamine-containing neurons of the SNpc by using DAT-Cre mice (**Figures 2A-2F, and S3A**), or on the projection neurons of the SNpr, known to express parvalbumin (PV) by using PV-Cre mice (**Figures 2G-2L, and S3B**). In the same brains, we analyzed distributions of the rabies-labeled cells within striosome or matrix compartments. We injected Cre-dependent rabies helper viruses^30–32^ into the SNpc of DAT-Cre or SNpr of PV-Cre transgenic mice (**Table S1**), 21 days later injected EnvA-pseudotyped rabies virus (RVΔG- EGFP^148^ or RVΔG-mCherry^147^) at the sites of helper-virus injection to produce EGFP- or mCherry-expressing rabies in ‘starter neurons’ that could retrogradely spread the virus to their presynaptic partners^30–32^, and one week later prepared the brains for histology.

**Figure 2.**
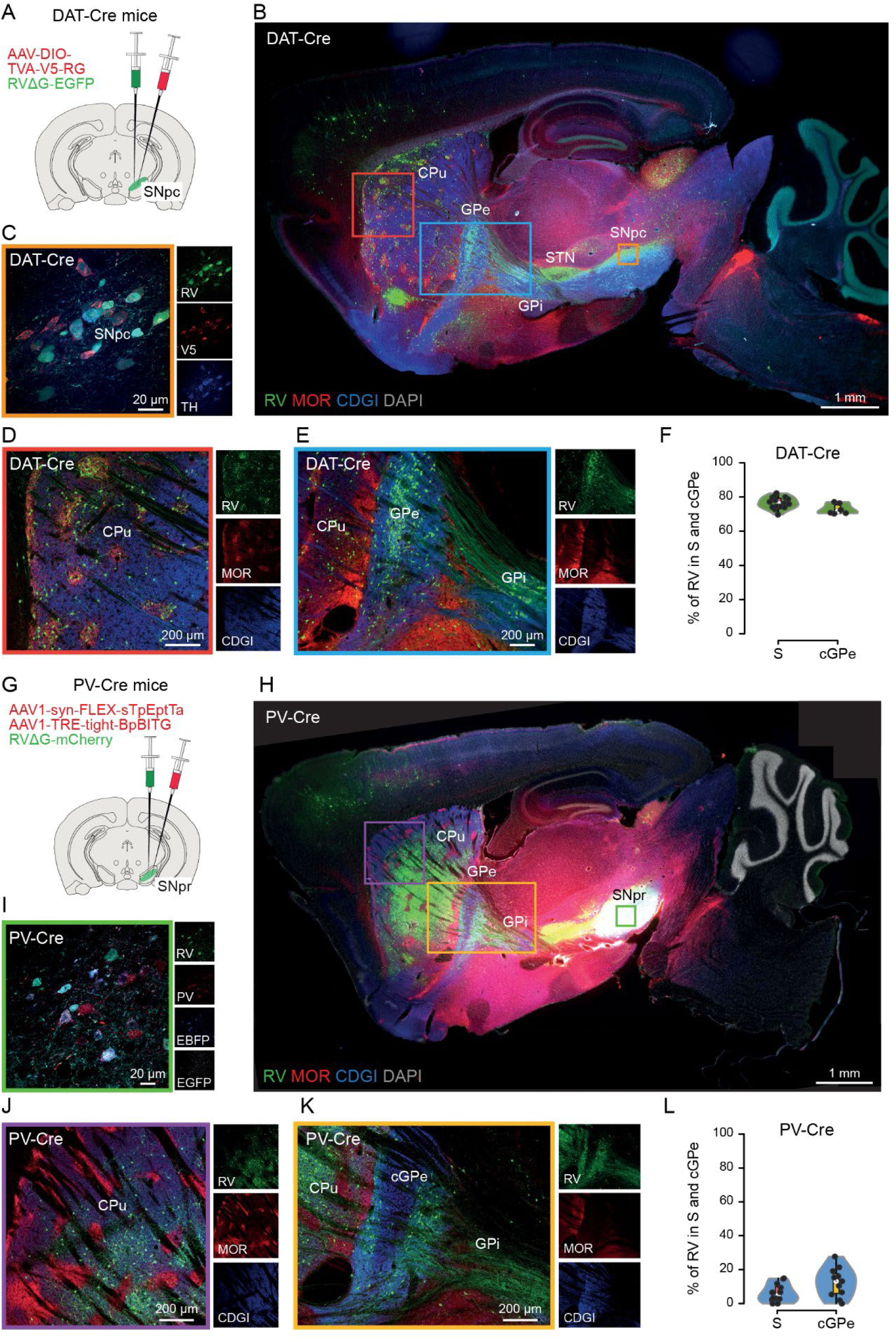
Distribution of RVΔG-labeled neurons targeting PV-positive neurons in SNpr or dopamine-containing neurons in SNpc. A-F. DAT-Cre RV tracing. A. Injection site and protocol. B. Sagittal section. RV-EYFP-labeled neurons counterstained with anti-MOR1 (striosome marker, red), anti-CalDAG-GEFI (CDGI, matrix marker, blue) and DAPI (gray). RV-EYFP-labeled presynaptic neurons (green) are located preferentially in striosomes and cGPe. CPu: caudoputamen. STN: subthalamic nucleus. C. SNpc injection site (orange box in B), with starter neurons co-expressing the RV (green), V5 (red) and tyrosine hydroxylase (TH, blue). D. Striatal region (red box in B). Presynaptic neurons projecting to dopamine cells are localized preferentially in striosomes. E. GPe region (blue box in B) showing that presynaptic neurons projecting to dopamine cells are distributed preferentially in the central MOR1-rich, CalDAG-GEFI-poor zone. F. Proportions of RV-labeled striatal (total n = 1544 neurons in 15 sections from 4 DAT-Cre mice) and GPe (total n = 658 neurons in 6 sections from 3 DAT-Cre mice) neurons targeting dopamine neurons in the SNpc that were identified in striosomes (S) and cPGe, respectively G-L. PV-Cre RV tracing. G. Injection site and protocol. H. Sagittal section depicting RVΔG-mCherry-labeled neurons (green) counterstained with anti-MOR1 (striosome marker, red), anti-CalDAG-GEFI (matrix marker, blue) and DAPI (gray). RVΔG-mCherry-labeled presynaptic neurons (green) are located preferentially in the matrix and the pGPe. I. SNpr injection site (green box in H), depicting starter neurons co-expressing the RVΔG-mCherry (green) and PV (red) together with EBFP and EGFP from the two helper viruses. J. Striatal region (purple box in H), showing presynaptic neurons projecting to SNpr-PV cells localized preferentially in matrix. K. GPe region (yellow box in H), showing presynaptic neurons projecting to SNpr-PV cells localized preferentially in pGPe, MOR1-poor/CalDAG-GEFI-rich zone. L. Proportions of RV-labeled striatal (total n = 1434 neurons in 12 sections from 3 PV-Cre mice) and GPe (total n = 305 neurons in 11 sections from 3 PV-Cre mice) neurons targeting PV neurons in SNpr located in striosomes and cGPe, respectively.

In the DAT-Cre mice, engineered to have Cre expression in dopamine-containing nigral neurons, rabies-expressing retrogradely labeled neurons were enriched in striosomes within the striatum (**Figures 2B, 2D and 2F**) and even more sharply so within a central zone of the GPe (**Figures 2B, 2E and 2F**). This zone corresponded to the region (cGPe) that we found to receive inputs from S-D2 striosomal SPNs in our experiments with engineered lines (**Figures 1B, S1 and S2**).

In the PV-Cre mice, with selective retrograde viral tracing from the PV-positive SNpr neurons, complementary distributions of labeled SPNs occurred both in the striatum and in the GPe (**Figures 2G-2L and S3B**). Retrogradely labeled SPNs targeting SNpr were largely distributed in the extra-striosomal matrix, mainly avoiding the striosomes (**Figures 2I, 2J, 2L and S3B**); and within the GPe, they were distributed largely in the peripheral zone that is, by immunohistochemical staining, a MOR1-poor and CDGI-rich zone (**Figures 2K, 2L and S3B**), the same territory receiving projections from the matrix (pGPe) (see **Figure 1** for anti-CalDAG-GEFI immuynostaining, compare also CalDAG-GEFI-GFP fluorescence in **Figures S1Q and S2A-S2C**). Fewer than 10% of the retrogradely labeled striatal SPNs appeared in striosomes, and fewer than 20% in the cGPe (**Figure 2L**; striatum: n = 1434 neurons, 12 sections, 3 PV-Cre mice; GPe: n = 305 neurons, 11 sections, 3 PV-Cre mice). In the DAT-Cre mice, most striatal neurons labeled were in striosomes in a topographically distributed dorsal region (nearly 80 %), and within the GPe, the labeled neurons were concentrated in the cGPe (∼80%), leaving its peripheral pGPe zone only sparsely labeled (**Figure 2F**; striatum: n = 1544 neurons, 15 sections, 4 DAT-Cre mice; GPe: n = 658 neurons, 6 sections, 3 DAT-Cre mice).

These retrograde rabies experiments confirmed a strong distinction between the striosomal and matrix projection patterns in the SNpc and SNpr, and further demonstrated an equally sharp output projection-based distinction between a central core of the GPe that projects to the dopamine-containing SNpc, corresponding to the D2-SPN cGPe projection zone, and a peripheral surrounding part of the GPe that projects to the PV-expressing nigral neurons of the SNpr and apparently corresponds to the matrix output zone. We note that the cGPe as defined by our genetic and rabies tracing experiments resembles the GPe described in previous anatomical work^33,34^. Further, a GPe-to-SNpc connection has been demonstrated electrophysiologically, but without specification of its topographic origins within GPe^35,36^. The clarity of the GPe core-periphery distinction found here is particularly notable given the high degree of collateralization of striatal projections to the GPe^7,21,25,29,37–41^.

### Striosomal SPNs target cGPe neurons that themselves target dopamine-containing nigral neurons

The input and output zones that we define appeared to strongly overlap, but this did not ensure that at a single-cell resolution there was direct cell-to-cell continuity. To probe the possibility that there is such direct connectivity from striosomes to the cells in cGPe that project to the SNpc, we labeled striosomal input fibers within GPe by their expression of N172, which labels both D1 and D2 striosomal neurons and is a strong marker of dorsal striosomes, (**Figures S1D, S1G and S1T**) in combination with rabies tracing of SNpc dopamine neurons in DAT-Cre/N172-tdTomato mice. Rabies-expressing SNpc-dopamine-targeting GPe neurons lay in close proximity to striosomal N172-labeled striosomal input fibers and putative endings in the cGPe zone (n = 6 DAT-Cre/N172-tdTomato mice; **Figures 3 and S4A-S4E**). We used synaptophysin targeting to synaptic terminals of the SPN inputs to the cGPe (**Figures 3D-3F**) combined with the SNpc retrograde rabies tracing. The results (**Figure 3F**) strongly supported direct synaptic terminal association of the striosomal inputs and GPe output cells projecting to the SNpc neurons: putative synaptic boutons of N172 striosomal axons were in contact, as seen at light microscopic levels, both with the cell bodies and the proximal dendrites of SNpc-dopamine-targeting cGPe neurons (**Figure 3F**). Approximately 80% of the N172 buttons, identified by shape and size (Imaris software) and co-localized with synaptophysin, lay in close proximity to RV-labeled dopamine-projecting cell bodies or processes in the cGPe. The majority of synaptophysin-positive N172 terminals were on the dendrites of the dopamine-projecting neurons, compared to the cell bodies (4 sections, average 228.5 ± 52.21sd N172-positive/SYP-positive/RV-positive dendrites, average 36.75 ± 10.7 sd N172-positive/SYP-positive/RV-positive cell bodies).

**Figure 3.**
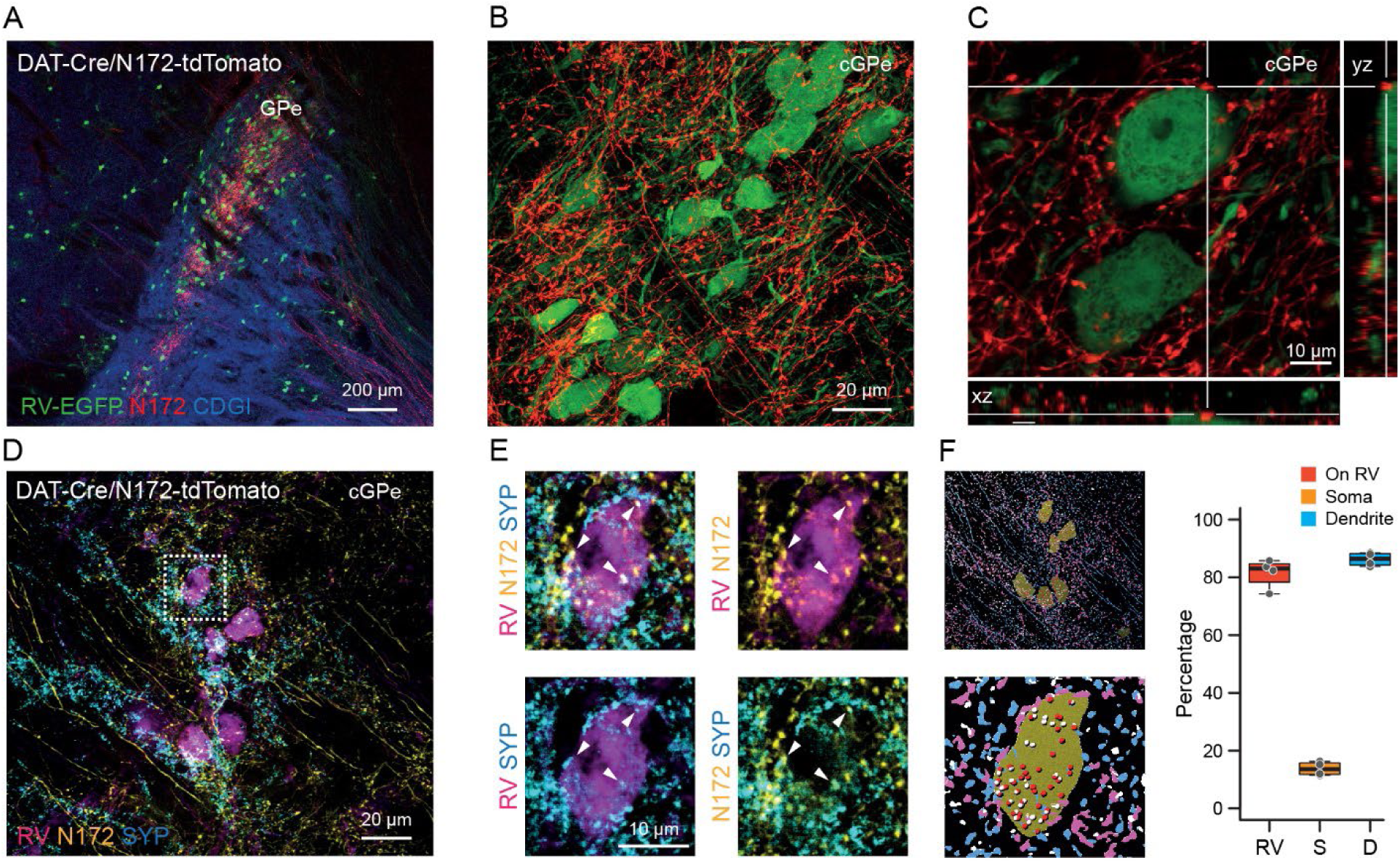
Striosomal SPNs target dopamine-projecting cGPe neurons. A. Terminals from N172-tdTomato-labeled striosomal neurons (red) on SNpc-dopamine-targeting cGPe neurons (RV-EGFP-labeled neurons, green). N = 6 mice. B. Magnification of two SNpc-targeting cGPe neurons with contacts from N172 striosomal neuron processes. White lines indicate the plane of the Z-stack projection for yz and xy image coordinates. C. Synaptophysin (SYP)-positive (blue) N172 striosomal neuron terminals (yellow) on SNpc-targeting cGPe neurons (magenta). D. Magnification of cell indicated by square in D. E. Left: Representative images (top: same cells as in D, bottom: mugnification of one cell) demonstrating the identification of boutons based on N172-positive/SYP-positive markers, as well as their sphericity and volume (see methods). Right: Quantification of colocalized terminals (4 sections) demonstrating that the majority of identified boutons are associated with RV-labeled cGPe neurons and that from these the majority of them were identified on their dendritic processes. RV: The percentage of N172-positive/SYP-positive boutons found on RV-labeled cells. S: The percentage of N172-positive/SYP-positive/RV-positive terminals found on the cell body of RV-labeled cells. D: The percentage of N172-positive/SYP-positive/RV-positive terminals found on the dendrites of RV-labeled cells.

### Rasa3 reporter line highlights cGPe zone

We looked for markers that would label the cGPe selectively to facilitate definition of striosome-cGPe connectivity and throughput to the SN. GPe projection neurons have been distinguished according to multiple markers including PV, somatostatin, Lhx6, Npas1, and FoxP2^24–29,57–59^. None of these markers, individually, defined a cell type that was differentially enriched in the cGPe, nor in the surrounding peripheral region around it (pGPe), according to examination of immunohistochemical labeling for Lhx6 or PV, or by detection of tdTomato reporter in Lhx6-Cre;Ai14, PV-Cre;Ai14 and Npas1-Cre;Ai14 mice. These findings (not illustrated) are consistent with those from comprehensive spatial mapping of these GPe neuronal markers^24–29^. We did identify one GFP reporter, the Rasa3-GFP BAC transgenic mouse, with strikingly enriched expression in the central GPe (**Figure S4F)** Rasa3 co-localized with retrogradely labeled SNpc-projecting neurons, and were not PV-positive (**Figures S4E and S4G-S4K**). However, this was true for the BAC reporter but not for expressed Raza3, which was not selective for cGPe and was down-regulated during development (**Figures S4L and S4M**). We could therefore not use this line experimentally, but the augmented labeling in the Rasa3-GFP BAC mouse indicated that the cGPe harbors a molecularly distinct cell type, differentially enriched relative to the surrounding peripheral (pGPe) regions. This well could become important in future experiments on development, as Rasa3 is a transcription factor associated with the immune system^61–63^.

### Demonstration of novel striatal enhancers preferentially expressed in striosomes or matrix: confirmation of striosomal direct and indirect pathways

We tested several enhancers developed at the Allen Institute (**Figure S5**): the striosomal enhancer DLX2.0-SYFP2 and the matrix-specific enhancer 452h-mTFP1 in Nts;Ai14 mice. The striosomal enhancer DLX2.0-SYFP2 showed an overlay of expression with Nts;Ai14 in striosomes (**Figure S5B**). The matrix-specific enhancer 452h-mTFP1 exhibited a distinct expression pattern in the matrix regions relative to the Nts;Ai14 SPNs (**Figure S5C**).

We also investigated the targeted expression in D1 or D2 striosomal SPNs in A2a-Cre mice. Intrastriatal injection of AAV-DLX2.0-minBG-FIPO-WPRE3-BGHpA was followed by intersectional viruses (Cre-On/Flp-On-EYFP for striosomal D2 and Cre-Off/Flp-On-mCherry for striosomal D1). This protocol resulted in the selective labeling of striosomal D1-SPNs with mCherry (red) and striosomal D2-SPNs with EYFP (green) within the striatum. The axonal projections of both D1 and D2 striosomal neurons highlighted dense terminal zones of S-D2 in the cGPe and the continuation of D1 striosomal neuron axons downstream to the GPi and substantia nigra (**Figure S5D**). Specific targeting of striosomal D2 SPNs in A2a-Cre mice was confirmed following an injection of the Cre-dependent striosomal enhancer virus DLX2.0-DIO-SYFP2, with green fluorescence marking the expression of the enhancer virus and red immunostaining for MOR1 confirming striosome specificity (**Figure S5E**). These findings provided important further evidence for a central zone of the GPe (cGPe) having differentially strong input from the S-D2 pathway, but having far less input from the M-D2 pathways as detected by our methods.

These multiple sets of experiments demonstrate that, in parallel with the canonical matrix D2 projection to the GPe, the origin of the classical indirect pathway of the basal ganglia, there is a non-canonical S-D2 pathway targeting a GP region distinct from the classical GPe-indirect pathway pallidum. The S-D2 indirect pathway innervates the dopamine-containing ventral tier of the SNpc, with no detectable labeling of the SNpr. By contrast, the matrix M-D2 pathway directly targets PV neurons of the SNpr.

### Functional differentiation of the striosomal direct and indirect pathways: contrasting effects on striatal dopamine release of the S-D1- and S-D2 enriched populations

We anticipated that, given their targeting of dopamine-containing neurons in the striatum in the SNpc ventral tier, the S-D1 and S-D2 pathways might each have identifiable effects on dopamine release in the striatum. We recorded striatal dopamine release in 9 Pnoc;Flp (S-D1 proxy) and 11 Nts;Flp (S-D2 proxy) mice as they moved voluntarily within a small open field. One month before testing, the mice received injections of Flp-dependent AAV-fDIO-Chrimson and AAV-hsyn-GRAB-DA3h and placement of an optogenetic stimulation fiber close by (**Figure 4A**; mid-mediolateral dorsal caudoputamen: AP = 0.86 mm, ML = 2.0 mm, DV = 2.2 mm). During the free open-field behavior, we applied optogenetic stimulation (8 sec, 40 Hz, 5 msec pulls duration, 10 trials with randomized inter-trial intervals of ∼1.5 min) to stimulate with AAV-fDIO-Chrimson as we recorded dopamine release with DA3h. We measured running trajectory lengths during the 8-sec stimulation periods and the 8-sec periods before and after the stimulation, and we compared the distance traveled and the activity levels during these times as analyzed with DeepLabCut and B-SOiD (**Figures 4, S7D and S7E**).

**Figure 4.**
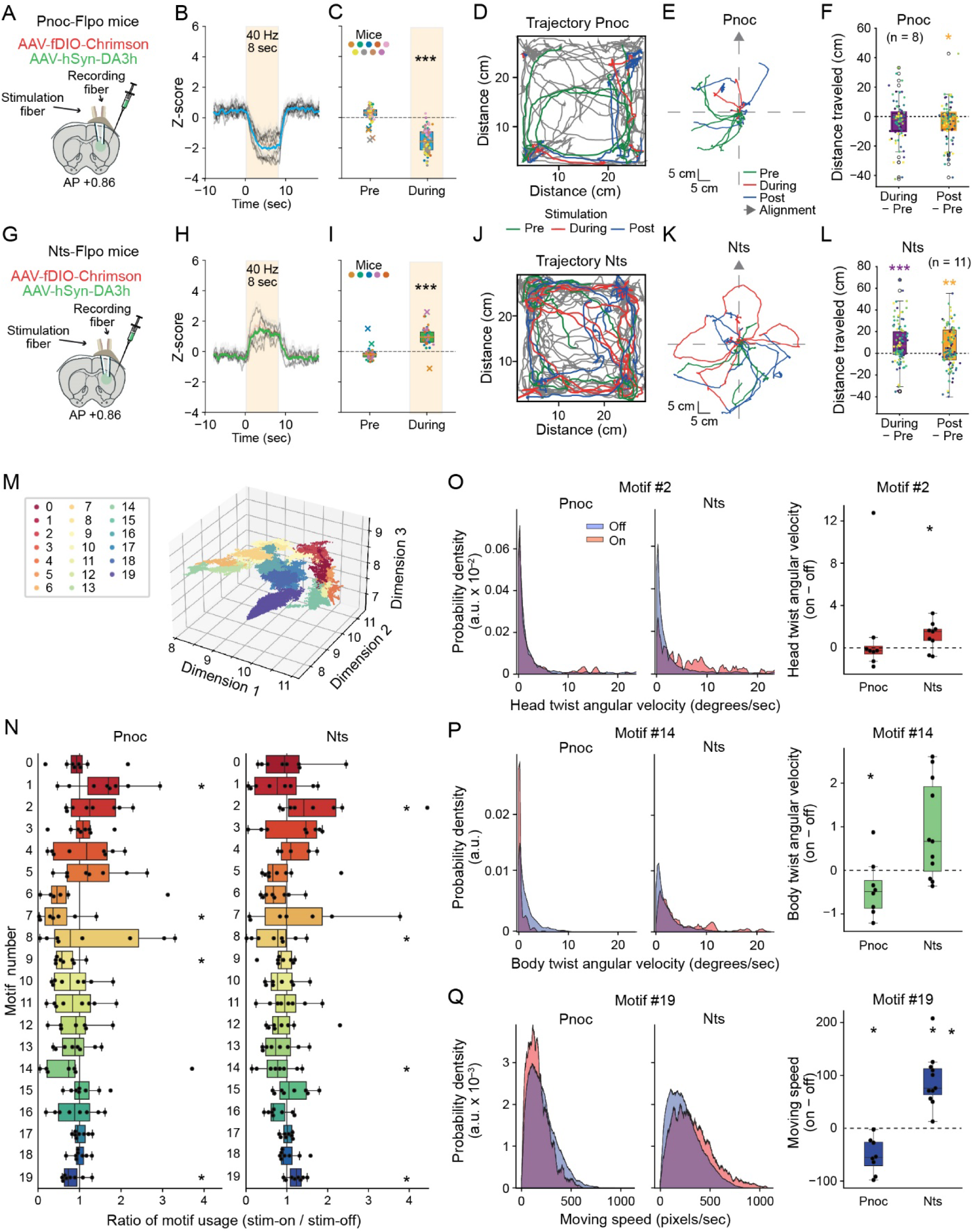
S-D1 and S-D2 effects on dopamine release and motor kinematics. A. Protocol for targeting S-D1 (Pnoc) SPNs for optogenetic stimulation. B. Average dopamine response to optogenetic stimulation (8-sec train at 40 Hz, 10 trials) C. Average response before (Pre) and during stimulation. D. Trajectories of mice freely moving in open field box (30 x 30cm) before (green), during (red) and after (blue) optogenetic stimulations of S-D1 SPNs. E. Trajectories before, during and after optogenetic stimulation for 10 trials reoriented to the axis defined by the base of the tail to neck one frame before the stimulation. F. Distance traveled during and after the optogenetic stimulation relative to equivalent time-period before stimulation. G. Schematic of protocol for targeting S-D2 SPNs. H. Average response to optogenetic stimulation (8-sec train at 40 Hz, 10 trials) of S-D2 (Nts) SPNs. I. Average response before and during the stimulation. J. Behavioral effects of optogenetic manipulation of S-D2 SPNs in self-paced actions. Conventions as in D. K. Traces around the optogenetic stimulation, as in E. L. Difference of distance traveled as in F. M. Behavioral motif clusters in 3D space using UMAP. Twenty unique motifs were identified by the B-SOiD clustering analysis. Each color represents a different behavioral motif cluster. N. Box plot illustrating the ratio of motif usage between stimulation-on and stimulation-off periods for 8 Pnoc and 11 Nts mice. Average motif usage rates were measured in frames per sec, calculated separately for each condition and analyzed for statistical differences with two-tailed t-test across different conditions (*p < 0.05) O-Q. Left: Distribution density of head twist angular velocity, defined by the angle between the direction from the center of the body to neck and from the neck to snout (O), body twist angular velocity (P) and moving speed, quantified by tracking the center of the body (Q) from 8 Pnoc an 11 Nts mice. Data were aggregated from all mice and analyzed separately for periods of stimulation-on and stimulation-off. Right: Box plots illustrating the ratio of average angular velocities and moving speed during stimulation-on versus stimulation-off periods, calculated for each mouse and statistically compared (*p < 0.05 by two-tailed t-test).

We stimulated either the putative S-D1 or putative S-D2 populations in 9 Pnoc;Flp mice and 5 Nts;Flp mice (with sufficient dopamine sensor signal) at randomly chosen times as the mice moved about. Stimulating the S-D1 (Pnoc;Flp) population decreased dopamine release (**Figures 4A-4C**), whereas stimulating the S-D2 (Nts-Flp) population increased dopamine release (**Figures 4G-4I)**.

This evidence that the S-D1 and S-D2 pathways had near opposite effects on striatal dopamine release, with the S-D1 pathway decreasing dopamine release, and the S-D2 pathway increasing release. We then simultaneously measured locomotor activity during the 8-sec pre-stimulation, 8-sec during stimulation, and 8-sec post-stimulation periods. When the S-D1 proxy Pnoc populations were stimulated, the mice decreased their movement trajectories relative to those during the pre-stimulation period (n = 8 Pnoc;Flp mice, **Figures 4D-4F, S7D and S7E**). When the S-D2 proxy Nts populations were stimulated, large increases in movement occurred (n = 11 Nts;Flp mice, **Figures 4J-4L, S7D and S7E**). These effects of stimulating the S-D1 and S-D2 populations were opposite to the classical effects on movement of the canonical D1-direct (‘Go’) and D2-indirect (‘No-Go’) pathways^1,2,6,18^.

Behavioral motifs, characterized using the B-SOiD clustering tool, identified twenty unique movement motifs from video recordings of the mice as annotated by DeepLabCut. The average frequency of these motifs, calculated in frames per second, showed variable usage between stimulation-on and stimulation-off periods (**Figures 4M-4Q**) and overall, a significant shift occurred during stimulation periods compared to off periods. With Pnoc-targeted activation, there was a reduction in time spent in mobility, along with an increase in sniffing, rearing and grooming (see motifs 1, 5, and 19 in **Figure 4N**). Conversely, Nts-targeted activation resulted in the opposite effect, increasing mobility and decreasing sniffing, rearing and grooming activities (**Figures 4M and 4N**). Kinematic analyses complemented these findings by quantifying head and body twist angular velocities, as well as moving speeds (**Figures 4O-4Q**). These measurements of the active behavior of the putative S-D1 and S-D2 populations again indicated that Pnoc stimulation reduced different aspects of movement, whereas the Nts activation increased these. We interpret this finding as indicating an alteration in behavioral states due to stimulation, with a notable difference in the influence of the S-D1 and S-D2 SPN populations within the caudoputamen.

### Dynamics of dopamine release in relation to the activities of S-D1 and S-D2-enriched SPN populations during probabilistic decision-making in a T-maze task

It has not been feasible with previous techniques to record selectively the D2 populations of striosomal neurons in awake behaving animals. Nor has such dual recording been possible with simultaneous recordings of dopamine release. We attempted to do this. We examined striatal neural population activity and dopamine release simultaneously in 5 Pnoc;Flp (proxy S-D1) mice and 6 Nts;Flp (proxy S-D2) mice that performed a well-learned T-maze task requiring them to turn right or left to receive a reward with a 75% probability for arriving at the correct port. No reward was given for an incorrect choice, and there was a 5% chance of receiving an air puff. The rewarded port was switched randomly every 7 to 24 trials, with no signal provided to indicate the change. The mice were implanted and injected bilaterally, medial striatum in the left hemisphere and lateral striatum in the right hemisphere, with optical probes to record striatal dopamine release (DA3h) and AAV-syn_FLEX-jGCaMP8m to record calcium transients of striosomal SPNs (**Figures 5A, 5B and S6**). Injection sites were not at medial or lateral extremes of the caudoputamen.

**Figure 5.**
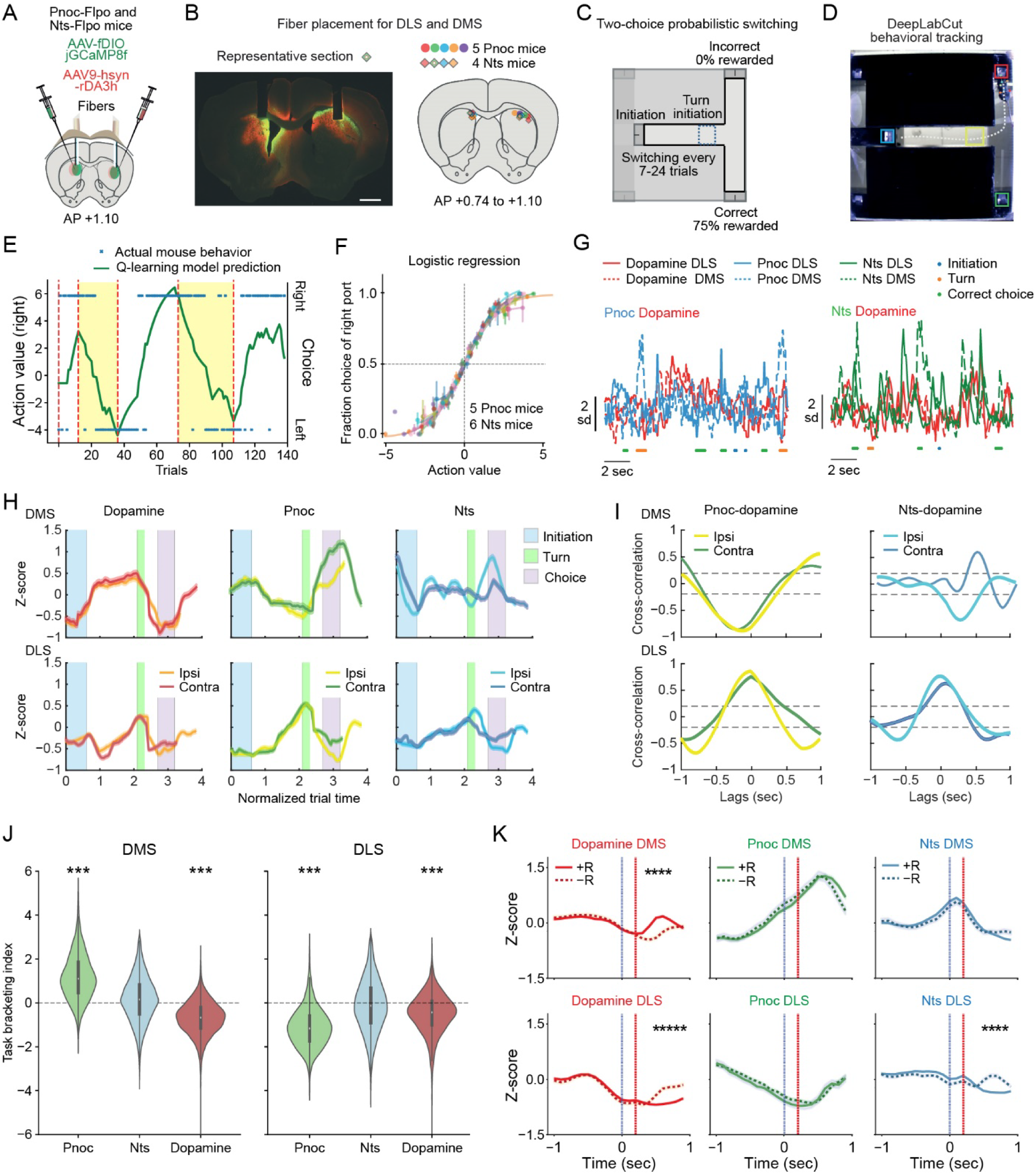
Simultaneous imaging of dopamine release and S-D1 (Pnoc proxy) or S-D2 (Nts proxy) neuronal activity during a 2-choice probabilistic switching task. A. Protocol for simultaneous imaging of dopamine release and neuronal activity in DMS and DLS. B. Cross-section from a representative Nts-Flpo mice illustrating fiber placements for simultaneous imaging of dopamine release and Nts neuron activity in the right DLS and left DMS (left), and mapping of fiber placement for all Nts-Flpo and Pnoc-Flpo mice used (right). C. Diagram of the 2-choice probabilistic switching task. D. Frame from behavioral recording annotated with DeepLabCut. E. Example of actual mouse behavior in an expert session (blue crosses) versus the fitted trial-by-trial Q-learning model (green trace) during 140 trials with reward at right (no shading) or left (yellow shading) port. Dashed red lines mark the switch of rewarded ports. F. Fraction of choices for the right port plotted against the relative action value (Q-learning), illustrating decision-making strategies. G. Examples of photometry traces showing dopamine level and Pnoc or Nts activity recorded from the DMS and DLS during task performance. H. Average photometry traces (mean ± SEM) showing activity of dopamine, Pnoc and Nts recorded in the DMS (top) and DLS (bottom) during the task. Activity was averaged separately for ipsilateral and contralateral choice trials, and normalized for time. Shading indicates the initiation, turn and choice phases of the trial. I. Cross-correlation analysis between Pnoc or Nts neuronal activity and dopamine signaling in the DMS (top) or DLS (bottom). Pnoc (S-D1) and dopamine signals were anticorrelated with a prominent inverse peak for DMS and positive peak in DLS with a negative lag, suggesting dopamine fluctuations precede changes in S-D1 neuron firing rates by ∼0.2 sec. By contrast, Nts (S-D2) neuron activity and dopamine signaling showed a positive correlation for contralateral choice trials and for both ipsilateral and contralateral choice trials in DLS and a negative correlation with ipsilateral choice trials in DMS. Peaks in the Nts neuron activity lead the increases in dopamine levels in DLS and lags in the DMS, with a lag time of ∼0.4 sec. J. Task-bracketing index for Pnoc (S-D1), Nts (S-D2) and dopamine. Pnoc has a positive bracketing index in DMS and negative index in DLS. Dopamine has a negative bracketing index for both. No significant bracketing index was found for Nts neuron activity. K. Pnoc and Nts activity do not dissociate rewarded and unrewarded outcomes. Dopamine shows a positive reward response only in the DMS. Dopamine release and Nts activity in DLS show increased activity in unrewarded trials (see Figure S7 for alignment to the port exit).

As the mice ran the maze task **(Figures 5C-5F)**, the levels of dopamine release in both the dorsomedial (DMS) and dorsolateral (DLS) striatum briefly fell at the initiation port but then steadily rose to a sharp peak just at the maze turn. Levels then fell to baseline levels at the choice ports and then rose after the choice and in the return to the initiation port (**Figures 5H and S7**). Thus, dopamine levels were highest during the runs as the mice accelerated to the turn point and then fell as they ran to the end port. Remarkably, as this strong modulation of dopamine levels occurred in the striatum, sharply different population activities occurred in SPNs of the S-D1 Pnoc- and S-D2 Nts populations and, as measured by recording of calcium transients, in the DMS and DLS (**Figures 6H and S7**).

**Figure 6.**
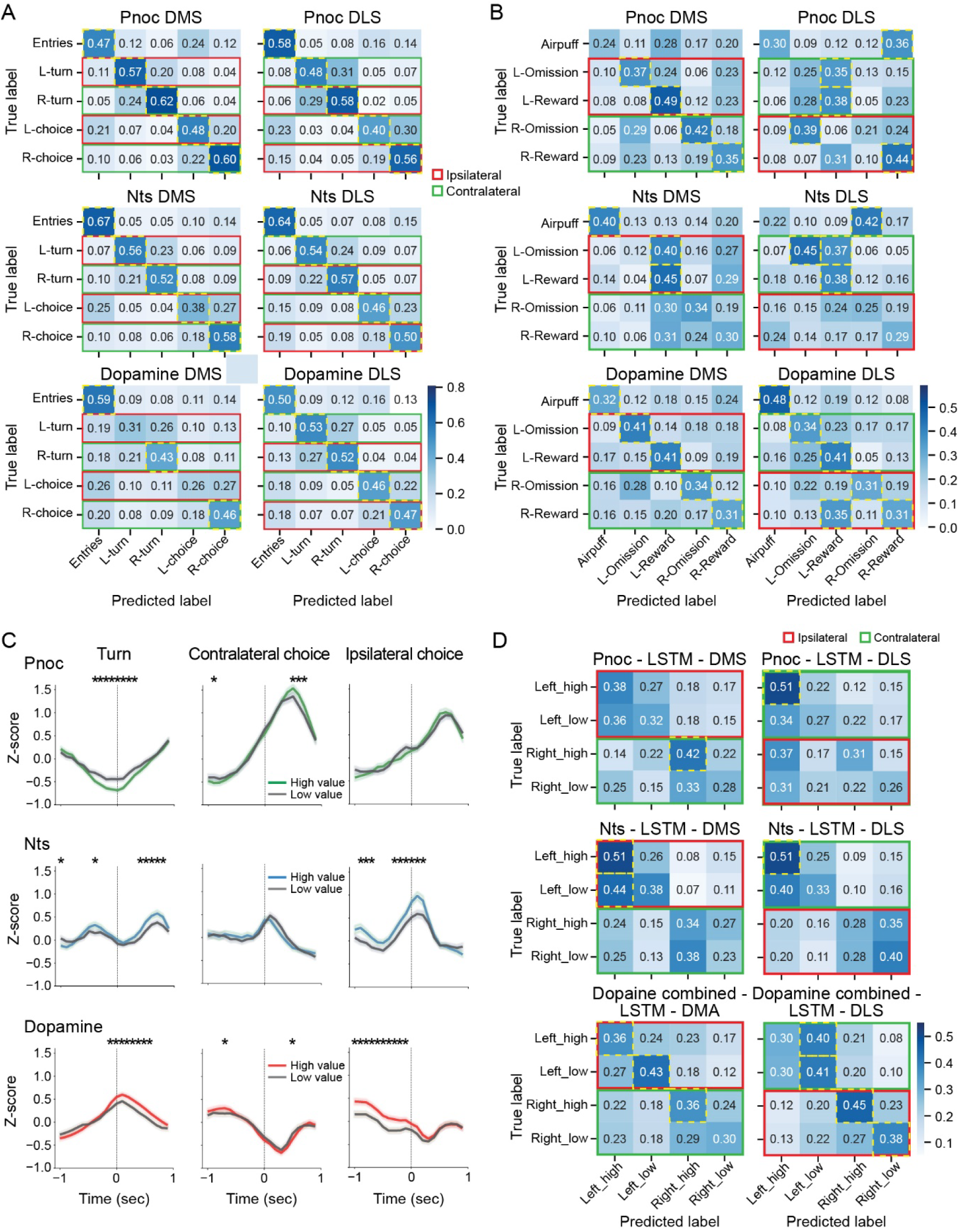
Performance of neural activity pattern classification in predicting task behaviors, reward outcomes and action values across Pnoc (S-D1), Nts (S-D2) and dopamine release signals in DMS and DLS. A. Confusion matrices show the accuracy of an LSTM model in classifying task space events (initiation, left turn, right turn, left choice and right choice) based on Pnoc (top), Nts (middle) and dopamine (bottom) signals recorded in DMS (left) and DLS (right). Red and green squares indicate, respectively, ipsilateral and contralateral events relative to the recording site. Dashed yellow squares highlight task events identified with significantly higher accuracy than chance. Values are averaged across all mice. Except for dopamine release in the DMS, all signals show higher-than-chance accuracy for all task events (diagonal). Dopamine release in the DMS shows significantly higher-than-chance accuracy for initiation and only for contralateral turn and choice events. B. Confusion matrices showing the accuracy of an LSTM model in classifying outcomes at choice ports (air puff, left rewarded, left unrewarded, right rewarded and right unrewarded). The same color coding and highlighting are used as in A. Dopamine provides the best performance, but overall accuracy is lower compared to task events classification in panel A. C. Average signals of Pnoc, Nts and dopamine release during turn, contralateral choice and ipsilateral choice, split by high and low action value trials as calculated with a Q-learning model. Significant differences in amplitude were found at different time points between high and low value trials across all signals. D. Confusion matrices using the same method and color coding as A and B for classifying left high value choice, left low value choice, right high value choice and right low value choice. This panel shows that the Pnoc (S-D1) signal in both DMS and DLS carries action value information only for high value contralateral actions. Nts (S-D2) shows poor separation of high and low value trials, and although dopamine contains value information, its accuracy is relatively low.

In the DMS, the activity of the S-D1 Pnoc population rose at the start, fell to a deep valley of low activity at the turn, and rose toward the choice point and reward ports. Then it fell to baseline levels. The end rise at the choice ports was even larger than the rise at the initiation site. This is notable because at the reward port the mice received larger reward (5Ml) but it was probabilistic reward, whereas at the initiation port they received less reward (3ml) but it was give 100% of entried. Raw traces of the activity of the S-D1 Pnoc-targeted and S-D2 Nts-targeted striatal neurons (**Figures 6H and S7**) indicated that the activity of the putative S-D1 cells in the DMS formed a pattern bracketing the runs with high activity at the beginning and end of the run trajectories (**Figure 5J**), highly reminiscent of the task bracketing pattern that we originally observed in the dorsal striatum of rats, mice and monkeys^126–137^. The S-D2 proxy Nts striatal population had a fluctuating fall-rise-fall pattern in the DMS, roughly opposite to the S-D1 proxy Pnoc population rises at the beginning-and-end pattern.

In the DLS, no such bracketing by the Pnoc population occurred. Instead, both the Pnoc and Nts populations started at very low levels and formed a broad rise and then fell to low levels at goal-reaching similar to the dopamine response (**Figure 5H**)^64^.

Cross-correlation analysis between the dopamine release and population activity levels of both Pnoc and Nts mice during the maze task performance (**Figure 5I**) confirmed that in the DMS, the dopamine fluctuations and Pnoc S-D1 population neuron activity exhibited an anti-correlated pattern. Conversely, in the DLS, the pattern for the Pnoc population showed a positive correlation with dopamine fluctuations, similar to that of the Nts population. The Nts population in the DMS showed lateralization effects: there was a positive correlation with dopamine for trials with contralateral turns and a negative correlation for ipsilateral trials.

With alignment activity to the choice port entry, we observed no significant modulation in response to the outcome for either the Pnoc or Nts populations in either the DMS or DLS across both and unrewarded trials (**Figure 5K**). This surprising result suggests a lack of selectivity in the outcome/reinforcement responses of the Pnoc and Nts populations: their activity at outcome was nearly identical regardless of reward condition. By contrast, dopamine release levels in the DMS were higher in rewarded trials than in unrewarded trials, indicating a positive modulation by outcome. Also unexpectedly, in the DLS, both dopamine release levels and Nts population activity were increased (positively modulated) in unrewarded trials. This activity was better aligned with the timing of port exit rather than with the reinforcement outcome itself, potentially indicating an ‘end signal’ or ‘try-again signal’ or response to the movement of exiting the port. The mice tended to leave the port more rapidly when reward was received (**Figure S7B**).

We used a classifier model to decode behavior and the outcome of choices. Dopamine in the DMS showed significantly higher-than-chance accuracy for initiation and contralateral turn and choice events, whereas Pnoc and Nts in both DMS and DLS, and dopamine in DLS, showed higher-than-chance accuracy for all task events (**Figure 6A**). Notably, dopamine showed better accuracy for predicting reward choices (**Figure 6B**).

We analyzed S-D1 and S-D2 neuronal activity patterns in relation to historical reward data derived from Q-learning algorithms, categorizing trials as high-value and low-value trials (**Figures 6C and 6D**). We observed that all signals were modulated, with significant differences in amplitude, by value during the turn phase, with a mixed preference for contralateral actions. The Pnoc (S-D1) activity showed significant modulation during contralateral choices, whereas the Nts (S-D2) activity was more active during ipsilateral choices. Additionally, dopamine release was modulated during ipsilateral choices, particularly before entry into the choice zone (**Figure 6C**). Using a classifier model to decode value (**Figure 6D**), we found that the Pnoc (S-D1) signal in both DMS and DLS carried action value information only for high-value contralateral actions. The Nts (S-D2) signal showed poor separation of high and low value trials, and although dopamine release encoded value information, its accuracy remained relatively low. Overall, dopamine served as a better predictor for value than either S-D1 or S-D2 activity, though its performance was still limited.

## DISCUSSION

Our findings introduce the concept of parallel direct and indirect basal ganglia pathways for both striosome and matrix compartments. These findings at once demonstrate striosomal direct-D1 and indirect-D2 pathways as parallel streams and suggest that the two great divisions of the striatum, the canonical direct D1-direct D2-indirect organization and the compartmentalization into striosomes and matrix, can be conceived of as intersecting both anatomically and functionally. First, we delineate, as a pair, direct-D1 and indirect-D2 pathways that arise in the striosomes of the dorsal striatum. This pair of pathways echoes the organization of the canonical direct-D1 and indirect-D2 motor pathways of the basal ganglia in having both direct and indirect access to their target destinations. But the striosomal direct-indirect pathways terminate in the dopamine-containing SNpc instead of the pallidonigral motor output nuclei of the basal ganglia, as do the canonical direct-indirect pathways arising in the matrix compartment. Second, we find for the functions of the striosomal pathways that they have opponent processing as do the canonical pathways, but their actions are the reverse of the canonical pathways’ actions. The striosomal D1 pathway tends to *reduce* movement and to reduce dopamine release, and the D2 striosomal pathway tends to *increase* movement and dopamine release—functions opposite to those of the canonical D1 and D2 pathways (**Figure 7**).

**Figure 7.**
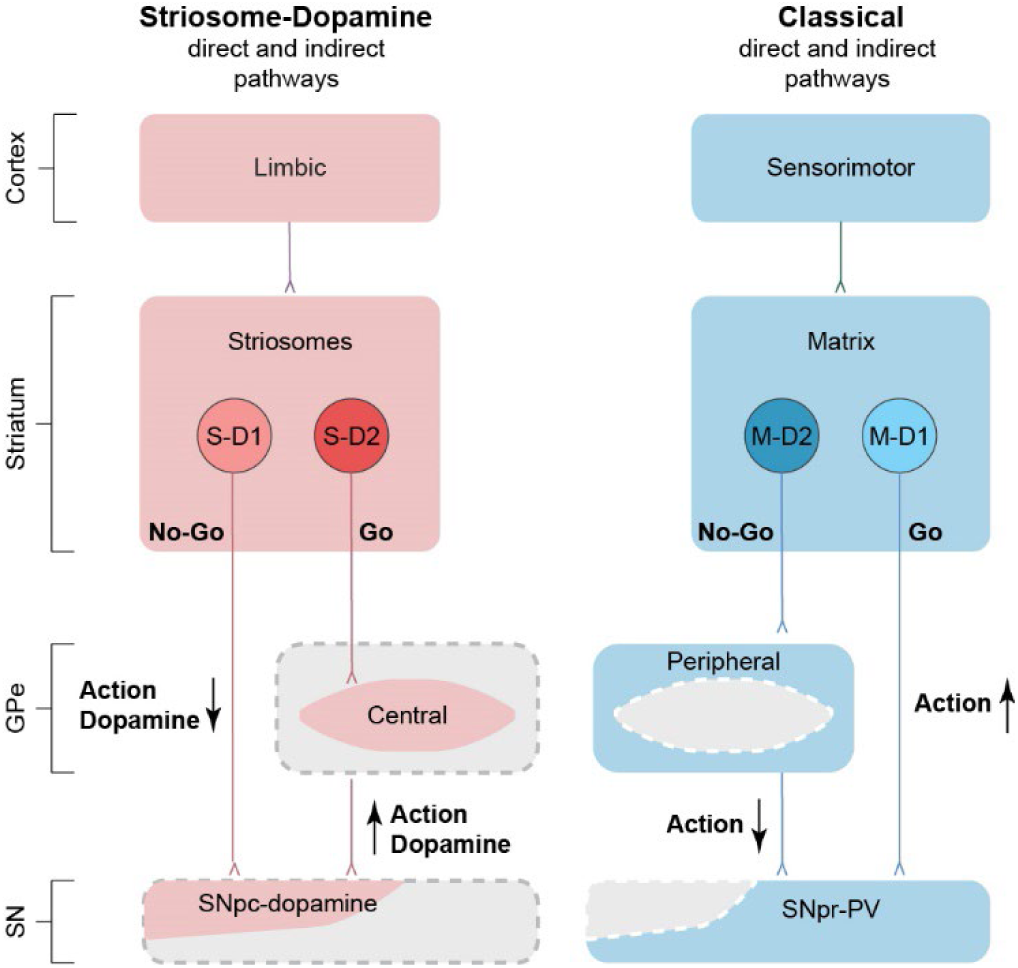
Dual direct-Indirect pathway model for basal ganglia output pathways modulating behavior. The striosome and matrix compartments are the sources of parallel D1-direct and D2-indirect pathways to the SNpc (striosomes) and to the SNpr (matrix). The matrix direct-indirect pathways modulate movement; the striosomal direct-indirect pathways, considering their links to the limbic system, could modulate mood state and motivation to act, collectively functions affected by basal ganglia disorders. SNpr-PV: substantia nigra pars reticulata parvalbumin-positive neurons; SNpc-dopamine: substantia nigra pars compacta dopamine-containing neurons.

Evidence across rodent and non-human primate species has suggested that striosomes might be particularly important for energizing or engaging beneficial, reward-producing task-related behavior^65^, for decisions involving motivationally challenging approach or avoidance in situations with conflicting favorable and unfavorable options^66–69^, and for behavioral challenges present along the anxiety-depression and repetitive behavior-addiction axes^66,68,70,71^. Our findings thus raise the possibility that the basal ganglia have in place two major direct-indirect systems to exert opposing control over behavior: one targeting motor output nuclei modulating action, and the other, from striosomes, targeting the nigral dopamine-containing neurons that modulate mood state and the motivation to act, and that likely influence the classical direct-indirect circuits.

At the most general level, these findings provide a framework for aligning cortico-basal ganglia-thalamo-cortical and brainstem circuits and their modulation by dopamine according to both of the great organizational systems of the striatum dividing the output of its projection neurons into a direct-indirect pathway system and an apparently orthogonal striosome-matrix compartmentalization. These organizational systems come together in the view proposed here, a finding that suggests an elegant evolution of circuit design in the forebrain.

We are aware that there are many other circuits in the complex basal ganglia connectome. For instance, the projection from striosomal cells to border GPi cells that project to the lateral habenula^24,32,72,78^, a node in GPi output circuits that has been implicated in negative reward signalling^32,73,74^ and depression^73,159^. Here we focus on the conclusion supported by our anatomical and functional findings, that there are parallel striosome-derived and matrix-derived direct and indirect pathways differentially targeting motor output neurons (matrix) and dopamine neurons (striosomes). This model raises the possibility that evolution devised a way to coordinate the different functional manifolds of the organizational systems, one centered on motor control and the other on modulating mood and motivational modulation of physical and cognitive action.

Our work builds on pioneering studies reporting in different segments of the pathways delineated here. Fujiyama and colleagues^34,40^ and Kita and Kitai^55^ found striatal neurons projecting to the zone of the pallidum likely corresponding to the GPe here demonstrated as a central, striosome-recipient zone of the GPe. The identity and dopamine expression characteristics of these SPNs was not known. Gittis and colleagues^28^ and Nelson and colleagues^80^ described a pallidal input to nigral dopamine neurons withoutidentifying the GPe cells of origin with respect to their striatal inputs but with identification of their projection to the cell bodies of the dopamine neurons. Evidence now even suggests that, due to extensive collateralization of the classic D1 direct pathway^21^, all GPe cells could receive D1 as well as the traditionally recognized D2 inputs. Exhaustive reviews of the connectivity of the pallidum have now further enriched appreciation of the complex anatomy of the pallidum^25,29^. What we emphasize here is evidence for synaptic throughput from D2 receptor-expressing striosomal SPNs to the central GPe zone (cGPe) and then to the SNpc of DAT-Cre mice. This D2 indirect pathway formed the missing circuit of what we now know to be a partner to the known S-D1 circuit, forming a complete D1-D2 direct-indirect striosomal system alongside the classical D1-D2 direct-indirect system.

### Effects of the S-D1 and S-D2 pathways on dopamine release in the striatum

The S-D1 and S-D2 pathways, as represented by their enriched Pnoc (S-D1 proxy) and Nts (S-D2 proxy), have largely opposing actions on striatal dopamine release, on self-paced free locomotor movement, on behavior in the probabilistic decision-making task, including on presence or absence of task-bracketing of the Pnoc and Nts population activity. Yet these are time-varying relationships, and they differ for the more medial and more lateral parts of the dorsal striatum. For example, the task-bracketing pattern for the Pnoc SPNs in the DMS is not apparent for the DLS Pnoc populations examined simultaneously alongside the DMS Pnoc neurons. These DMS-DLS differences extend much pivotal previous work on differences between signaling in the DMS and DLS neurons unmarked with respect to their striosomal or matrix origins^81–89^, thus indicating that topographic constraints are a general feature of striatal circuits. Our findings also extend pioneering efforts to identify functional characteristics of D1 and D2 neurons in the striatum ^1–22,41,83,86,98,106^. Our work builds on these and other studies.

### Striosomal direct and indirect pathways are parts of the nigro-striato-nigral loop

The nigro-striato-nigral loop is central to nearly all work on the basal ganglia related to disorders of mood and movement. These include neurodegenerative diseases, as witnessed by Parkinson’s disease and Huntington’s disease^90–92^, and by a range of neuropsychiatric conditions^93–98^ and states of addiction^66,68,70,71^. The nigro-striato-nigral loop is also central to reinforcement learning and normal adaptive behavioral control^99–101^. Here we introduce a new view of the loop control pathways that can affect levels of dopamine in the striatum. Remarkably, despite the relatively small total volume of the striosomal system as compared to that of the matrix around it, striosomes appear to have a strong influence on the nigrostriatal dopamine system and thus on corticostriatal and other inputs to the striatum^71,102^ as well as local circuits of the striatum^103,104^ that are modulated by and themselves modulate dopamine. It is thought that striosomes are the main or at least highly predominant source of striatal input to the SNpc^42,80,105^, although there are indications^106^ (and unpublished observations) that this view may neglect some inputs from the matrix. The S-D1 pathway to the SNpc has already been implicated in the effects of psychomotor stimulants^107–110^, opioid receptor-mediated negative affect^111^, the craving induced by fentanyl withdrawal^111^, and behavioral stereotypy^119^. Further, this S-D1 pathway to the striosome-dendron bouquets is associated with the expression of endocannabinoid CB-1 receptors^112^, and constitutive deletion of these receptors negatively affects the bouquet system^102^. There is a pressing need now for further work to discover more about the striosomal control in different behavioral contexts and in different normal and abnormal conditions.

### S-D2 circuit emphasizes potential target of dopamine D2 receptor actions of clinical and therapeutic significance

A rich body of pharmacologic work and pioneering therapeutic treatments of basal ganglia disorders target D2 receptors^113–116^. Here, we present evidence of a previously unrecognized D2 circuit arising from SPNs in striosomes that targets a specialized central region of the external pallidum that we here call the core region of the GPe (cGPe) by reference to the murine GPe. The S-D2 SPN cell type has explicitly been singled out by single-nucleus RNA-sequencing (snRNA-seq) as the most vulnerable cell type in the brains of Huntington’s disease mouse model mutants^117^, a disorder in which striosomes can be differentially affected in patients exhibiting mood symptoms^118,119^. This vulnerability could affect the S-D2 indirect pathway identified here, leading to altered modulation of dopamine functions as well as corticostriatal and thalamostriatal function. More generally, effects on this indirect S-D2 circuit and its extension to dopamine-containing nigral neurons could account for some of the effects, including unwanted side effects, of a broad range of D2 agonist or antagonist treatments used in clinical neurology and psychiatry. More highly targeted therapies directed at the S-D2 or M-D2 pathways could improve treatment efficacy, if the differential functions of striosomes and matrix as we and others have so far found in mice^65,68,69,111,160^ hold true within the D2 population and also holds true in humans.

### Opposing pathways

The classical direct-indirect model heavily relies on the clinical symptomatology of neurodegenerative disorders such as Parkinson’s disease and Huntington’s disease^1,2,6^. In the dopamine-depleted state, lower dopamine tone decreases activity in the ‘Go’ (D1) pathway, and increases activity in the ‘No-Go’ (D2) pathway, favoring bradykinesia. In Huntington’s disease, preferential depletion of the No-Go (D2) pathway disinhibits motor commands, resulting in hyperkinesia. Supporting this standard model, experimental evidence indicates that the function of the classic direct pathway is to disinhibit motor neurons, for example, to release motor commands by inhibiting omnipause neurons in the motor output nuclei of basal ganglia^120,121^. By contrast, the function of the indirect pathway has remained obscure: for example, how it inhibits unwanted or irrelevant movements (as by targeted coactivation^18,122^), and how its deficits are manifested as cholera, tics or dystonia. Here we show the striosomal system can exhibit effects opposite to those of the canonical pathways, the S-D1 suppressing and S-D2 pathway facilitating locomotion and associated movements. How the preferential depletion of S-D2^117^ contributes to early-stage symptoms of Huntington’s or Parkinson’s disease awaits discovery, but one possibility is that this depletion contributes to deficits in incentive motives for actions (e.g., apathy and depression) or psychiatric symptoms (e.g., psychosis, delusions) observed years before clinical onset^123–125^.

### Timing in the striosomal circuits targeting dopamine-containing neurons of the substantia nigra

Work in slice preparations suggests the GPe input to the SNpc dopamine neurons terminates on the cell bodies of these neurons^28,33,106^. These connections likely include the S-D2 pathway inputs identified here. This situation could set up a potentially important way that the S-D1 and S-D2 pathways operate, given that the S-D1 pathway terminates mainly on the descending dendrites of ventral tier SNpc neurons, including dendrons of the striosome-dendron bouquets as well as the still mysterious ‘posterior cell cluster’ neurons as they are currently denominated. Different timing of these somatic S-D2 and dendritic S-D1 inputs could be critical to the functions of the nigrostriatal control circuits, in coordination with timed inputs from the cerebellum^154^. Striosomal direct-indirect circuits with different lags could dynamically adjust the correlations between S-D1 and S-D2 effects in relation to dopamine release, leading to differential control of the canonical direct and indirect pathways.

### Task-bracketing and inverse task-bracketing

We found in the DMS an inverse relationship between the activity of the S-D1 and S-D2 population activities during performance of the probabilistic decision-making task. For presumed driving of contralateral striatal function, the S-D1 (Pnoc) population activity peaked at the beginning and end of the runs, including the entire out-back run sequence, whereas the S-D2 (Nts) population mainly peaked at the turn. Opposite effects present for task-bracketing by populations of striatal neurons were first noted in tetrode recording patterns^126–132^: such bracketing patterns have been found in recordings in the macaque prefrontal cortex^133^ and in the striatum bracketing pattern in rats, mice^132,134–137^ and macaques^138^. In the rodent tetrode studies^127^, the task-bracketing was found in the DLS, but not in the DMS, for which the activity instead resembled the S-D2 recordings in the DLS (gradual rise and fall) and for which the decline appeared to exert a permissive role in allowing behavioral expression of the bracketing^127^ .These recordings did not differentiate D1- and D2-expressing populations or striosomal and matrix sub-populations, but did demonstrate inverse mid-run peaks in co-recorded striatal PV interneurons^135^.

Analysis by training classifiers suggested that dopamine in the DMS exhibited significantly higher-than-chance accuracy for initiating actions and making contralateral choices, whereas Pnoc signals in both DMS and DLS exhibited robust predictive power across all task events. The predictive accuracy of dopamine for reward-related events was the best among the signals but was overall lower than its ability to predict task events. Remarkably, neither Pnoc nor Nts striosomal populations showed significant predictive power for reward-related outcomes: the encoding of action value was primarily present in the dopamine signals, not in the Pnoc and Nts striosomal population signals. These results clearly differentiate the population encoding on the part of striosomal S-D1 and S-D2 populations and the encoding by dopamine release recorded in the same striatal subdivisions. Task events preceding reward can be decoded by the S-D1 and S-D2 population activities, but it is the dopamine release that can be used to decode subjective outcome value.

### Evolutionary perspective evoked by the current findings

We find a similar circuit design for the classic direct-indirect basal ganglia pathways and the striosomal direct-indirect pathways delineated here. The many overlapping and bridging collateral innervations in the GPe and elsewhere could be seen as diluting the “direct” and “indirect” pathway concept beyond usefulness^7,21,22,40^. But this is not so when attention is given to the precise and differential targeting of these pathways at their terminations. We have scarcely any full accounting of even the basic anatomy of these pathways in different species. The snRNA-seq data available suggest conservation of S-D1 and S-D2 identities from mouse to human, though with many different mRNAs expressed across these species^117,141,161^. In the human brain, there are three distinguishable segments of the pallidum. Whether these relate to our findings is unclear but of obvious interest. Very little is known about the posterior cell cluster, a major target of the S-D1 pathway along with the striosome-dendron bouquet system^42^, and there are frequently found contrasts between the DMS and DLS in rodents, as we found here^81–89^. These have possible, but not definitely identified, correlates in, respectively, parts of the caudate nucleus and putamen of primates including humans.

### Future work and limitations to be addressed

Our experiments leave many questions for future work, and we note limitations of this initial delineation of the paired striosomal direct S-D1 and indirect S-D2 pathways. We used proxies for the S-D1 and S-D2 SPN populations, because we could not use direct labeling of the S-D1 and S-D2 populations given that the direct and indirect matrix pathways also terminate in or pass through the GPe. These proxies are clearly imperfect. Pnoc-Cre targets only ∼40% of S-D1 neurons (**Figure S1**), and the Nts-Cre expression pattern includes the scattered, un-clustered neurons that occupy the dorsolateral calbindin-poor crescent. Some non-Pnoc neurons express *prodynorphin* (*Pdyn*), reported to be associated with reward and appetitive responses^160^, others express *Tshz1* ^160^, reported to be related to punishment and avoidant behavior, and some Pnoc reporter-positive neurons express both Pdyn and Tshz1^158,160^. Pnoc-Cre targeted neurons did not exclusively overlap with either *Pdyn* or *Tshz1* labeled S-D1 neurons^158^.

A major knowledge gap is comparable information about the matrix. Considering that the volume of the matrix population exceeds that of the striosomal system in most parts of the striatum, it might therefore dominate the results of studies not differentiating the two compartments. Our coverage here of the dorsal striatum was limited; but the sizable differences between activity in DMS and DLS sites align with evidence^81–89^ that topographic specializations are of paramount importance in analyzing the striatum, as in the neocortex^142–145^. Finally, many other circuits can influence the striatum and its interactions with the nigral dopamine system, including circuits that link striatum with the cerebellum^154^. Our work thus only introduces the S-D1 and S-D2 striosomal direct and indirect pathways as paralleling the canonical M-D1 and M-D2 pathways; we look forward to comprehensive comparisons between these pairs of direct-indirect pathways and computational models based on our findings. The evidence for differential circuit targeting and functions of the striosomal D1 and D2 pathways reported strongly suggest a need for fundamentally revised models of basal ganglia function. It is hoped that this work can benefit the clinic and the millions of individuals affected by disorders referable to basal ganglia dysfunction.

## ACKNOWLEDGMENTS

This research was supported by NIH/NIMH R01 MH060379 (AMG), NIH/NIMH P50 MH119467 (AMG), the Saks-Kavanaugh Foundation (AMG), the William N and Bernice, E. Bumpus Foundation (AMG, AM), Jim & Joan Schattinger (AMG), the Hock E. Tan and K. Lisa Yang Center for Autism Research (AMG), Mr. Robert Buxton, the Simons Foundation grant to the Simons Center for the Social Brain at MIT (AMG), the CHDI Foundation (AMG), Ellen Schapiro & Gerald Axelbaum Investigator BBRF Young Investigator Grant (IL), and NIH BRAIN Armamentarium Grant UF1MH128339 (JTT). We thank Henry Hall for his help in many ways; Samitha Venu and the Swanson Biotechnology Center at the MIT Koch Institute for expert assistance in generating the N172 mouse line; Ximena OptizAraya for cloning of enhancer AAV vectors; Allen Institute Viral Core team for enhancer AAV virus production; and Bargavi Thyagarajan for Allen Institute project management support.

## AUTHOR CONTRIBUTIONS

JRC and AMG conceived and designed the initial experiments demonstrating anatomically the striosomal direct and indirect pathways; IL designed and implemented the rabies experiments and demonstrated the throughput from S-D2 cells to cGPe to SNpc; VS implemented rabies experiments; JRC, TY, IL, ArM and JHL collected anatomical data; IL, JRC, EH, TY, ArM and AMG performed image analysis interpretation and cell counting; IL, ArM collected behavioral and functional data; IL, GA, KH, KP performed behavioral and signal analysis; GA developed and employed analytic pipelines and used multiple models to analyze the functional data in relation to the neural data; GA and IL performed the statistical analyses; KM supervised VS and provided viral reagents; JTT and IRW designed and generated viral reagents; JRC and AMG wrote an initial draft and AMG and IL wrote the full draft with edits by EH, AyM, and JRC.

## Methods

### Mouse husbandry

All procedures were approved by the Committee on Animal Care at the Massachusetts Institute of Technology, which is AAALAC accredited. **Table S1** summarizes the mouse lines used. Experimental mice were female and male and maintained on a standard 12/12 hr light/dark cycle with free access to food and water. P172 mice were evaluated at age 3-5 weeks. The other mice were evaluated at ages ranging from 2-10 months of age. Table S1 describes the various transgenic mice used.

### Generation of AT1-tdTomato BAC transgenic mice

The BAC containing AGTR1 (RP24-63B19) was obtained from the BACPAC Resource Center at CHORI. Plasmids for cloning and recombineering were obtained from The Rockefeller University and Jonathan Ting then in the laboratory of Professor Feng of the MIBR at MIT. Two flanking sequences for homologous recombination consisted each of 500 basepairs, one upstream of the ATG start codon and the other beginning in the following intron, in order to remove the first coding exon and prevent overexpression. The iCre-P2A-tdTomato expression cassette was amplified from pAAV-EF1a-iCre-P2A-tdTomato. PCR-amplified and gel-purified Homology Box A and iCre-P2A-tdTomato were inserted into iTV vector by In-Fusion (Clontech) and transformed into DH10B bacteria. In-Fusion was then performed between the iTV-iCre-P2A-tdTomato-Box A and homology box B and the resulting vector was linearized and purified for homologous recombination. The Agtr1 BAC was transformed into el250 bacteria and selected on chloramphenicol-media plates from which colonies were picked and grown overnight in LB-Lennox at 32°C. 400 ml of culture was transferred to 20 ml of LB-Lennox and placed at 32°C until A600 = ∼0.4. Cells were then transferred to Eppendorf tubes and heat shocked at 42°C for 15 min with occasional shaking. The bacteria were cooled for 5 min on wet ice and then centrifuged at 10,000 x g for 15 sec. Washes with cold 10% glycerol were performed twice and the cells were resuspended in the remaining volume. Linearized iTV-iCre-P2A-tdTomato-Boxes vector was electroporated (1.75 kV, 25 µF and 200 Ω) into the Agtr1-BAC bacteria, put on ice for 2 min, resuspended in 1 ml of LB-Lennox, and incubated at 32°C for 2 hr. Cells were then pelleted at 4000 rpm for 4 min, plated onto LB-agar chloramphenicol/kanamycin plates and incubated at 32°C overnight. Colonies were picked and grown overnight in 5 ml of LB, glycerol stocks made and a columnless Qiagen miniprep was performed. Bacteria were streaked out onto chloramphenicol kanamycin LB-agar plates from the glycerol stocks of positive clones. Colonies were selected and grown in 1 ml LB + chloramphenicol/kanamycin at 32°C for 6-8 hr with addition of arabinose during the last hour to induce Cre expression and removal of the neomycin cassette. Bacteria were plated and colony PCR was used to identify neomycin-negative clones that were inoculated into 5 mL LB (Miller) with chloramphenicol and grown for 8 hr and 32°C. This culture was added to 200 mL LB (Miller), grown overnight and used for DNA preparation with the Nucleobond Xtra BAC kit. Approximately 7 mg BAC DNA was then linearized with NotI. The linearized DNA was then run on PFGE in a single well. The next day, the sides of the gel were stained, the unstained was aligned, and the appropriately sized band was excised. The gel slice was cut in half length-wise and put into Spectra/Por Dialysis tubing (MWCO:6-8,000) to which 500 ml TE was added. The clamped tubing was put into a pulsed field gel electrophoresis box perpendicular to the flow of the current and run under the following settings: 35 sec initial, 35 sec final and a run time of 8 hr. After 8 hr, the tubing was rotated 180° and run for another 6 min. The eluate was removed from the tubing and then run over a column prepared as follows: add water to the column, add 1.5 ml G50 sephadex beads to column, fill column with 1x TE. Samples were collected in 2-drop fractions and 5 ml of each fraction was run on an agarose to find the 5 most concentrated fractions, which were then combined. Microinjection buffer (100 mM NaCl, 10 mM Tris HCl, 0.1 mM EDTA, pH7.4) was filtered through a 0.2um syringe filter and put into a beaker. A spot dialysis disc (0.025 µm VSWP) was placed on top of the microinjection buffer, and then sample was put on the disc. The sample was left to dialyze overnight at 4°C. AT1-BAC-tdTomato DNA (0.5–1 ng/ml) with polyamines added 1 week prior, was given to the MIT transgenic facility for C57B6/N pronuclear injection. Multiple founder mouse lines were screened for optimal expression of tdTomato in histological brain sections.

### Generation of rabies viruses

Cloning of AAV genome plasmids pAAV-syn-FLEX-splitTVA-EGFP-tTA (Addgene 100798) and pAAV-TREtight-mTagBFP2-B19G (Addgene 100799) has been described^146^. These genomes were packaged in serotype 1 AAV capsids by, and are available for purchase from, Addgene (catalog numbers 100798-AAV1, and 100799-AAV1) and diluted in Dulbecco’s phosphate-buffered saline (DPBS) (Fisher, 14-190-250) by factors of 1:200 and 1:10, respectively, to final titers (determined by Addgene by qPCR) of 8.5 × 10^10 gc/ml and 1.6 × 10^12 gc/ml, respectively, then combined in a 50/50 ratio by volume as described 56 before injection. EnvA-enveloped rabies virus RVΔG-4mCherry (EnvA)^147^ (Addgene #52488) and RVΔG-EGFP^148^ (Addgene #52487) was produced as described previously^147–150^ but using helper plasmids pCAG-B19N (Addgene #59924), pCAG-B19P (Addgene #59925), pCAG-B19G (Addgene #59921), pCAG-B19L (Addgene #59922), and pCAG-T7pol (Addgene #59926) for the rescue step^150^, with a final titer of 1.70 × 10^10 infectious units/ml as determined by infection of TVA-expressing cells as described previously^148^.

### Striatal cell type enhancer discovery

The DLX2.0 enhancer is a 3x concatenated core of the hDLX enhancer and has been previously described in terms of neocortical GABAergic cell type specificity and rapid onset of expression in human *ex vivo* brain slice cultures^140,141^. Here we describe for the first time the unique enrichment of transgene expression driven by the DLX2.0 enhancer in striosomes of the dorsal striatal region following mouse *in vivo* stereotaxic injection. The striosome enrichment pattern was not observed following intravenous virus administration using PHP.eB serotype in mice (data not shown). Striatal brain region and cell-type-specific enhancers 452h and 444h were identified from Roussos lab Brain Open Chromatin Atlas (BOCA) resource^155^, a publicly available human postmortem bulk dataset using assay for transposase accessible chromatin with RNA sequencing (ATAC-seq) profiling. The BOCA resource covers various cortical brain regions, amygdala, thalamus, hippocampus, striatum brain regions and clustering of neuronal vs. non-neuronal cell types. We identified putative enhancers based on differential chromatin accessibility in putamen and nucleus accumbens regions of the human striatum relative to other brain regions, as well as proximity to known marker genes for major striatal medium spiny neuron types (e.g., DRD2 as a marker gene for indirect pathway SPNs).

### Enhancer AAV cloning

Candidate enhancers were PCR amplified from human genomic DNA and cloned using standard restriction enzyme digestion and ligation into AAV expression vectors upstream of a minimal beta-globin promoter (minBG) and mTFP1, bright monomeric teal fluorescent protein that is well tolerated in neurons. The resultant AAV plasmids were rAAV-452h-minBG-mTFP1-WPRE3-BGHpA (Addgene plasmid #191708, alias AiP12700) and rAAV-444h-minBG-mTFP1-WPRE3-BGHpA (Addgene plasmid #191729, alias AiP12965). To investigate Cre- and Flp-dependent intersectional strategies for striosome labeling, we additionally constructed rAAV-DLX2.0-DIO-SYFP2 (plasmid AiP14533) and rAAV-DLX2.0-FlpO-WPRE3-BGHpA (plasmid AiP4532) vector designs by standard subcloning.

### Virus production

We prepared endotoxin free maxiprep DNA for packaging AAV plasmids into PHP.eB serotypes AAV particles. For initial enhancer-AAV screening by intravenous delivery in mouse we generated small-scale crude AAV preps by transfecting 15 µg maxiprep enhancer-reporter DNA,15 µg PHP.eB cap plasmid, and 30 µg pHelper plasmid into one 15-cm dish of confluent HEK-293T cells using PEI-Max. At one day post-transfection the medium was changed to 1% fetal bovine serum, and after 3 days the cells and supernatant were collected, freeze-thawed 3x to release AAV particles, treated with benzonase (1 µl) for 1 hr to degrade free DNA, then clarified (at 3000 g for 10 min) and concentrated to approximately 150 µl by using an Amicon Ultra-15 centrifugal filter unit at 5000 g for 30-60 min (NMWL 100 kDa, Sigma #Z740210-24EA). For large-scale gradient preps for intraparenchymal injection into mouse brain, we instead transfected 10 x 15- cm plates of HEK-293T cells and purified the cell lysates by iodixanol gradient centrifugation. We titered both crude and gradient AAV preps by digital droplet PCR.

### Stereotactic injections with viruses

AAV9-CMV-Flex-synaptophysin-mCherry (1 ml, 2X1013 vg/ml, purchased from Dr. Rachael Neve) was injected into the dorsal striatum (AP: 1.2 mm, ML: ±1.4 mm, DV: −2.0 mm) of one AT1- Cre-tdTomato line F mice, and 5 weeks later the brain was harvested and processed for immunolabeling with dopamine transporter (DAT). The mCherry fluorescence was imaged directly so as to minimize cross-detection of AT1-tdTomato, which was relatively weak.

PV-Cre or DAT-Cre heterozygous mice or double transgenic PV-Cre;P172-mCitrine (hemizygous) and DAT-Cre; P172-mCitrine (hemizygous) were kept under deep anesthesia with a continuous flow of 2% isoflurane (Southmedic Inc.) in an oxygen mixture, delivered by a nose- cone attached to a stereotax. Mice were given intranigral injections of a mixture of pAAV-syn-FLEX-splitTVA-EGFP-tTA and pAAV-TREtight-mTagBFP2-B19G in 0.4 ml DPBS, via NanoFil microsyringe (World Precision Instruments) at the following stereotactic coordinates: AP = −2.7 mm, ML = ±1.7 mm and DV = −3.7 mm to target the ventral SNpc in DAT-Cre mice, and AP = −2.8 mm, ML = ±1.7 mm and DV = −4.1 mm to target the ventral SNpr in PV-Cre mice. Rabies virus, in 0.4 ml DPBS, was injected into the same site 7 days later and brains were perfused 7 days after rabies injection. Stereotactic coordinates were modified slightly in single-transgenic mice that were older, to account for their larger brain size (< 4-week-old mice are necessary for P172-mCitrine transgene imaging).

### Tissue preparation, immunolabeling and microscopy

Mice were anesthetized with Euthasol (Virbac AH Inc.; pentobarbital sodium and phenytoin sodium) or isoflurane and then trans-cardially perfused with 0.9% saline, followed by 4% (wt/vol) paraformaldehyde in 0.1 M NaKPO4 buffer solution (PBS). Brains were then dissected, post-fixed for 90 min, stored in 20% (vol/vol) glycerin sinking solution overnight or longer, and cut into transverse 30 µm sections on a freezing microtome. Sections were stored in 0.1% sodium azide in 0.1 M PBS made from NaKPO4 until use.

Free-floating sections were rinsed 3 times for 2 min each in 0.01 M PBS with 0.2% Triton X-100 and then were blocked for 20 min in TSA Blocking Reagent (Perkin Elmer). Sections were incubated with primary antibodies (**Table S2**) suspended in TSA Blocking Reagent overnight at 4°C on a shaker. Following primary incubation, sections were rinsed 3 times for 2 min each in 0.01 M PBS with 0.2% Triton X-100 and were incubated in Alexa Fluor secondary antibodies (Thermo Fisher Scientific) (**Table S2**) suspended in TSA Blocking Reagent for 2 hr at room temperature. Following secondary incubation, sections were rinsed 3 times for 2 min each in 0.1 M PBS. For confocal microscopy, sections were mounted on subbed glass slides and coverslipped with ProLong antifade reagent with DAPI (Thermo Fisher Scientific).

A Zeiss Axiozoom microscope was used to obtain wide-field images with standard epifluorescence filter sets for DAPI (365 excitation, 395 beamsplitter, 445/50 emission), eGFP/AF488 (470/40 excitation, 495 beam splitter, 525/50 emission), tdTomato/AF546 (550/25 excitation, 570 beamsplitter, 605/70 emission) and Cy5/AF647 (640/30 excitation, 660 beamsplitter, 690/50 emission). Confocal imaging was performed with a Zeiss LSM510 with diode lasers (473, 559, 653 nm) for excitation. Images were collected with a 10x 0.4 NA objective and a 60X silicon oil 1.3 NA objective lens. Optical sectioning was optimized according to the microscope software. Images were processed and analyzed with Fiji software. Maximum intensity projection images were generated from optical sections using Image J. Background was subtracted and contrast was adjusted for presentation. Figures were prepared with Photoshop and Adobe Illustrator 6.0.

### Cell counting in transgenic mice

Two female and two male mice were used for each mouse line, and three coronal sections, taken from the anterior, mid-level and caudal regions, were evaluated for each mouse. Neurons were counted in all of the MOR1-identified striosomes that were dorsal to the anterior commissure. A composite image of the 3 channels GFP (green), tdTomato (red) and MOR1 (cyan) was created in the Fiji software. Individual and merged channels were viewed with the ‘channels’ menu, and the brightness and contrast of the image was adjusted as necessary with the ‘brightness/contrast’ menu. To divide the dorsal striatum into medial and lateral sides, a horizontal line was drawn across the widest part of the striatum with the ‘straight line’ tool. A vertical line was drawn to bisect the striatum, and then it was traced (pink) with the pen tool. The cyan channel only was viewed, and the MOR1 striosomes were outlined (teal) with the pen tool. To define the area of the matrix, we outlined matrix with the ‘freehand selections’ tool, and moved the selected area to an area next to the striosomes for measurement of an equivalent area. For each mouse line, cells were counted in the following categories: GFP-labeled cells in the MOR1 striosome, GFP-labeled cells in the matrix, GFP and tdTomato double-labeled cells in the total MOR1 striosomes and matrix, GFP single-labeled cells in the total MOR1 striosomes and matrix, double-labeled cells in the MOR1 striosomes, GFP single-labeled cells in the MOR1 striosomes, double-labeled cells in the matrix, GFP single-labeled cells in the matrix, double-labeled cells in the matrix, tdTomato-labeled cells in the MOR1 striosomes, and tdTomato-labeled cells in the matrix (P172 and AT1LineF).

### Photometry recordings

Calcium activity was recorded utilizing GCaMP8s and the red-shifted dopamine sensor DA3h with two-color bundle-imaging fiber photometry system (Doric lenses). This setup allowed for the sequential recording of fiber photometry fluorescence data at a sampling rate of 10 Hz across three distinct channels: a reference channel (400-410 nm), a green channel (460-490 nm), and a red channel (555-570 nm). Video recordings of mouse behavior were synchronized with photometry signals, capturing the xy-coordinates of five body parts: the tip of the tail, the base of the tail, the center of the body, the neck and the snout/head, using DeepLabCut software. Both video and photometry frames, alongside task events or optogenetic stimulation, were timestamped, employing Bonsai software, which interfaced with external hardware via an Arduino microcontroller operating on the Firmata-plus protocol.

The preprocessing of photometry recordings involved scaling the 400-410 nm control signal to the 460-490 nm neuronal activity signal or 555-570 nm dopamine sensor signal via linear least-squares regression, followed by subtraction to eliminate motion and autofluorescence-related artifacts. The baseline fluorescence of the activity signal was estimated over time using a least-squares regression line applied to the scaled control signal. This corrected activity signal was then normalized against the raw baseline estimate, resulting in a ΔF/F trace that was adjusted for photobleaching, motion, and autofluorescence.

### Open field

In the open field, mice were subjected to overnight water deprivation freely explored a 30x30 cm arena for 15 min, allowing evaluation of locomotor activity. The floor, made of clear transparent acrylic, facilitated video capture from below using a high-resolution Oryx 10GigE camera (Flir), recording at 30 Hz synchronized with fiber photometry data, capturing eural activity, dopamine release, and behavioral patterns. This approach enabled correlating locomotor behaviors with neural and dopamine dynamics. Movement was analyzed with DeepLabCut ^157^, where 1000 frames from all sessions were manually labeled to identify the tip of the tail, the base of the tail, the center of the body, the neck and snout. After applying DeepLabCut to the videos, we converted video coordinates in pixels to coordinates in centimeters through a transform using the dimensions of the arena. Optogenetic stimulation of Chrimson was delivered using a 593 nm laser at 40 Hz. The duration of the stimulation was 8 seconds, repeated for 10 trials with a randomized inter-trial interval of 1.5 min. Trajectory lengths for each mouse during pre-stimulation, stimulation and post-stimulation periods were calculated to determine overall displacement. The analysis compared these displacements to identify changes in movement patterns related to the optogenetic stimulation. Speed of movement was assessed across experimental phases and categorized into activity levels: immobile (<2 cm/sec), mobile (2-6 cm/sec), and running (>6 cm/sec). Time spent in each activity level was quantified across optogenetic manipulation phases: pre-, during, and post-stimulation.

### Behavioral motif analysis

Behavioral motifs were identified using B-SOiD^156^, a tool for clustering behavior into short-length movement motifs. For clustering, video recordings were preprocessed with DeepLabCut^157^, which annotated six body parts: snout, tail base, left front paw, right front paw, left hind paw, and right hind paw. We then calculated the average motif usage frequency (in frames per sec) for both the stimulation-on and stimulation-off periods. The stimulation-on period was defined as the time during which optical stimulation was delivered (8 sec per instance, ten times per session), whereas the stimulation-off period was determined from 8 sec before to the start of each stimulation. We evaluated changes in motif usage between these conditions using a two-tailed t-test and computed the ratio of motif usage between the stimulation-off and stimulation-on periods. Motif usage ratios were visualized using box plots generated with a Python library, displaying quartiles of the dataset and extending whiskers to show the distribution, excluding outliers determined by the 1.5 inter-quartile range rule. Motifs that were absent during sessions for certain mice were excluded from the analysis for those specific individuals, leading to some of the plots that do not include all the mice (n = 8 for Pnoc and 11 for Nts).

### Kinematic analysis

Kinematic data were calculated from the body parts coordination data, which were annotated by DeepLabCut. Movement speed was calculated using the center of body coordinate. Body twist angle was defined as the angle between the “tail base to center of body” and “center of body to neck” vectors. Similarly, head twist angle was measured between the “center of body to neck” and “neck to snout” vectors. We aggregated kinematic data from all mice in the Pnoc and Nts groups to plot the total distribution density of each variable. Histograms of angular velocities were made in 0.2 degree/sec bins from 0 to 20 degrees/sec (100 bins in total), smoothed over 5 bins to enhance visual clarity. Those for movement speed were made in 1 frame/sec bins from 0 to 1000 frames/sec (1000 bins in total), smoothed over 50 bins. All the histograms were then converted to distribution densities by normalizing the area under the curve to be 1. We calculated average head-twist and body-twist angular velocities, and movement speeds for each mouse, separating data according to motif. We then compared these kinematic variables between stimulation-off and stimulation-on periods to assess the effects of light stimulation on mouse movement kinetics, using a two-tailed t-test. The results were visually represented in box plots with swarm plots detailing the ratio of each kinematic variable between the stimulation-on and stimulation-off periods (n = 8 for Pnoc and 11 for Nts).

### T-maze probabilistic 2-choice switching task

This task is a modified version of the probabilistic 2-choice switching task^151^, implemented in a T-maze to spatially and temporally extend the action selection process, thereby facilitating the identification of action value encoding in neural signals. In this adapted version, mice were trained to choose between two options with reinforcement outcomes that intermittently swapped without warning. Notably, a 3-μl sucrose reward was introduced at the initiation port, and an air puff was administered in 5% of incorrect choices as a form of negative reinforcement. This setup aimed to encourage a “win-stay, lose-switch” strategy, requiring mice to adjust their choices in response to changing reinforcement outcomes. Specifically, one port had a 75% chance of providing a sucrose reward, whereas the other was non-reinforced. The reinforcement schedule for these ports was subject to random switches every 7-24 rewarded trials. Video data were collected and processed with Bonsai software, enabling real-time detection of corridor end zone entries. The system was automated through two Arduino microcontrollers managing the water spouts and interfacing with Bonsai via the Firmata protocol. High-resolution video frames from a FLIR-Oryx 10GigE camera were synchronized with the photometry setup, recording at 30 frames per sec. This setup facilitated the simultaneous recording of neuronal activity and dopamine release at a 10 Hz sampling rate across three channels: a reference (400-410 nm), green (460-490 nm), and red (555-570 nm). Behavior and action value were analyzed using a Q-learning model for each animal individually, assessing adjustments in decision-making strategies in response to dynamic changes in reinforcement.

### Action value estimation using Q-learning

Behavior and action values were analyzed using a Q-learning model for each mouse individually, assessing adjustments in decision-making strategies in response to dynamic changes in reinforcement. We used the Q-learning model to calculate the action values for different choices made during the task because it allowed us to update the estimated value of each option based on reinforcement outcomes. We used the following update rule for action values:

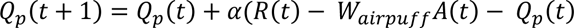

where Qp(t): Current action value of choosing port *p* at time t, *α*: Learning rate, which controls how quickly the values are updated, R(t): Outcome received at time t (1 for a reward, 0 for an omission, *W_airpuff_* : Penalty associated with receiving an airpuff, A(t): Binary indicator of whether an airpuff trial occurred. We then used a logistic function to calculate the probability of choosing the right port:

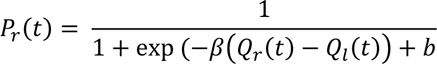

where *β* : parameter controlling the slope of the function and the explore-exploit trade-off, b: static bias toward one side.

We first estimated the parameters, Wairpuff, and b for each mouse individually by minimizing the negative log-likelihood:

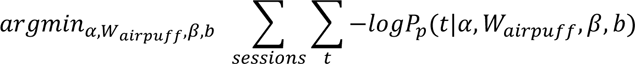

The parameters were optimized using the downhill simplex algorithm implemented through the scipy.optimize.fmin function in python. However, we observed a significant number of outliers in the Wairpuff values, likely due to the limited number of trials in each session. To address this, we recalculated the mean value of Wairpuff across sessions and used it as a default Wairpuff, subsequently only refitted the parameters, and b using adjusted mean of Wairpuff.

### Signal analysis

Signal data were analyzed using custom scripts developed in MATLAB. Five key events were identified for analysis: Trial Start, Initiation Port Exit, Turn, Choice Port Entry, and Choice Port Exit. For each event, we calculated the median inter-event intervals from successive pairs. These intervals were used to define a symmetric time window centered on each event, extending from half the median interval before the event to half the median interval after the event. These windows were then combined, allowing for a comprehensive visualization of all trials, accommodating variations in inter-event intervals. This approach facilitated a detailed examination of the temporal dynamics across different trial phases.

### Task bracketing index

The Task Bracketing Index was calculated by first isolating segments of each trial, spanning from initiation to choice. The index was then derived by measuring the mean signal value at the beginning and end segments of the trial—each defined as 25% of the total trial length—and subtracting the mean signal value of the central segment, which comprises the remaining 50% of the trial.

### Cross-correlation

For each trial, we compared the signal activity of one signal (e.g., neuronal signal Pnoc, Nts) against another signal (e.g., dopamine release) using the MATLAB crosscorr function. We computed the cross-correlation coefficient across various time lags, ranging from −1 sec to +1 sec. The cross-correlation coefficients were plotted against the lag values, with positive lags indicating that the first signal leads the second, and negative lags indicating that the second signal leads the first.

### CNN classifiers

We developed convolutional neural network (CNN) classifiers to categorize task space and choice space events. The task space was divided into five principal categories: Initiation, Left Choice, Left Turn, Right Choice and Right Turn. The choice space was divided into five categories: Airpuff, Left Omission, Left Rewarded, Right Omission and Right Rewarded. Each event category contained time windows from −2 sec to +2 sec relative to the event.

Given the time-series nature of the signal data, we chose a 1D CNN model. The dataset was partitioned into a training set (80% of the data) and a test set (20% of the data). To address class imbalance in the training set, class weights were computed using the compute_class_weight function from scikit-learn with the ’balanced’ option and then converted into a dictionary for compatibility with the Keras API. The CNN model was constructed using the Keras Sequential API and included the following layers: a 1D convolutional layer with 64 filters and ReLU activation, a max pooling layer with a pool size of 2, a dropout layer with a 0.3 dropout rate, a flatten layer, a dense layer with 64 units and L2 regularization (0.001), another dropout layer with a 0.3 dropout rate, and a softmax output layer for classification. The model was compiled using the Adam optimizer and the sparse categorical cross-entropy loss function, with accuracy as the performance metric. To mitigate overfitting and improve generalization, we used a ReduceLROnPlateau callback that monitored validation loss and reduced the learning rate by a factor of 0.1 if no improvement was seen for 50 epochs. The model was trained over a specified number of epochs with the validation set used for performance monitoring and early stopping. Class weights were incorporated to handle imbalance, and regularization techniques such as dropout and L2 regularization were used to prevent overfitting. The performance of the model was evaluated using accuracy metrics on the test set.

### LSTM classifier

We developed Long Short-Term Memory (LSTM) classifiers to categorize task space and choice space events, similar to our CNN approach. The LSTM model was constructed using the Keras Sequential API and included the following layers: an LSTM layer with 32 units and return_sequences=True, a dropout layer with a 0.3 dropout rate, another LSTM layer with 32 units and return_sequences=False, another dropout layer with a 0.3 dropout rate, a dense layer with 16 units and ReLU activation with L2 regularization (0.01), and a softmax output layer for classification. The model was compiled using the Adam optimizer and the sparse categorical cross-entropy loss function, with accuracy as the performance metric.

### Value classification

To investigate the neural encoding of action values, we categorized trials into three distinct groups based on the absolute values of the action values, using the first (Q1) and third (Q3) quartiles in each session as thresholds. Trials were classified as high-value when the absolute action values exceeded the Q3 threshold and as low-value when the absolute action values fell below the Q1 threshold. This classification was performed individually for each session to ensure an accurate reflection of the session-specific distribution of action values.

We used a total of 6 sec of data from each signal, spanning from −3 sec to +3 sec relative to the middle region of interest (ROI), as input. The data were then categorized into four classes: high-value left turn trials, high-value right turn trials, low-value left turn trials, and low-value right turn trials. For classification, we employed the same CNN and LSTM models described previously. The classification models were developed on a per-animal basis by merging all sessions from a single animal, as individual sessions contained insufficient data to train the classifiers effectively.

---

**Figure S1.**
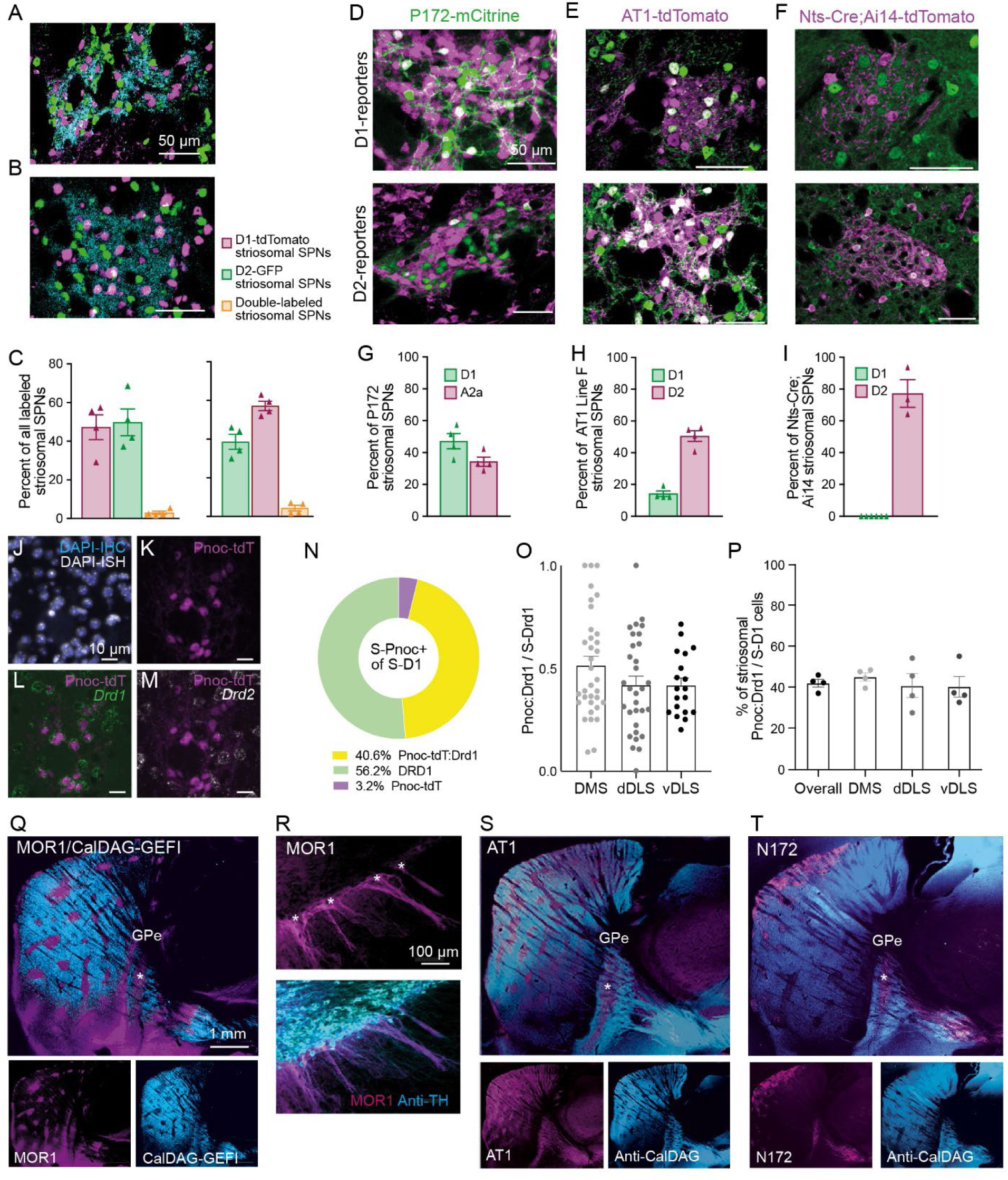
Characterization of striosomal lines: striosomal and matrix neurons target different regions of the globus pallidus, related to Figure 1 A-C. Striosomes contain similar proportions of neurons expressing the D1 or D2 dopamine receptor subtype. Striatal sections from D1-tdTomato;D2-GFP double-transgenic (A) and D1-GFP;A2a-Cre;Ai14-tdTomato triple-transgenic mice (B) were immunolabeled for tdTomato, GFP, and the striosome marker MOR1 (cyan). Within striosomes, counts for cells with the D1 and D2 markers were similar. The fluorescence for D1-tdTomato was stronger than for D1-GFP, and higher numbers of D1-type neurons were observed in (C left) than in (C right), consistent with previous reports^162^. Error bars show SEM. Two male and 2 female mice for each genotype. Three coronal sections (anterior, mid-level and caudal regions) were evaluated for each mouse. Neurons were counted in all the MOR1-identified striosomes that were dorsal to the anterior commissure. D-F. Striosome lines show differential reporter expression in D1 (top) and D2 (bottom) striosomal SPNs. Representative images of striosomes used to measure reporter expression in D1 and D2 striosomal SPNs are shown for mice labeled for P172-mCitrine striosomal SPNs (green, D), AT1-tdTomato soma-targeting line F (magenta, E) and Nts-Cre and Ai14-tdTomato striosomal SPNs (magenta, F). Double-labeled cells appear white. G-I. Plots showing the percent of SPNs positive for striosome marker that are double-labeled for D1 or D2/A2a reporters in mice for striosome marker P172-mCitrine (G), AT1-tdTomato (H) or Nts-Cre (I). For G and H, n = 4 mice for each double-transgenic genotype, balanced for sex. For I, n = 2 mice for Nts-Cre;Ai14;D1-GFP and n = 1 mouse for Nts-Cre;Ai14;D2-GFP. For each mouse, three coronal sections, taken from the anterior, mid-level and caudal regions, were evaluated. Counts were made only for cells within striosomes identified by MOR1 (for clarity, not shown). J. Sequential immunostaining (IHC) and *in situ* hybridization (ISH) protocols were performed on the same brain sections, and resulting images aligned according to DAPI-IHC (blue) and DAPI-ISH (white) signals. K. Striosomal Pnoc;Ai14-tdTomato neurons detected by clusters of striatal tdTomato-expressing cells. L and M. *Drd1* (L, green) and *Drd2* (M, white) signals were labeled by ISH. N. Striosomal Pnoc;Ai14 tdTomato reporter-expressing neurons labeled by ISH for *Drd1* accounted for 40% of all striosomal *Drd1* neurons. O. Variation in the proportion of striosomal *Drd1*-labeled neurons expressing Pnoc reporter, tdTomato, is shown across individual striosomes (dots) in the DMS, dorsal DLS (dDLS) and ventral DLS (vDLS). P. Percentage of cells averaged across mice (4 sections from 4 mice; 22, 19, 21, 23 striosomes per section; 500.5 ± 64 striosomal cells per section), shown as mean ± SEM. Q. Representative sagittal brain section from double-transgenic mice carrying MOR1-mCherry knock-in transgene (magenta) and the CalDAG-GEFI-GFP BAC (cyan). R. Coronal hemisections through the anterior SNpc of MOR1-mCherry line in immunolabeled for the striosome reporter (magenta, top) and the dopamine transporter (cyan, bottom) to identify dopaminergic cell bodies and descending dendrites. Asterisks designate the tops of striosome-dendron bouquets. S and T. Representative sagittal section from AT1-tdTomato (magenta, S) and P172-mCitrine (magenta, T) transgenic mouse brain immunolabeled for CalDAG-GEFI (cyan).

**Figure S2.**
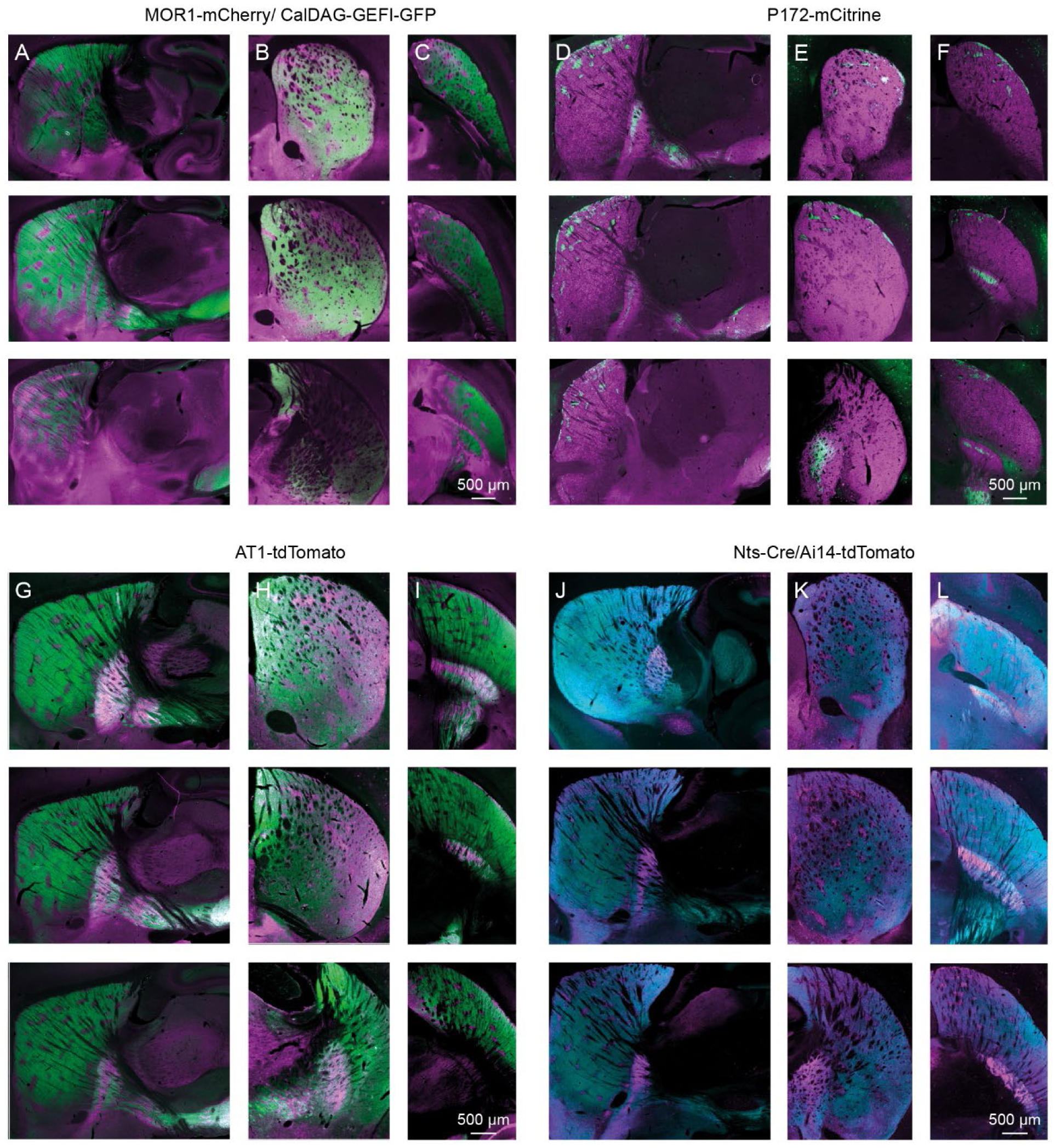
Projection pattern of striosomal lines, related to Figure 1 A-C. MOR1-mCherry/ CalDAG-GEFI-GFP. Images of sagittal (A), coronal (B) and para-horizontal (C) sections at three levels (from top to bottom, lateral to medial in A, anterior to posterior in B and dorsal to ventral in C) through the striatum of double-transgenic mice carrying MOR1-mCherry knock-in transgene (magenta) and the CalDAG-GEFI-GFP BAC, immunolabeled for mCherry (magenta) and GFP (green). D-F. Projection pattern of P172-mCitrine. Images of sagittal (D), coronal (E) and para-horizontal (F) sections at three striatal levels described in A-C from P172-mCitrine mice, immunolabeled for mCitrine (green) and CalDAG-GEFI (cyan). G-I. Projection pattern of AT1-tdTomato. Images of striatal sections described in A-C from double-transgenic mice carrying BACs for AT1-tdTomato axon-targeting line 14 and CalDAG-GEFI-GFP, immunolabeled for tdTomato (magenta) and GFP (green). J-L. Projection pattern of Nts-Cre/Ai14-tdTomato. Images of striatal sections described in A-C from double-transgenic mice carrying the Nts-Cre knock-in transgene and Cre-dependent Ai14-tdTomato (magenta), immunolabeled for mCherry (magenta) and CalDAG-GEFI (cyan).

**Figure S3.**
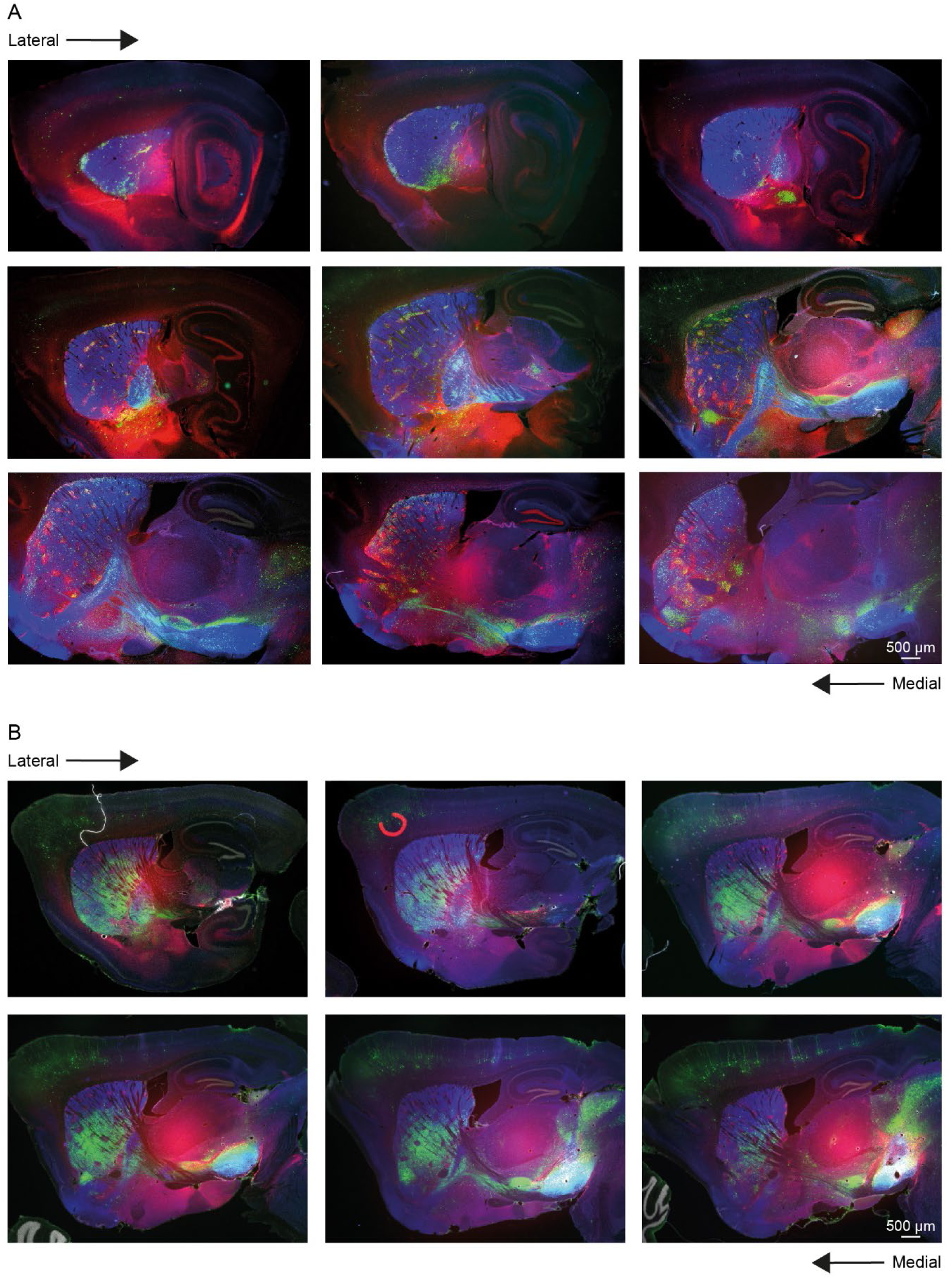
Images of series of sagittal sections from a PV-Cre and DAT-Cre mice with retrograde monosynaptic labeling of PV or dopamine cells in SNpr or SNpc, respectively, related to Figure 2. A. RV-mCherry-labeled neurons (green) counterstained for anti-MOR1 (striosome marker, red), anti-CalDAG-GEFI (matrix marker, blue) and DAPI (gray). B. RV-EYFP-labeled neurons (green) counterstained for anti-MOR1 (striosome marker, red), anti-CalDAG-GEFI (matrix marker, blue) and DAPI (gray).

**Figure S4.**
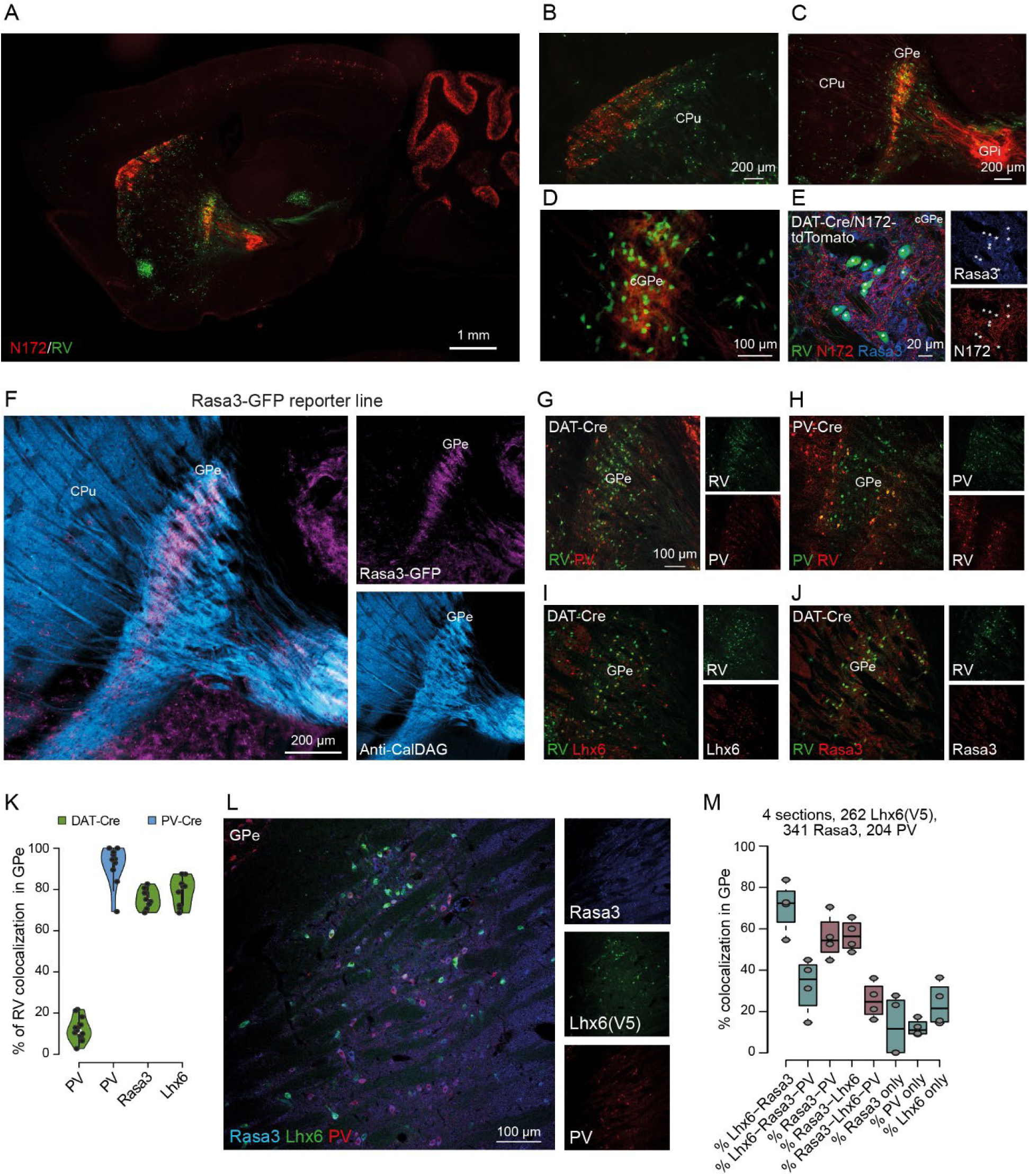
RV tracing in N172-tdTomato/DAT-Cre mice, related to Figure 3. A-D. Rabies-expressing SNpc-dopamine-targeting GPe neurons coinciding with striosomal D2/N172-labeled terminals in the cGPe zone. Sagittal sections from two N172-tdTomato/DAT-Cre transgenic mice that received injections of rabies-EGFP and Cre-dependent helper AAVs into the SNpc (n = 6). Retrogradely infected neurons as defined by co-labeling for the striosomal SPN markers P172-mCitrine (red). B shows enlargement of CPu. Retrogradely infected neurons are also strikingly enriched in the cGPe defined by striosomal SPN neuropil in red (C and D showing enlargement of GPe). E-M. Rasa3-GFP and central zone dopamine-projecting neuron characterization. E. Dopamine-projecting cGPe neurons, targeted by N172 striosomal terminals, are Rasa3 positive. F. Sagittal section from a Rasa3-GFP BAC transgenic mouse brain shows preferential distribution of GFP-positive cells (magenta) in the cGPe, relative to the pGPe that is targeted by matrix SPN axons immunolabeled for CalDAG-GEFI (cyan). G-J. Representative images of RV-labeled neurons and colocalization with PV (G), Lhx6 (I) and Rasa3 (J) in DAT-Cre tracing and colocalization with PV in PV-Cre tracing (H). K. Quantification of PV-, Rasa3- and Lhx6-positive RV-labeled neurons in the GPe. PV in DAT-Cre mice: 613 RV-labeled neurons in 12 sections from 3 mice. PV in PV-Cre mice: 291 RV-labeled neurons in 10 sections from 3 mice. Rasa3 in DAT-Cre mice: 1053 RV-labeled neurons in 10 sections from 3 mice. Lhx6 in DAT-Cre mice: 1029 RV-labeled neurons in 9 sections from 3 mice. L. Representative image of Lhx6-Cre mice injected with AAV-DIO-EYFP and immunolabeled for PV and Rasa3 (quantification in M). M. Quantification of PV, Rasa3 and Lhx6 colocalization in the GPe (see representative image in L).

**Figure S5.**
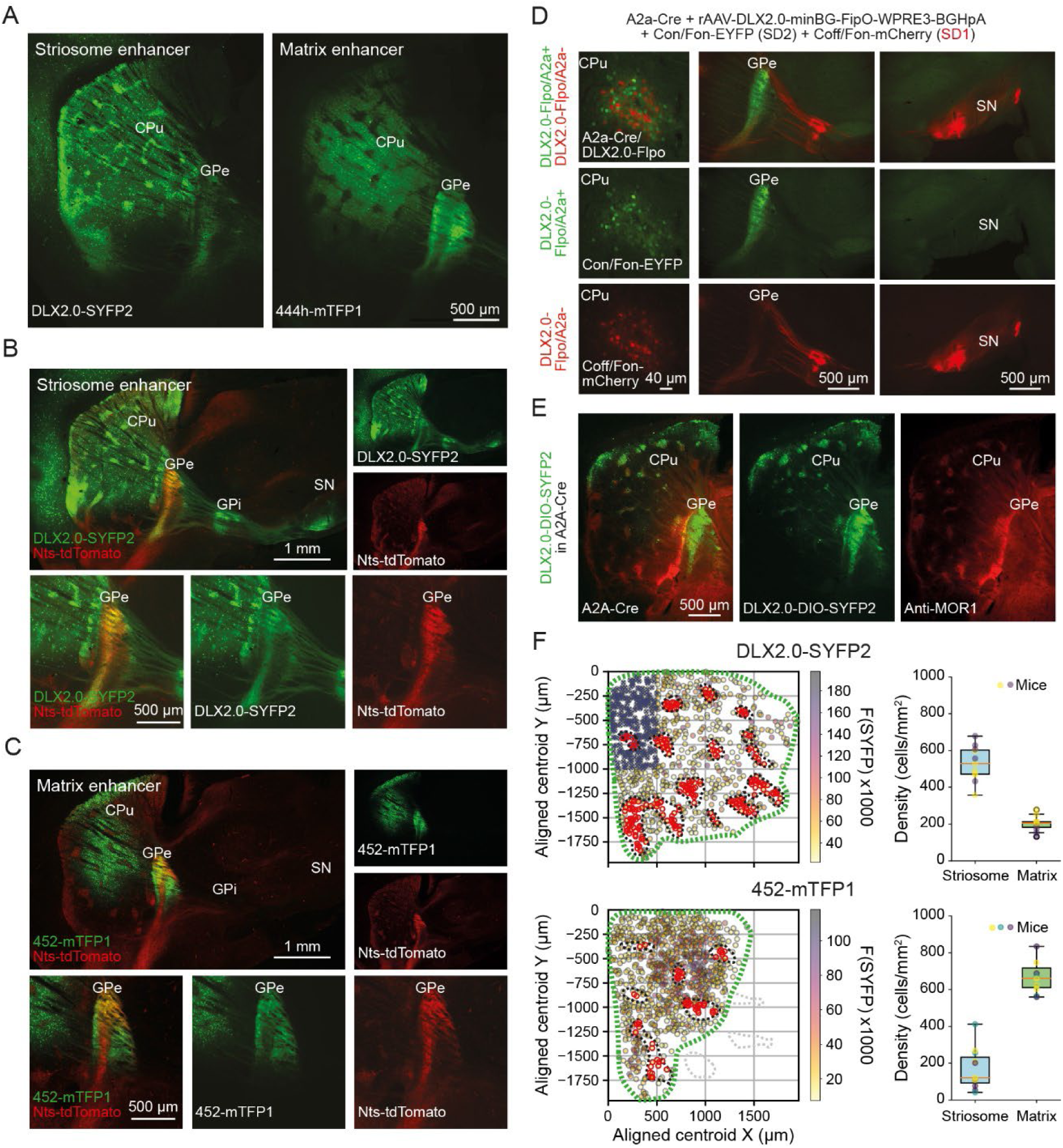
Striosome and matrix targeting with enhancer viruses, related to Figures 1-3. A. Sagittal section demonstrating the expression pattern of DLX2.0-SYFP2 (rAAV-DLX2.0-minBglobin-SYFP2-WPRE3-BGHpA, left) and rAAV-eHGT_444h-minBglobin-mTFP1-WPRE3-BGHpA (444h-mTFP1, right). Both viruses were injected with the same volume (500 nl) and at the same coordinates (AP = +0.86 mm, ML = 1.6 mm, and DV = 2.8 mm). Mice were perfused three weeks after injection. B. Comparison of striosome enhancer virus expression in Nts;tdTomato mice. Top panels show representative sagittal section depicting the expression patterns following the intrastriatal injection of the striosomal enhancer DLX2.0-SYFP2, as described in A. The left panel illustrates the overlay of DLX2.0-SYFP2 (green) and Nts;tdTomato (red) expression patterns, and right panels show separated channels for DLX2.0-SYFP2 (green) and Nts;tdTomato (red). Bottom Panels show, from left to right, enlarged views of the GPe with the combined fluorescence of both DLX2.0-SYFP2 (green) and Nts;tdTomato (red), followed by individual channels with DLX2.0-SYFP2 expression and Nts;tdTomato, respectively. C. Matrix enhancer virus expression in Nts;tdTomato mice. Top panels show sagittal sections from Nts;tdTomato mice, with the same injection parameters as those in A and B. These panels highlight the expression pattern of the matrix-specific enhancer virus rAAV-eHGT_452h-minBglobin-mTFP1-WPRE3-BGHpA (452h-mTFP1, green) in relation to the striosomal Nts;tdTomato SPNs. Bottom panels show sequence of images, from left to right, depicting the overlay of the matrix enhancer 452h-mTFP1 (green) and Nts;tdTomato (red), followed by the separated green and red channels. D. Targeted expression in D1 or D2 striosomal SPNs. Top left panel shows a representative image of specific targeting within A2a-Cre mice, achieved through an intrastriatal injection of AAV-DLX2.0-minBG-FIPO-WPRE3-BGHpA, followed by injection of intersectional viruses (Con/Fon-EYFP for striosomal D2 and Coff/Fon-mCherry for striosomal D1). This resulted in the selective labeling of striosomal D1-SPNs with mCherry (red) and striosomal D2-SPNs with EYFP (green) within the striatum. Top middle image depicts the axonal projections of both D1 and D2 striosomal neurons in the GPe, highlighting the dense terminal zones of S-D2 SPNs in the cGPe and the continuation of specifically the D1 striosomal neuron axons downstream to the GPe to the GPi, and substantia nigra (top right). Bottom panels show individual channels, which separately depict the red fluorescence of D1-SPNs and the green fluorescence of D2 SPNs. E. Specific targeting of striosomal D2 SPNs in A2a-Cre mice following an injection of the Cre-dependent striosomal enhancer virus DLX2.0-DIO-SYFP2 (AAV-DLX2.0-minBG-cDIO-SYFP2- WPRE-HGHpA). The overlay (left) illustrates the enhancer expression (green) and anti-MOR1 immunostaining (red) to confirm specificity to striosomes, with individual channels of green fluorescence marking the expression of the Cre-dependent enhancer virus (middle) and red immunostaining for MOR1 (right). F. Cell density analysis in striosomal (top) and matrix (bottom) regions. Left panels show 2D plot of the spatial coordinates of cells labeled with DLX2.0-SYFP2 (top) or 452h-SYFP (bottom) from a representative section. Cells are color-coded according to fluorescence intensity. Cells located within striosomal regions are highlighted in red, with boundaries delineated by black dotted lines based on MOR1 staining. Green dotted lines indicate the area used for calculating cell density. An excluded area in the dorsomedial part of the striatum is marked with ’X’ due to consistently decreased striosomal specificity in DLX2.0 expression in these striosomal area. Right panels show box plots representing the density of DLX2.0-SYFP2- or 452h-SYF-labeled cells in striosomal and matrix regions.

**Figure S6.**
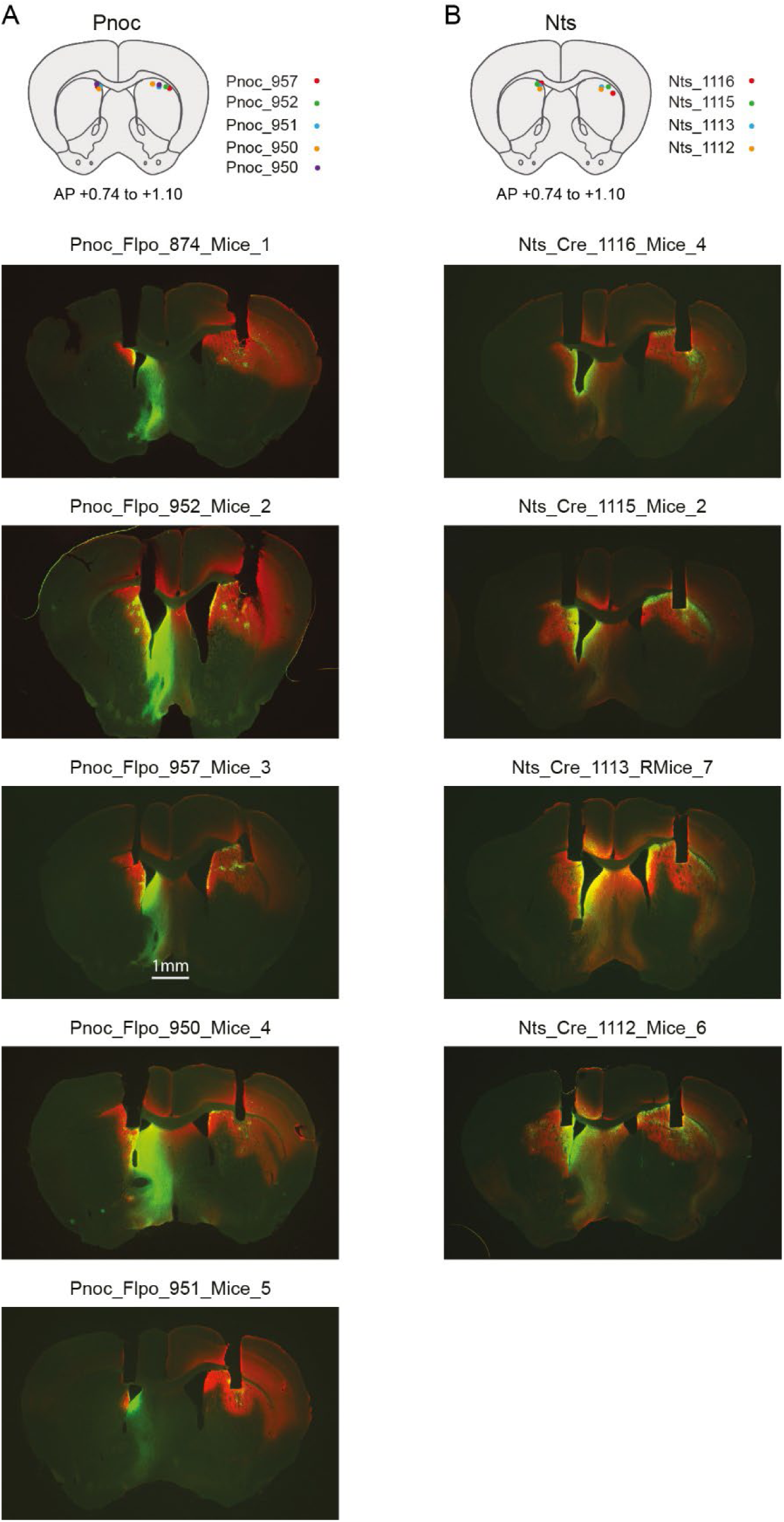
Injection sites and fiber placement for the 2-choice probabilistic switching maze task, related to Figure 6. Injection sites and fiber placements in Pnoc (A) and Nts (B) mice included in the analysis for the 2-choice probabilistic switching maze task presented in Figure 6.

**Figure S7.**
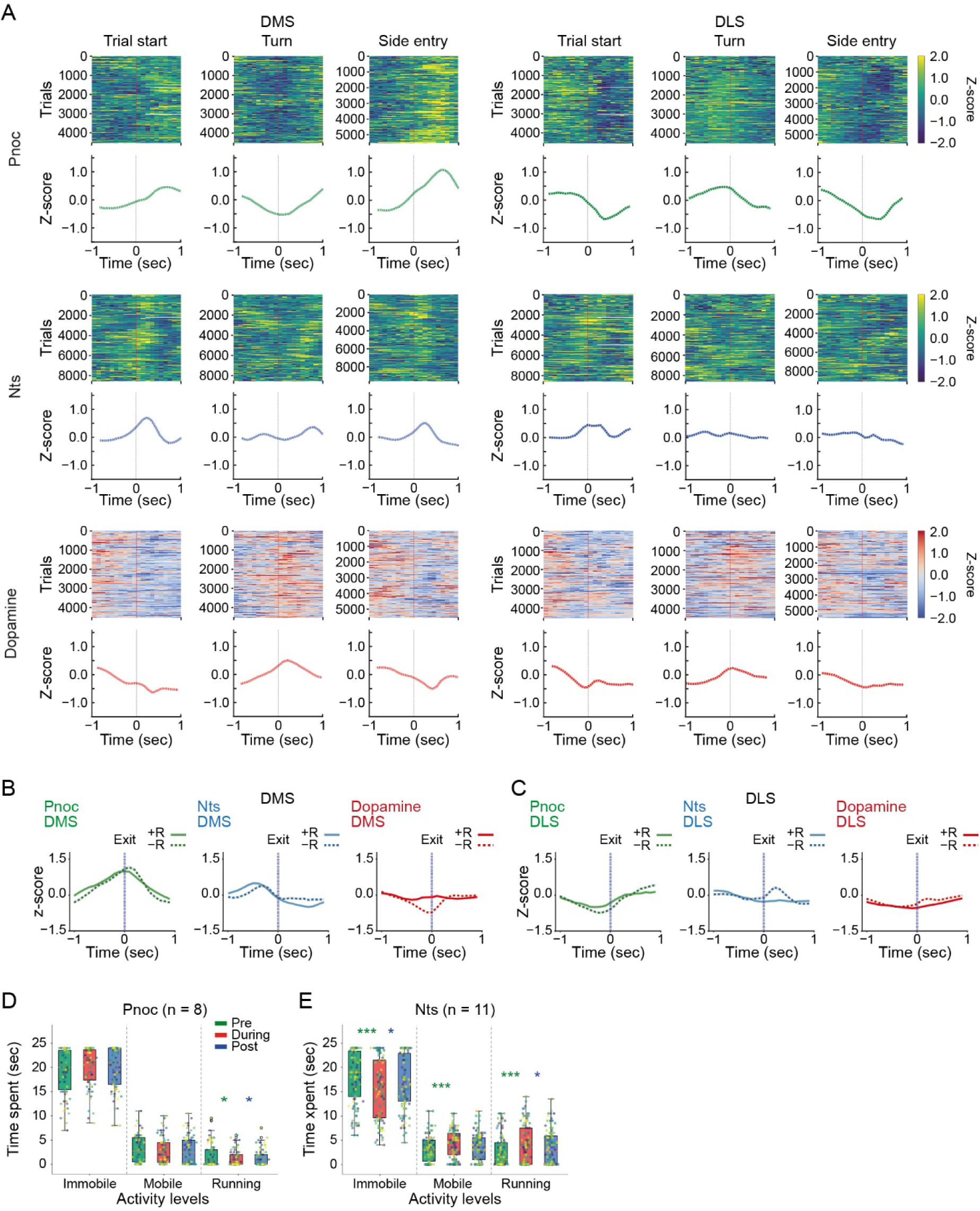
Temporal dynamics of Pnoc, Nts and dopamine activity, related to figure 6 and mobility score from optogenetic experiments, related to Figure 5. A. Temporal dynamics of Pnoc (top), Nts (middle) and dopamine (bottom) activity (average from all mice all sessions) recorded in the DMS (left) and DLS (right) during trials. Columns correspond to different behavioral events. B. Temporal dynamics of Pnoc, Nts and dopamine activity in DMS, aligned to the initiation, left and right ports exit for rewarded and unrewarded trials. Dopamine activity differentiates rewarded (+R) from unrewarded (−R) trials before the exit aligned to the reward delivery time (see Figure 6K). C. Same as B, but for activity in DLS. Nts activity deference between rewarded and unrewarded trials aligns well to the port exit but not to the outcome delivery time D and E. Time spent in immobile (<2 cm/sec), mobile (2-6 cm/sec) and running (>6 cm/sec) activity levels before, during and after the ontogenetic stimulation of S-D1 (D) and S-D2 (E) SPNs.

**Table S1.**
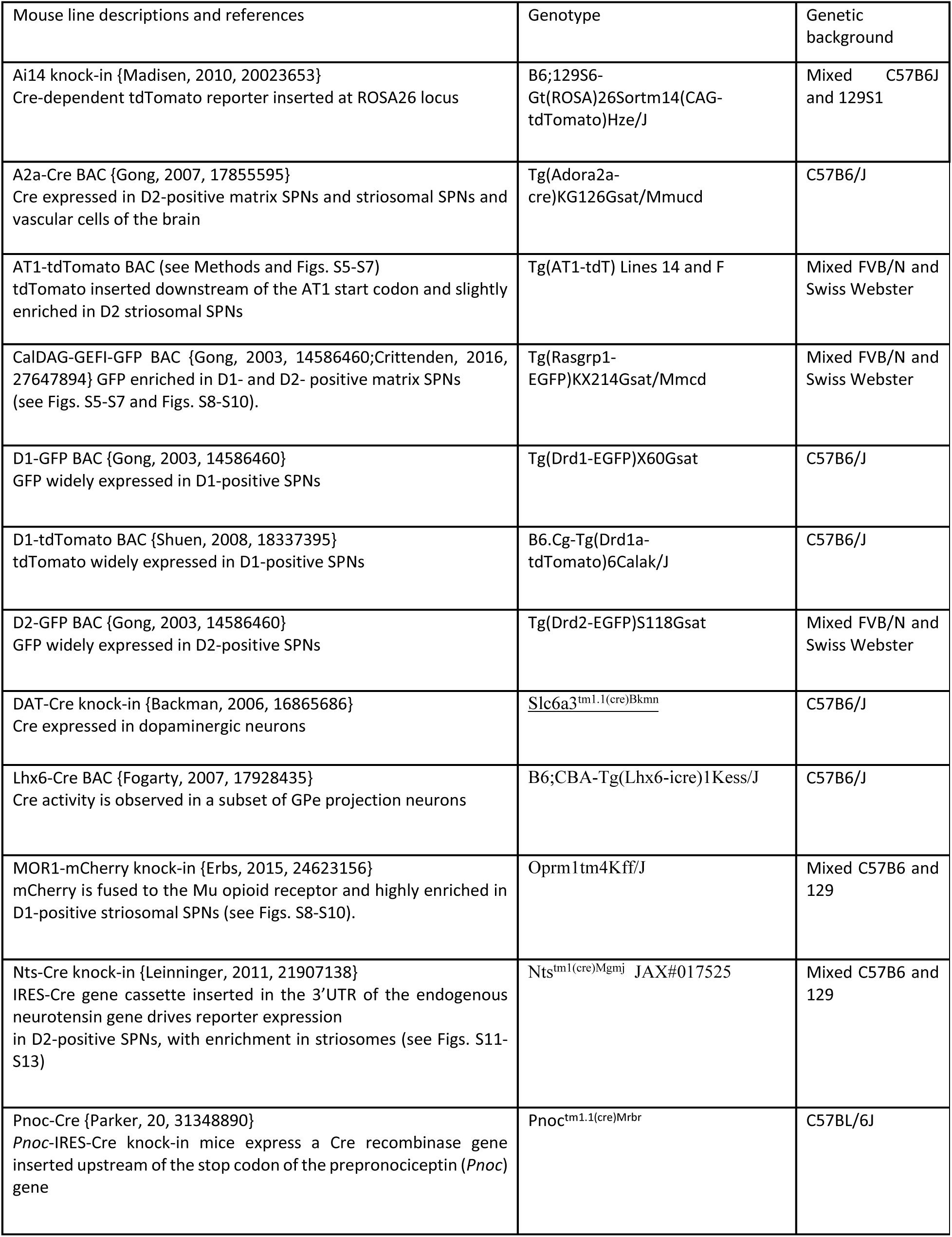
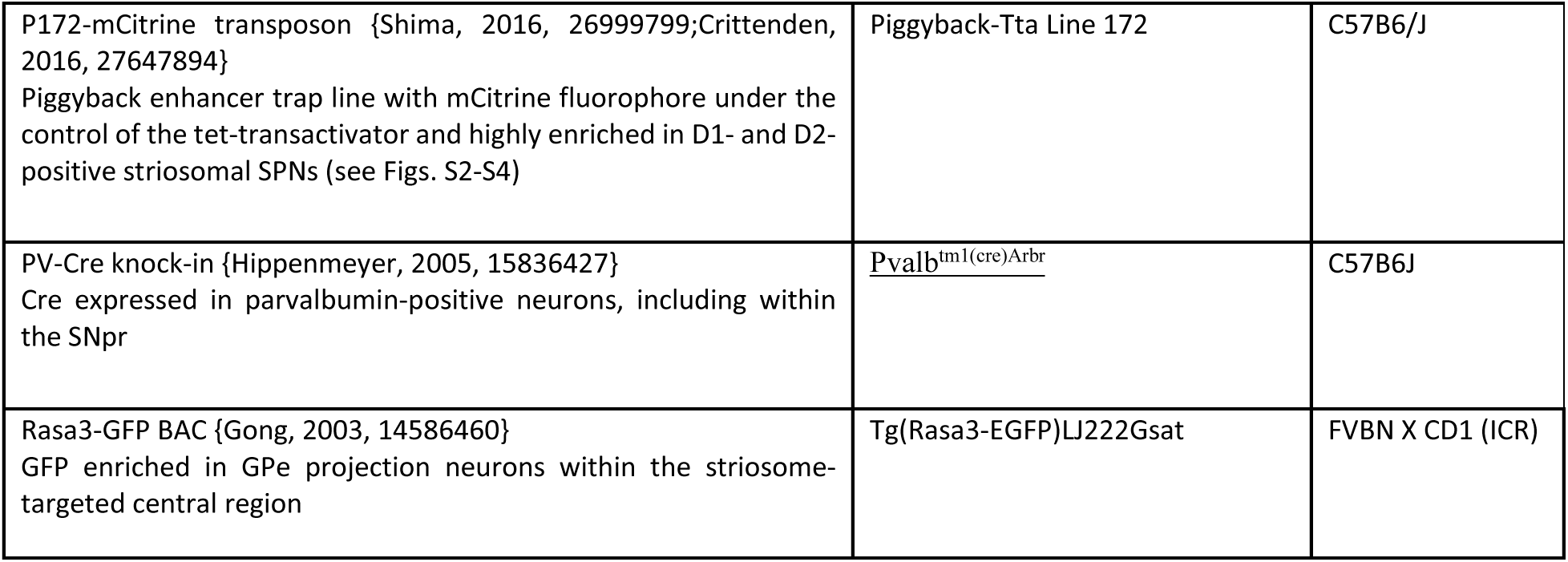
Transgenic mouse lies.

| Mouse line descriptions and references | Genotype | Genetic background |
| --- | --- | --- |
| Ai14 knock-in {Madisen, 2010, 20023653}<br>Cre-dependent tdTomato reporter inserted at ROSA26 locus | B6;129S6-Gt(ROSA)26Sortm14(CAG-tdTomato)Hze/J | Mixed C57B6J and 129S1 |
| A2a-Cre BAC {Gong, 2007, 17855595}<br>Cre expressed in D2-positive matrix SPNs and striosomal SPNs and vascular cells of the brain | Tg(Adora2a-cre)KG126Gsat/Mmucd | C57B6/J |
| AT1-tdTomato BAC (see Methods and Figs. S5-S7)<br>tdTomato inserted downstream of the AT1 start codon and slightly enriched in D2 striosomal SPNs | Tg(AT1-tdT) Lines 14 and F | Mixed FVB/N and Swiss Webster |
| CalDAG-GEFI-GFP BAC {Gong, 2003, 14586460;Crittenden, 2016, 27647894} GFP enriched in D1- and D2- positive matrix SPNs (see Figs. S5-S7 and Figs. S8-S10). | Tg(Rasgrp1-EGFP)KX214Gsat/Mmcd | Mixed FVB/N and Swiss Webster |
| D1-GFP BAC {Gong, 2003, 14586460}<br>GFP widely expressed in D1-positive SPNs | Tg(Drd1-EGFP)X60Gsat | C57B6/J |
| D1-tdTomato BAC {Shuen, 2008, 18337395}<br>tdTomato widely expressed in D1-positive SPNs | B6.Cg-Tg(Drd1a-tdTomato)6Calak/J | C57B6/J |
| D2-GFP BAC {Gong, 2003, 14586460}<br>GFP widely expressed in D2-positive SPNs | Tg(Drd2-EGFP)S118Gsat | Mixed FVB/N and Swiss Webster |
| DAT-Cre knock-in {Backman, 2006, 16865686}<br>Cre expressed in dopaminergic neurons | <u>Slc6a3<sup>tm1.1(cre)</sup>Bkmm</u> | C57B6/J |
| Lhx6-Cre BAC {Fogarty, 2007, 17928435}<br>Cre activity is observed in a subset of GPe projection neurons | B6;CBA-Tg(Lhx6-icre)1Kess/J | C57B6/J |
| MOR1-mCherry knock-in {Erbs, 2015, 24623156}<br>mCherry is fused to the Mu opioid receptor and highly enriched in D1-positive striosomal SPNs (see Figs. S8-S10). | Oprm1<sup>tm4Kff</sup>/J | Mixed C57B6 and 129 |
| Nts-Cre knock-in {Leininger, 2011, 21907138}<br>IRES-Cre gene cassette inserted in the 3'UTR of the endogenous neurotensin gene drives reporter expression in D2-positive SPNs, with enrichment in striosomes (see Figs. S11-S13) | Nts<sup>tm1(cre)</sup>Mgmj JAX#017525 | Mixed C57B6 and 129 |
| Pnoc-Cre {Parker, 20, 31348890}<br>Pnoc-IRES-Cre knock-in mice express a Cre recombinase gene inserted upstream of the stop codon of the prepronociceptin (*Pnoc*) gene | Pnoc<sup>tm1.1(cre)</sup>Mrbr | C57BL/6J |
| P172-mCitrine transposon {Shima, 2016, 26999799;Crittenden, 2016, 27647894}<br>Piggyback enhancer trap line with mCitrine fluorophore under the control of the tet-transactivator and highly enriched in D1- and D2-positive striosomal SPNs (see Figs. S2-S4) | Piggyback-Tta Line 172 | C57B6/J |
| PV-Cre knock-in {Hippenmeyer, 2005, 15836427}<br>Cre expressed in parvalbumin-positive neurons, including within the SNpr | <u>Pvalb</u><sup>tm1(cre)Arbr</sup> | C57B6J |
| Rasa3-GFP BAC {Gong, 2003, 14586460}<br>GFP enriched in GPe projection neurons within the striosome-targeted central region | Tg(Rasa3-EGFP)LJ222Gsat | FVBN X CD1 (ICR) |

**Table S2:**
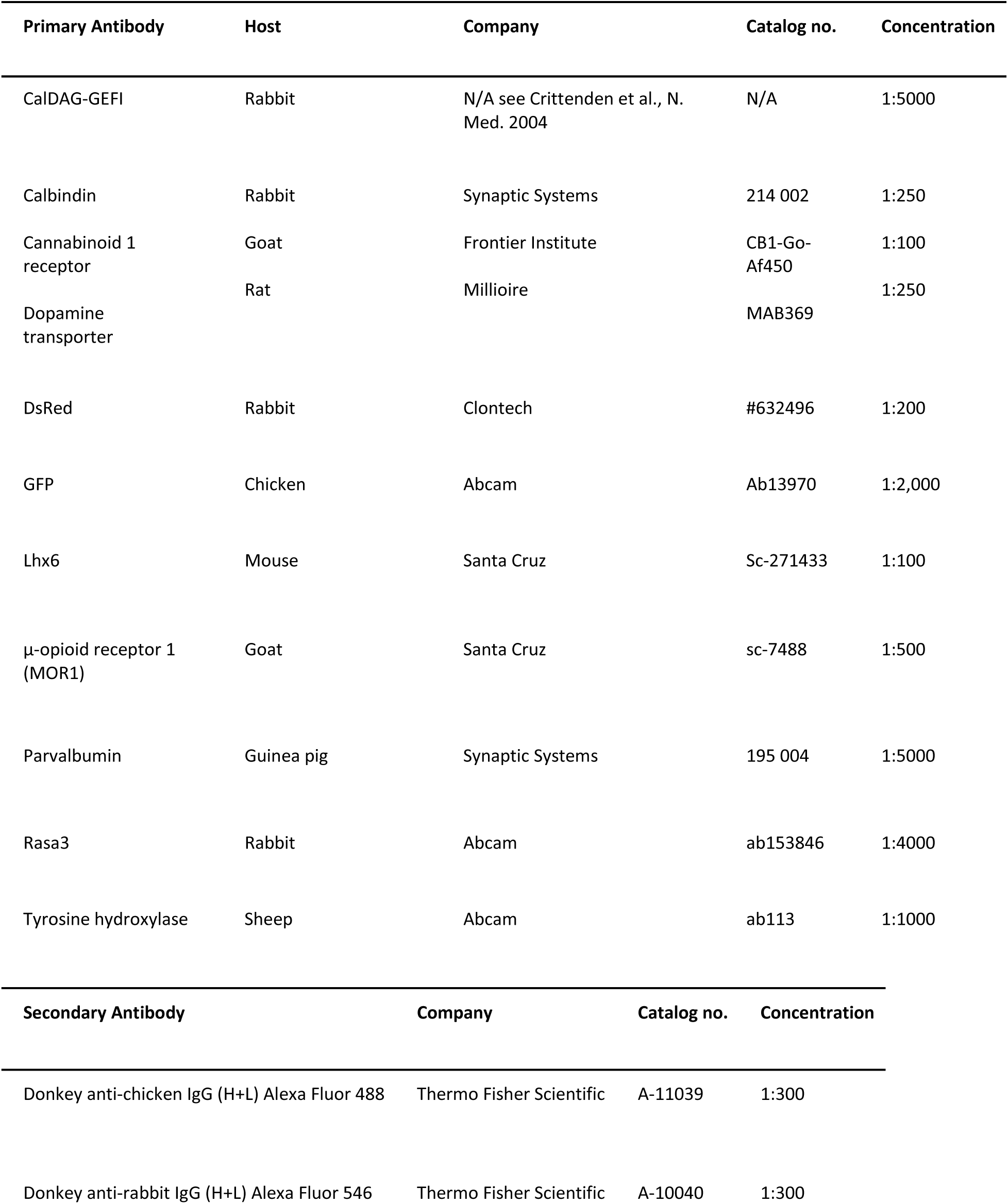
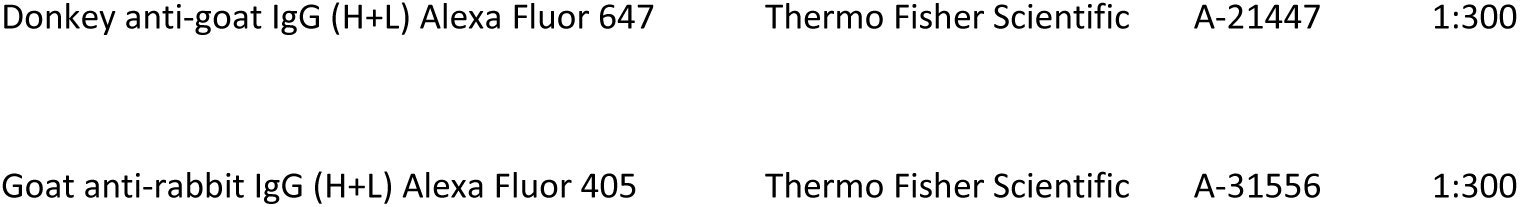
Antibodies.

| Primary Antibody | Host | Company | Catalog no. | Concentration |
| --- | --- | --- | --- | --- |
| CalDAG-GEFI | Rabbit | N/A see Crittenden et al., N. Med. 2004 | N/A | 1:5000 |
| Calbindin | Rabbit | Synaptic Systems | 214 002 | 1:250 |
| Cannabinoid 1 receptor | Goat | Frontier Institute | CB1-Go-Af450 | 1:100 |
| Dopamine transporter | Rat | Millioire | MAB369 | 1:250 |
| DsRed | Rabbit | Clontech | #632496 | 1:200 |
| GFP | Chicken | Abcam | Ab13970 | 1:2,000 |
| Lhx6 | Mouse | Santa Cruz | Sc-271433 | 1:100 |
| μ-opioid receptor 1 (MOR1) | Goat | Santa Cruz | sc-7488 | 1:500 |
| Parvalbumin | Guinea pig | Synaptic Systems | 195 004 | 1:5000 |
| Rasa3 | Rabbit | Abcam | ab153846 | 1:4000 |
| Tyrosine hydroxylase | Sheep | Abcam | ab113 | 1:1000 |
| Secondary Antibody | Company |  | Catalog no. | Concentration |
| Donkey anti-chicken IgG (H+L) Alexa Fluor 488 | Thermo Fisher Scientific |  | A-11039 | 1:300 |
| Donkey anti-rabbit IgG (H+L) Alexa Fluor 546 | Thermo Fisher Scientific |  | A-10040 | 1:300 |

|  |  |  |  |
| --- | --- | --- | --- |
| Donkey anti-goat IgG (H+L) Alexa Fluor 647 | Thermo Fisher Scientific | A-21447 | 1:300 |
| Goat anti-rabbit IgG (H+L) Alexa Fluor 405 | Thermo Fisher Scientific | A-31556 | 1:300 |

## REFERENCES

1. Albin, R.L., Young, A.B., and Penney, J.B. (1989). The functional anatomy of basal ganglia disorders. Trends Neurosci. 12, 366–375.

2. DeLong, M.R. (1990). Primate models of movement disorders of basal ganglia origin. Trends Neurosci. 13, 281–285.

3. Crossman, A.R. (1989). Neural mechanisms in disorders of movement. Comp. Biochem. Physiol. A Comp. Physiol. 93, 141–149.

4. Bateup, H.S., Santini, E., Shen, W., Birnbaum, S., Valjent, E., Surmeier, D.J., Fisone, G., Nestler, E.J., and Greengard, P. (2010). Distinct subclasses of medium spiny neurons differentially regulate striatal motor behaviors. Proc. Natl. Acad. Sci. U. S. A. 107, 14845– 14850.

5. Alexander, G.E., and Crutcher, M.D. (1990). Functional architecture of basal ganglia circuits: neural substrates of parallel processing. Trends Neurosci. 13, 266–271.

6. Mink, J.W. (2003). The Basal Ganglia and involuntary movements: impaired inhibition of competing motor patterns. Arch. Neurol. 60, 1365–1368.

7. Kawaguchi, Y., Wilson, C.J., and Emson, P.C. (1990). Projection subtypes of rat neostriatal matrix cells revealed by intracellular injection of biocytin. J. Neurosci. 10, 3421– 3438.

8. Oldenburg, I.A., and Sabatini, B.L. (2015). Antagonistic but Not Symmetric Regulation of Primary Motor Cortex by Basal Ganglia Direct and Indirect Pathways. Neuron 86, 1174– 1181.

9. Nishi, A., Snyder, G.L., and Greengard, P. (1997). Bidirectional regulation of DARPP-32 phosphorylation by dopamine. J. Neurosci. 17, 8147–8155.

10. Greengard, P., Allen, P.B., and Nairn, A.C. (1999). Beyond the dopamine receptor: the DARPP-32/protein phosphatase-1 cascade. Neuron 23, 435–447.

11. Gerfen, C.R., Engber, T.M., Mahan, L.C., Susel, Z., Chase, T.N., Monsma, F.J., Jr, and Sibley, D.R. (1990). D1 and D2 dopamine receptor-regulated gene expression of striatonigral and striatopallidal neurons. Science 250, 1429–1432.

12. Mink, J.W., and Thach, W.T. (1991). Basal ganglia motor control. III. Pallidal ablation: normal reaction time, muscle cocontraction, and slow movement. J. Neurophysiol. 65, 330–351.

13. Mink, J.W. (1996). The basal ganglia: focused selection and inhibition of competing motor programs. Prog. Neurobiol. 50, 381–425.

14. Nambu, A., Tokuno, H., Hamada, I., Kita, H., Imanishi, M., Akazawa, T., Ikeuchi, Y., and Hasegawa, N. (2000). Excitatory cortical inputs to pallidal neurons via the subthalamic nucleus in the monkey. J. Neurophysiol. 84, 289–300.

15. Hikosaka, O., Takikawa, Y., and Kawagoe, R. (2000). Role of the basal ganglia in the control of purposive saccadic eye movements. Physiol. Rev. 80, 953–978.

16. Vicente, A.M., Galvão-Ferreira, P., Tecuapetla, F., and Costa, R.M. (2016). Direct and indirect dorsolateral striatum pathways reinforce different action strategies. Curr. Biol. 26, R267–R269.

17. Isomura, Y., Takekawa, T., Harukuni, R., Handa, T., Aizawa, H., Takada, M., and Fukai, T. (2013). Reward-modulated motor information in identified striatum neurons. J. Neurosci. 33, 10209–10220.

18. Cui, G., Jun, S.B., Jin, X., Pham, M.D., Vogel, S.S., Lovinger, D.M., and Costa, R.M. (2013). Concurrent activation of striatal direct and indirect pathways during action initiation. Nature 494, 238–242.

19. Surmeier, D.J., Mercer, J.N., and Chan, C.S. (2005). Autonomous pacemakers in the basal ganglia: who needs excitatory synapses anyway? Curr. Opin. Neurobiol. 15, 312– 318.

20. Gatev, P., Darbin, O., and Wichmann, T. (2006). Oscillations in the basal ganglia under normal conditions and in movement disorders. Mov. Disord. 21, 1566–1577.

21. Okamoto, S., Sohn, J., Tanaka, T., Takahashi, M., Ishida, Y., Yamauchi, K., Koike, M., Fujiyama, F., and Hioki, H. (2020). Overlapping Projections of Neighboring Direct and Indirect Pathway Neostriatal Neurons to Globus Pallidus External Segment. iScience 23, 101409.

22. Parent, A., Sato, F., Wu, Y., Gauthier, J., Lévesque, M., and Parent, M. (2000). Organization of the basal ganglia: the importance of axonal collateralization. Trends Neurosci. 23, S20–S27.

23. Graybiel, A.M., and Ragsdale, C.W., Jr (1978). Histochemically distinct compartments in the striatum of human, monkeys, and cat demonstrated by acetylthiocholinesterase staining. Proc. Natl. Acad. Sci. U. S. A. 75, 5723–5726.

24. Wallace, M.L., Saunders, A., Huang, K.W., Philson, A.C., Goldman, M., Macosko, E.Z., McCarroll, S.A., and Sabatini, B.L. (2017). Genetically Distinct Parallel Pathways in the Entopeduncular Nucleus for Limbic and Sensorimotor Output of the Basal Ganglia. Neuron 94, 138–152.e5.

25. Courtney, C.D., Pamukcu, A., and Chan, C.S. (2023). Cell and circuit complexity of the external globus pallidus. Nat. Neurosci. 26, 1147–1159.

26. Mallet, N., Micklem, B.R., Henny, P., Brown, M.T., Williams, C., Bolam, J.P., Nakamura, K.C., and Magill, P.J. (2012). Dichotomous organization of the external globus pallidus. Neuron 74, 1075–1086.

27. Lilascharoen, V., Wang, E.H.-J., Do, N., Pate, S.C., Tran, A.N., Yoon, C.D., Choi, J.-H., Wang, X.-Y., Pribiag, H., Park, Y.-G., et al. (2021). Divergent pallidal pathways underlying distinct Parkinsonian behavioral deficits. Nat. Neurosci. 24, 504–515.

28. Mastro, K.J., Bouchard, R.S., Holt, H.A.K., and Gittis, A.H. (2014). Transgenic mouse lines subdivide external segment of the globus pallidus (GPe) neurons and reveal distinct GPe output pathways. J. Neurosci. 34, 2087–2099.

29. Dong, J., Hawes, S., Wu, J., Le, W., and Cai, H. (2021). Connectivity and Functionality of the Globus Pallidus Externa Under Normal Conditions and Parkinson’s Disease. Front. Neural Circuits 15, 645287.

30. Wickersham, I.R., Lyon, D.C., Barnard, R.J.O., Mori, T., Finke, S., Conzelmann, K.-K., Young, J.A.T., and Callaway, E.M. (2007). Monosynaptic restriction of transsynaptic tracing from single, genetically targeted neurons. Neuron 53, 639–647.

31. Ährlund-Richter, S., Xuan, Y., van Lunteren, J.A., Kim, H., Ortiz, C., Pollak Dorocic, I., Meletis, K., and Carlén, M. (2019). A whole-brain atlas of monosynaptic input targeting four different cell types in the medial prefrontal cortex of the mouse. Nat. Neurosci. 22, 657–668.

32. Lazaridis, I., Tzortzi, O., Weglage, M., Märtin, A., Xuan, Y., Parent, M., Johansson, Y., Fuzik, J., Fürth, D., Fenno, L.E., et al. (2019). A hypothalamus-habenula circuit controls aversion. Mol. Psychiatry 24, 1351–1368.

33. Kita, H., and Kitai, S.T. (1994). The morphology of globus pallidus projection neurons in the rat: an intracellular staining study. Brain Res. 636, 308–319.

34. Fujiyama, F., Nakano, T., Matsuda, W., Furuta, T., Udagawa, J., and Kaneko, T. (2016). A single-neuron tracing study of arkypallidal and prototypic neurons in healthy rats. Brain Struct. Funct. 221, 4733–4740.

35. Oh, Y.-M., Karube, F., Takahashi, S., Kobayashi, K., Takada, M., Uchigashima, M., Watanabe, M., Nishizawa, K., Kobayashi, K., and Fujiyama, F. (2017). Using a novel PV-Cre rat model to characterize pallidonigral cells and their terminations. Brain Struct. Funct. 222, 2359–2378.

36. Celada, P., Paladini, C.A., and Tepper, J.M. (1999). GABAergic control of rat substantia nigra dopaminergic neurons: role of globus pallidus and substantia nigra pars reticulata. Neuroscience 89, 813–825.

37. Chang, H.T., Wilson, C.J., and Kitai, S.T. (1981). Single neostriatal efferent axons in the globus pallidus: a light and electron microscopic study. Science 213, 915–918.

38. Wu, Y., Richard, S., and Parent, A. (2000). The organization of the striatal output system: a single-cell juxtacellular labeling study in the rat. Neurosci. Res. 38, 49–62.

39. Lévesque, M., and Parent, A. (2005). The striatofugal fiber system in primates: a reevaluation of its organization based on single-axon tracing studies. Proc. Natl. Acad. Sci. U. S. A. 102, 11888–11893.

40. Fujiyama, F., Sohn, J., Nakano, T., Furuta, T., Nakamura, K.C., Matsuda, W., and Kaneko, T. (2011). Exclusive and common targets of neostriatofugal projections of rat striosome neurons: a single neuron-tracing study using a viral vector. Eur. J. Neurosci. 33, 668–677.

41. Labouesse, M.A., Torres-Herraez, A., Chohan, M.O., Villarin, J.M., Greenwald, J., Sun, X., Zahran, M., Tang, A., Lam, S., Veenstra-VanderWeele, J., et al. (2023). A non-canonical striatopallidal Go pathway that supports motor control. Nat. Commun. 14, 6712.

42. Crittenden, J.R., Tillberg, P.W., Riad, M.H., Shima, Y., Gerfen, C.R., Curry, J., Housman, D.E., Nelson, S.B., Boyden, E.S., and Graybiel, A.M. (2016). Striosome-dendron bouquets highlight a unique striatonigral circuit targeting dopamine-containing neurons. Proc. Natl. Acad. Sci. U. S. A. 113, 11318–11323.

43. Gong, S., Zheng, C., Doughty, M.L., Losos, K., Didkovsky, N., Schambra, U.B., Nowak, N.J., Joyner, A., Leblanc, G., Hatten, M.E., et al. (2003). A gene expression atlas of the central nervous system based on bacterial artificial chromosomes. Nature 425, 917–925.

44. Shima, Y., Sugino, K., Hempel, C.M., Shima, M., Taneja, P., Bullis, J.B., Mehta, S., Lois, C., and Nelson, S.B. (2016). A Mammalian enhancer trap resource for discovering and manipulating neuronal cell types. Elife 5, e13503.

45. Erbs, E., Faget, L., Scherrer, G., Matifas, A., Filliol, D., Vonesch, J.-L., Koch, M., Kessler, P., Hentsch, D., Birling, M.-C., et al. (2015). A mu-delta opioid receptor brain atlas reveals neuronal co-occurrence in subcortical networks. Brain Struct. Funct. 220, 677–702.

46. Parker, K.E., Pedersen, C.E., Gomez, A.M., Spangler, S.M., Walicki, M.C., Feng, S.Y., Stewart, S.L., Otis, J.M., Al-Hasani, R., McCall, J.G., et al. (2019). A Paranigral VTA Nociceptin Circuit that Constrains Motivation for Reward. Cell 178, 653–671.e19.

47. Leinninger, G.M., Opland, D.M., Jo, Y.-H., Faouzi, M., Christensen, L., Cappellucci, L.A., Rhodes, C.J., Gnegy, M.E., Becker, J.B., Pothos, E.N., et al. (2011). Leptin action via neurotensin neurons controls orexin, the mesolimbic dopamine system and energy balance. Cell Metab. 14, 313–323.

48. Madisen, L., Zwingman, T.A., Sunkin, S.M., Oh, S.W., Zariwala, H.A., Gu, H., Ng, L.L., Palmiter, R.D., Hawrylycz, M.J., Jones, A.R., et al. (2010). A robust and high-throughput Cre reporting and characterization system for the whole mouse brain. Nat. Neurosci. 13, 133–140.

49. Kawasaki, H., Springett, G.M., Toki, S., Canales, J.J., Harlan, P., Blumenstiel, J.P., Chen, E.J., Bany, I.A., Mochizuki, N., Ashbacher, A., et al. (1998). A Rap guanine nucleotide exchange factor enriched highly in the basal ganglia. Proc. Natl. Acad. Sci. U. S. A. 95, 13278–13283.

50. Crittenden, J.R., Dunn, D.E., Merali, F.I., Woodman, B., Yim, M., Borkowska, A.E., Frosch, M.P., Bates, G.P., Housman, D.E., Lo, D.C., et al. (2010). CalDAG-GEFI down-regulation in the striatum as a neuroprotective change in Huntington’s disease. Hum. Mol. Genet. 19, 1756–1765.

51. Gerfen, C.R., Paletzki, R., and Heintz, N. (2013). GENSAT BAC cre-recombinase driver lines to study the functional organization of cerebral cortical and basal ganglia circuits. Neuron 80, 1368–1383.

52. Liu, F.C., and Graybiel, A.M. (1992). Heterogeneous development of calbindin-D28K expression in the striatal matrix. J. Comp. Neurol. 320, 304–322.

53. Sgobio, C., Wu, J., Zheng, W., Chen, X., Pan, J., Salinas, A.G., Davis, M.I., Lovinger, D.M., and Cai, H. (2017). Aldehyde dehydrogenase 1-positive nigrostriatal dopaminergic fibers exhibit distinct projection pattern and dopamine release dynamics at mouse dorsal striatum. Sci. Rep. 7, 5283.

54. Azcorra, M., Gaertner, Z., Davidson, C., He, Q., Kim, H., Nagappan, S., Hayes, C.K., Ramakrishnan, C., Fenno, L., Kim, Y.S., et al. (2023). Unique functional responses differentially map onto genetic subtypes of dopamine neurons. Nat. Neurosci. 26, 1762– 1774.

55. Kita, H., and Kita, T. (2001). Number, origins, and chemical types of rat pallidostriatal projection neurons. J. Comp. Neurol. 437, 438–448.

56. Miyamoto, Y., Katayama, S., Shigematsu, N., Nishi, A., and Fukuda, T. (2018). Striosome-based map of the mouse striatum that is conformable to both cortical afferent topography and uneven distributions of dopamine D1 and D2 receptor-expressing cells. Brain Struct. Funct. 223, 4275–4291.

57. Smith, Y., and Bolam, J.P. (1989). Neurons of the substantia nigra reticulata receive a dense GABA-containing input from the globus pallidus in the rat. Brain Res. 493, 160–167.

58. Connelly, W.M., Schulz, J.M., Lees, G., and Reynolds, J.N.J. (2010). Differential short-term plasticity at convergent inhibitory synapses to the substantia nigra pars reticulata. J. Neurosci. 30, 14854–14861.

59. Hegeman, D.J., Hong, E.S., Hernández, V.M., and Chan, C.S. (2016). The external globus pallidus: progress and perspectives. Eur. J. Neurosci. 43, 1239–1265.

60. Saunders, A., Macosko, E.Z., Wysoker, A., Goldman, M., Krienen, F.M., de Rivera, H., Bien, E., Baum, M., Bortolin, L., Wang, S., et al. (2018). Molecular Diversity and Specializations among the Cells of the Adult Mouse Brain. Cell 174, 1015–1030.e16.

61. Wu, B., Zhang, S., Guo, Z., Wang, G., Zhang, G., Xie, L., Lou, J., Chen, X., Wu, D., Bergmeier, W., et al. (2018). RAS P21 Protein Activator 3 (RASA3) Specifically Promotes Pathogenic T Helper 17 Cell Generation by Repressing T-Helper-2-Cell-Biased Programs. Immunity 49, 886–898.e5.

62. Johansen, K.H., Golec, D.P., Okkenhaug, K., and Schwartzberg, P.L. (2023). Mind the GAP: RASA2 and RASA3 GTPase-activating proteins as gatekeepers of T cell activation and adhesion. Trends Immunol. 44, 917–931.

63. Johansen, K.H., Golec, D.P., Huang, B., Park, C., Thomsen, J.H., Preite, S., Cannons, J.L., Garçon, F., Schrom, E.C., Courrèges, C.J.F., et al. (2022). A CRISPR screen targeting PI3K effectors identifies RASA3 as a negative regulator of LFA-1-mediated adhesion in T cells. Sci. Signal. 15, eabl9169.

64. Kubota, Y., Liu, J., Hu, D., DeCoteau, W.E., Eden, U.T., Smith, A.C., and Graybiel, A.M. (2009). Stable encoding of task structure coexists with flexible coding of task events in sensorimotor striatum. J. Neurophysiol. 102, 2142–2160.

65. Friedman, A., Hueske, E., Drammis, S.M., Toro Arana, S.E., Nelson, E.D., Carter, C.W., Delcasso, S., Rodriguez, R.X., Lutwak, H., DiMarco, K.S., et al. (2020). Striosomes Mediate Value-Based Learning Vulnerable in Age and a Huntington’s Disease Model. Cell 183, 918–934.e49.

66. Amemori, S., Graybiel, A.M., and Amemori, K.-I. (2021). Causal Evidence for Induction of Pessimistic Decision-Making in Primates by the Network of Frontal Cortex and Striosomes. Front. Neurosci. 15, 649167.

67. Amemori, S., Amemori, K.-I., Yoshida, T., Papageorgiou, G.K., Xu, R., Shimazu, H., Desimone, R., and Graybiel, A.M. (2020). Microstimulation of primate neocortex targeting striosomes induces negative decision-making. Eur. J. Neurosci. 51, 731–741.

68. Friedman, A., Homma, D., Bloem, B., Gibb, L.G., Amemori, K.-I., Hu, D., Delcasso, S., Truong, T.F., Yang, J., Hood, A.S., et al. (2017). Chronic Stress Alters Striosome-Circuit Dynamics, Leading to Aberrant Decision-Making. Cell 171, 1191–1205.e28.

69. Friedman, A., Homma, D., Gibb, L.G., Amemori, K.-I., Rubin, S.J., Hood, A.S., Riad, M.H., and Graybiel, A.M. (2015). A Corticostriatal Path Targeting Striosomes Controls Decision-Making under Conflict. Cell 161, 1320–1333.

70. Graybiel, A.M. (2008). Habits, rituals, and the evaluative brain. Annu. Rev. Neurosci. 31, 359–387.

71. Graybiel, A.M., and Matsushima, A. (2023). Striosomes and Matrisomes: Scaffolds for Dynamic Coupling of Volition and Action. Annu. Rev. Neurosci. 46, 359–380.

72. Hong, S., and Hikosaka, O. (2008). The globus pallidus sends reward-related signals to the lateral habenula. Neuron 60, 720–729.

73. Hikosaka, O. (2010). The habenula: from stress evasion to value-based decision-making. Nat. Rev. Neurosci. 11, 503–513.

74. Matsumoto, M., and Hikosaka, O. (2007). Lateral habenula as a source of negative reward signals in dopamine neurons. Nature 447, 1111–1115.

75. Matsumoto, M., and Hikosaka, O. (2009). Two types of dopamine neuron distinctly convey positive and negative motivational signals. Nature 459, 837–841.

76. Steinberg, E.E., Gore, F., Heifets, B.D., Taylor, M.D., Norville, Z.C., Beier, K.T., Földy, C., Lerner, T.N., Luo, L., Deisseroth, K., et al. (2020). Amygdala-Midbrain Connections Modulate Appetitive and Aversive Learning. Neuron 106, 1026–1043.e9.

77. Menegas, W., Akiti, K., Amo, R., Uchida, N., and Watabe-Uchida, M. (2018). Dopamine neurons projecting to the posterior striatum reinforce avoidance of threatening stimuli. Nat. Neurosci. 21, 1421–1430.

78. Hong, S., Amemori, S., Chung, E., Gibson, D.J., Amemori, K.-I., and Graybiel, A.M. (2019). Predominant Striatal Input to the Lateral Habenula in Macaques Comes from Striosomes. Curr. Biol. 29, 51–61.e5.

79. Rajakumar, N., Elisevich, K., and Flumerfelt, B.A. (1993). Compartmental origin of the striato-entopeduncular projection in the rat. J. Comp. Neurol. 331, 286–296.

80. McGregor, M.M., McKinsey, G.L., Girasole, A.E., Bair-Marshall, C.J., Rubenstein, J.L.R., and Nelson, A.B. (2019). Functionally Distinct Connectivity of Developmentally Targeted Striosome Neurons. Cell Rep. 29, 1419–1428.e5.

81. Yin, H.H., Mulcare, S.P., Hilário, M.R.F., Clouse, E., Holloway, T., Davis, M.I., Hansson, A.C., Lovinger, D.M., and Costa, R.M. (2009). Dynamic reorganization of striatal circuits during the acquisition and consolidation of a skill. Nat. Neurosci. 12, 333–341.

82. Lerner, T.N., Shilyansky, C., Davidson, T.J., Evans, K.E., Beier, K.T., Zalocusky, K.A., Crow, A.K., Malenka, R.C., Luo, L., Tomer, R., et al. (2015). Intact-Brain Analyses Reveal Distinct Information Carried by SNc Dopamine Subcircuits. Cell 162, 635–647.

83. Lee, J., Wang, W., and Sabatini, B.L. (2020). Anatomically segregated basal ganglia pathways allow parallel behavioral modulation. Nat. Neurosci. 23, 1388–1398.

84. van Elzelingen, W., Warnaar, P., Matos, J., Bastet, W., Jonkman, R., Smulders, D., Goedhoop, J., Denys, D., Arbab, T., and Willuhn, I. (2022). Striatal dopamine signals are region specific and temporally stable across action-sequence habit formation. Curr. Biol. 32, 1163–1174.e6.

85. Mohebi, A., Wei, W., Pelattini, L., Kim, K., and Berke, J.D. (2024). Dopamine transients follow a striatal gradient of reward time horizons. Nat. Neurosci. 27, 737–746.

86. Ambrosi, P., and Lerner, T.N. (2022). Striatonigrostriatal circuit architecture for disinhibition of dopamine signaling. Cell Rep. 40, 111228.

87. Dhawale, A.K., Wolff, S.B.E., Ko, R., and Ölveczky, B.P. (2021). The basal ganglia control the detailed kinematics of learned motor skills. Nat. Neurosci. 24, 1256–1269.

88. Muñoz, B., Fritz, B.M., Yin, F., and Atwood, B.K. (2018). Alcohol exposure disrupts mu opioid receptor-mediated long-term depression at insular cortex inputs to dorsolateral striatum. Nat. Commun. 9, 1318.

89. Hilario, M., Holloway, T., Jin, X., and Costa, R.M. (2012). Different dorsal striatum circuits mediate action discrimination and action generalization. Eur. J. Neurosci. 35, 1105–1114.

90. Dexter, D.T., Carayon, A., Javoy-Agid, F., Agid, Y., Wells, F.R., Daniel, S.E., Lees, A.J., Jenner, P., and Marsden, C.D. (1991). Alterations in the levels of iron, ferritin and other trace metals in Parkinson’s disease and other neurodegenerative diseases affecting the basal ganglia. Brain 114 ( *Pt 4*), 1953–1975.

91. Marsden, C.D. (1982). Basal ganglia disease. Lancet 2, 1141–1147.

92. Gerfen, C.R., and Surmeier, D.J. (2011). Modulation of striatal projection systems by dopamine. Annu. Rev. Neurosci. 34, 441–466.

93. Howes, O.D., and Kapur, S. (2009). The dopamine hypothesis of schizophrenia: version III--the final common pathway. Schizophr. Bull. 35, 549–562.

94. Brisch, R., Saniotis, A., Wolf, R., Bielau, H., Bernstein, H.-G., Steiner, J., Bogerts, B., Braun, K., Jankowski, Z., Kumaratilake, J., et al. (2014). The role of dopamine in schizophrenia from a neurobiological and evolutionary perspective: old fashioned, but still in vogue. Front. Psychiatry 5, 47.

95. Menon, V., Palaniyappan, L., and Supekar, K. (2023). Integrative Brain Network and Salience Models of Psychopathology and Cognitive Dysfunction in Schizophrenia. Biol. Psychiatry 94, 108–120.

96. Kegeles, L.S., Abi-Dargham, A., Frankle, W.G., Gil, R., Cooper, T.B., Slifstein, M., Hwang, D.-R., Huang, Y., Haber, S.N., and Laruelle, M. (2010). Increased synaptic dopamine function in associative regions of the striatum in schizophrenia. Arch. Gen. Psychiatry 67, 231–239.

97. Abi-Dargham, A., Rodenhiser, J., Printz, D., Zea-Ponce, Y., Gil, R., Kegeles, L.S., Weiss, R., Cooper, T.B., Mann, J.J., Van Heertum, R.L., et al. (2000). Increased baseline occupancy of D2 receptors by dopamine in schizophrenia. Proc. Natl. Acad. Sci. U. S. A. 97, 8104–8109.

98. Piantadosi, S.C., Manning, E.E., Chamberlain, B.L., Hyde, J., LaPalombara, Z., Bannon, N.M., Pierson, J.L., K Namboodiri, V.M., and Ahmari, S.E. (2024). Hyperactivity of indirect pathway-projecting spiny projection neurons promotes compulsive behavior. Nat. Commun. 15, 4434.

99. Packard, M.G., and Knowlton, B.J. (2002). Learning and memory functions of the Basal Ganglia. Annu. Rev. Neurosci. 25, 563–593.

100. Schultz, W. (2016). Dopamine reward prediction-error signalling: a two-component response. Nat. Rev. Neurosci. 17, 183–195.

101. Liljeholm, M., and O’Doherty, J.P. (2012). Contributions of the striatum to learning, motivation, and performance: an associative account. Trends Cogn. Sci. 16, 467–475.

102. Crittenden, J.R., Yoshida, T., Venu, S., Mahar, A., and Graybiel, A.M. (2022). Cannabinoid Receptor 1 Is Required for Neurodevelopment of Striosome-Dendron Bouquets. eNeuro 9. 10.1523/ENEURO.0318-21.2022.

103. Gittis, A.H., and Kreitzer, A.C. (2012). Striatal microcircuitry and movement disorders. Trends Neurosci. 35, 557–564.

104. Assous, M., Kaminer, J., Shah, F., Garg, A., Koós, T., and Tepper, J.M. (2017). Differential processing of thalamic information via distinct striatal interneuron circuits. Nat. Commun. 8, 15860.

105. Matsushima, A., and Graybiel, A.M. (2022). Combinatorial Developmental Controls on Striatonigral Circuits. Cell Rep. 38, 110272.

106. Evans, R.C., Twedell, E.L., Zhu, M., Ascencio, J., Zhang, R., and Khaliq, Z.M. (2020). Functional Dissection of Basal Ganglia Inhibitory Inputs onto Substantia Nigra Dopaminergic Neurons. Cell Rep. 32, 108156.

107. Graybiel, A.M., Moratalla, R., and Robertson, H.A. (1990). Amphetamine and cocaine induce drug-specific activation of the c-fos gene in striosome-matrix compartments and limbic subdivisions of the striatum. Proc. Natl. Acad. Sci. U. S. A. 87, 6912–6916.

108. Moratalla, R., Vallejo, M., Elibol, B., and Graybiel, A.M. (1996). D1-class dopamine receptors influence cocaine-induced persistent expression of Fos-related proteins in striatum. Neuroreport 8, 1–5.

109. Canales, J.J., and Graybiel, A.M. (2000). Patterns of gene expression and behavior induced by chronic dopamine treatments. Ann. Neurol. 47, S53–S59.

110. Moratalla, R., Vickers, E.A., Robertson, H.A., Cochran, B.H., and Graybiel, A.M. (1993). Coordinate expression of c-fos and jun B is induced in the rat striatum by cocaine. J. Neurosci. 13, 423–433.

111. Wang, W., Xie, X., Zhuang, X., Huang, Y., Tan, T., Gangal, H., Huang, Z., Purvines, W., Wang, X., Stefanov, A., et al. (2023). Striatal μ-opioid receptor activation triggers direct-pathway GABAergic plasticity and induces negative affect. Cell Rep. 42, 112089.

112. Davis, M.I., Crittenden, J.R., Feng, A.Y., Kupferschmidt, D.A., Naydenov, A., Stella, N., Graybiel, A.M., and Lovinger, D.M. (2018). The cannabinoid-1 receptor is abundantly expressed in striatal striosomes and striosome-dendron bouquets of the substantia nigra. PLoS One 13, e0191436.

113. Seeman, P., and Lee, T. (1975). Antipsychotic drugs: direct correlation between clinical potency and presynaptic action on dopamine neurons. Science 188, 1217–1219.

114. Creese, I., Burt, D.R., and Snyder, S.H. (1996). Dopamine receptor binding predicts clinical and pharmacological potencies of antischizophrenic drugs. J. Neuropsychiatry Clin. Neurosci. 8, 223–226.

115. Howes, O., McCutcheon, R., and Stone, J. (2015). Glutamate and dopamine in schizophrenia: an update for the 21st century. J. Psychopharmacol. 29, 97–115.

116. McCutcheon, R.A., Abi-Dargham, A., and Howes, O.D. (2019). Schizophrenia, Dopamine and the Striatum: From Biology to Symptoms. Trends Neurosci. 42, 205–220.

117. Matsushima, A., Pineda, S.S., Crittenden, J.R., Lee, H., Galani, K., Mantero, J., Tombaugh, G., Kellis, M., Heiman, M., and Graybiel, A.M. (2023). Transcriptional vulnerabilities of striatal neurons in human and rodent models of Huntington’s disease. Nat. Commun. 14, 282.

118. Tippett, L.J., Waldvogel, H.J., Thomas, S.J., Hogg, V.M., van Roon-Mom, W., Synek, B.J., Graybiel, A.M., and Faull, R.L.M. (2007). Striosomes and mood dysfunction in Huntington’s disease. Brain 130, 206–221.

119. Hedreen, J.C., Berretta, S., and White, C.L., Iii (2024). Postmortem neuropathology in early Huntington disease. J. Neuropathol. Exp. Neurol. 83, 294–306.

120. Hikosaka, O., and Wurtz, R.H. (1985). Modification of saccadic eye movements by GABA-related substances. II. Effects of muscimol in monkey substantia nigra pars reticulata. J. Neurophysiol. 53, 292–308.

121. Keller, E.L. (1974). Participation of medial pontine reticular formation in eye movement generation in monkey. J. Neurophysiol. 37, 316–332.

122. Mink, J.W., and Thach, W.T. (1993). Basal ganglia intrinsic circuits and their role in behavior. Curr. Opin. Neurobiol. 3, 950–957.

123. Poewe, W. (2008). Non-motor symptoms in Parkinson’s disease. Eur. J. Neurol. 15 *Suppl 1*, 14–20.

124. Sharma, S., Moon, C.S., Khogali, A., Haidous, A., Chabenne, A., Ojo, C., Jelebinkov, M., Kurdi, Y., and Ebadi, M. (2013). Biomarkers in Parkinson’s disease (recent update). Neurochem. Int. 63, 201–229.

125. Julien, C.L., Thompson, J.C., Wild, S., Yardumian, P., Snowden, J.S., Turner, G., and Craufurd, D. (2007). Psychiatric disorders in preclinical Huntington’s disease. J. Neurol. Neurosurg. Psychiatry 78, 939–943.

126. Jog, M.S., Kubota, Y., Connolly, C.I., Hillegaart, V., and Graybiel, A.M. (1999). Building neural representations of habits. Science 286, 1745–1749.

127. Thorn, C.A., Atallah, H., Howe, M., and Graybiel, A.M. (2010). Differential dynamics of activity changes in dorsolateral and dorsomedial striatal loops during learning. Neuron 66, 781–795.

128. Desrochers, T.M., Jin, D.Z., Goodman, N.D., and Graybiel, A.M. (2010). Optimal habits can develop spontaneously through sensitivity to local cost. Proc. Natl. Acad. Sci. U. S. A. 107, 20512–20517.

129. Barnes, T.D., Mao, J.-B., Hu, D., Kubota, Y., Dreyer, A.A., Stamoulis, C., Brown, E.N., and Graybiel, A.M. (2011). Advance cueing produces enhanced action-boundary patterns of spike activity in the sensorimotor striatum. J. Neurophysiol. 105, 1861–1878.

130. Smith, K.S., Virkud, A., Deisseroth, K., and Graybiel, A.M. (2012). Reversible online control of habitual behavior by optogenetic perturbation of medial prefrontal cortex. Proc. Natl. Acad. Sci. U. S. A. 109, 18932–18937.

131. Smith, K.S., and Graybiel, A.M. (2013). A dual operator view of habitual behavior reflecting cortical and striatal dynamics. Neuron 79, 361–374.

132. Smith, K.S., and Graybiel, A.M. (2016). Habit formation coincides with shifts in reinforcement representations in the sensorimotor striatum. J. Neurophysiol. 115, 1487– 1498.

133. Fujii, N., and Graybiel, A.M. (2003). Representation of action sequence boundaries by macaque prefrontal cortical neurons. Science 301, 1246–1249.

134. Barnes, T.D., Kubota, Y., Hu, D., Jin, D.Z., and Graybiel, A.M. (2005). Activity of striatal neurons reflects dynamic encoding and recoding of procedural memories. Nature 437, 1158–1161.

135. Martiros, N., Burgess, A.A., and Graybiel, A.M. (2018). Inversely Active Striatal Projection Neurons and Interneurons Selectively Delimit Useful Behavioral Sequences. Curr. Biol. 28, 560–573.e5.

136. Jin, X., and Costa, R.M. (2010). Start/stop signals emerge in nigrostriatal circuits during sequence learning. Nature 466, 457–462.

137. Geddes, C.E., Li, H., and Jin, X. (2018). Optogenetic Editing Reveals the Hierarchical Organization of Learned Action Sequences. Cell 174, 32–43.e15.

138. Desrochers, T.M., Amemori, K.-I., and Graybiel, A.M. (2015). Habit Learning by Naive Macaques Is Marked by Response Sharpening of Striatal Neurons Representing the Cost and Outcome of Acquired Action Sequences. Neuron 87, 853–868.

139. Graybuck, L.T., Daigle, T.L., Sedeño-Cortés, A.E., Walker, M., Kalmbach, B., Lenz, G.H., Morin, E., Nguyen, T.N., Garren, E., Bendrick, J.L., et al. (2021). Enhancer viruses for combinatorial cell-subclass-specific labeling. Neuron 109, 1449–1464.e13.

140. Mich, J.K., Graybuck, L.T., Hess, E.E., Mahoney, J.T., Kojima, Y., Ding, Y., Somasundaram, S., Miller, J.A., Kalmbach, B.E., Radaelli, C., et al. (2021). Functional enhancer elements drive subclass-selective expression from mouse to primate neocortex. Cell Rep. 34, 108754.

141. Lee, H., Fenster, R.J., Pineda, S.S., Gibbs, W.S., Mohammadi, S., Davila-Velderrain, J., Garcia, F.J., Therrien, M., Novis, H.S., Gao, F., et al. (2020). Cell Type-Specific Transcriptomics Reveals that Mutant Huntingtin Leads to Mitochondrial RNA Release and Neuronal Innate Immune Activation. Neuron 107, 891–908.e8.

142. Berretta, S., Robertson, H.A., and Graybiel, A.M. (1992). Dopamine and glutamate agonists stimulate neuron-specific expression of Fos-like protein in the striatum. J. Neurophysiol. 68, 767–777.

143. Parthasarathy, H.B., and Graybiel, A.M. (1997). Cortically driven immediate-early gene expression reflects modular influence of sensorimotor cortex on identified striatal neurons in the squirrel monkey. J. Neurosci. 17, 2477–2491.

144. Manning, E.E., Dombrovski, A.Y., Torregrossa, M.M., and Ahmari, S.E. (2019). Impaired instrumental reversal learning is associated with increased medial prefrontal cortex activity in Sapap3 knockout mouse model of compulsive behavior. Neuropsychopharmacology 44, 1494–1504.

145. Märtin, A., Calvigioni, D., Tzortzi, O., Fuzik, J., Wärnberg, E., and Meletis, K. (2019). A Spatiomolecular Map of the Striatum. Cell Rep. 29, 4320–4333.e5.

146. Liu, K., Kim, J., Kim, D.W., Zhang, Y.S., Bao, H., Denaxa, M., Lim, S.-A., Kim, E., Liu, C., Wickersham, I.R., et al. (2017). Lhx6-positive GABA-releasing neurons of the zona incerta promote sleep. Nature 548, 582–587.

147. Weible AP, Schwarcz L, Wickersham IR, Deblander L, Wu H, Callaway EM, Seung HS, Kentros CG. Transgenic targeting of recombinant rabies virus reveals monosynaptic connectivity of specific neurons. J Neurosci. 2010 Dec 8;30(49):16509–13.

148. Wickersham, I.R., Sullivan, H.A., and Seung, H.S. (2010). Production of glycoprotein-deleted rabies viruses for monosynaptic tracing and high-level gene expression in neurons. Nat. Protoc. 5, 595–606.

149. Wickersham, I.R., and Sullivan, H.A. (2015). Rabies viral vectors for monosynaptic tracing and targeted transgene expression in neurons. Cold Spring Harb. Protoc. 2015, 375–385.

150. Chatterjee, S., Sullivan, H.A., MacLennan, B.J., Xu, R., Hou, Y., Lavin, T.K., Lea, N.E., Michalski, J.E., Babcock, K.R., Dietrich, S., et al. (2018). Nontoxic, double-deletion-mutant rabies viral vectors for retrograde targeting of projection neurons. Nat. Neurosci. 21, 638– 646.

151. Tai, L.-H., Lee, A.M., Benavidez, N., Bonci, A., and Wilbrecht, L. (2012). Transient stimulation of distinct subpopulations of striatal neurons mimics changes in action value. Nat. Neurosci. 15, 1281–1289.

152. Weglage, M., Wärnberg, E., Lazaridis, I., Calvigioni, D., Tzortzi, and O., Meletis, K. (2021) Complete representation of action space and value in all dorsal striatal pathways. Cell Rep. 36,109437.

153. He, M., Tucciarone, J., Lee, S., Nigro, M.J., Kim, Y., Levine, J.M., Kelly, S.M., Krugikov, I., Wu, P., Chen, Y., Gong, L., Hou, Y., Osten, P., Rudy, B., and Huang, Z.J. (2016) Strategies and tools for combinatorial targeting of GABAergic neurons in mouse cerebral cortex. Neuron. 91, 1228–1243.

154. Yoshida, J., Oñate, M., Khatami, L., Vera, J., Nadim, F., and Khodakhah, K. (2022) Cerebellar contributions to the basal ganglia influence motor coordination, reward processing, and movement vigor. J Neurosci. 42, 8406–8415.

155. Fullard, J.F., Hauberg, M.E., Bendl, J., Egervari, G., Cirnaru, M.D., Reach, S.M., Motl, J., Ehrlich, M.E., Hurd, Y.L., and Roussos, P. (2018) An atlas of chromatin accessibility in the adult human brain. Genome Res. 28, 1243–1252.

156. Hsu, A.I., and Yttri, E.A. (2021) B-SOiD, an open-source unsupervised algorithm for identification and fast prediction of behaviors. Nat. Commun. 12, 5188.

157. Mathis, A., Mamidanna, P., Cury, K.M., Abe, T., Murthy, V.N., Mathis, M.W., and Bethge, M. (2018) DeepLabCut: markerless pose estimation of user-defined body parts with deep learning. Nat. Neurosci. 21, 1281–1289.

158. Hueske, E., Stine, C., Yoshida, T., Crittenden, J.R., Gupta, A., Johnson, J.C., Achanta, A.S., Loftus, J., Mahar, A., Hu, D., Azocar, J., Gray, R.J., Bruchas, M.R., and Graybiel, A.M. (2024) Developmental and adult striatal patterning of nociceptin ligand marks striosomal population with direct dopamine projections. bioRxiv 2024 May 15:2024.05.15.594426.

159. Li, B., Piriz, J., Mirrione, M., Chung, C., Proulx, C.D., Schulz, D., Henn, F., and Malinow, R. (2011) Synaptic potentiation onto habenula neurons in the learned helplessness model of depression. Nature 470, 535–9.

160. Xiao, X., Deng, H., Furlan, A., Yang, T., Zhang, X., Hwang, G.R., Tucciarone, J., Wu, P., He, M., Palaniswamy, R., Ramakrishnan, C., Ritola, K., Hantman, A., Deisseroth, K., Osten, P., Huang, Z.J., and Li, B. (2020) A genetically defined compartmentalized striatal direct pathway for negative reinforcement. Cell 183, 211–227.e20.

161. Saunders, A., Macosko, E.Z., Wysoker, A., Goldman, M., Krienen, F.M., de Rivera, H., Bien, E., Baum, M., Bortolin, L., Wang, S., Goeva, A., Nemesh, J., Kamitaki, N., Brumbaugh, S., Kulp, D., and McCarroll, S.A. (2018) Molecular diversity and specializations among the cells of the adult mouse brain. Cell 174, 1015–1030.e16.

162. Shuen, J.A., Chen, M., Gloss, B., and Calakos, N. (2008) Drd1a-tdTomato BAC transgenic mice for simultaneous visualization of medium spiny neurons in the direct and indirect pathways of the basal ganglia. J Neurosci. 28, 2681–5.

